# PHoNUPS: Open-Source Software for Standardized Analysis and Visualization of Multi-Instrument Extracellular Vesicle Measurements

**DOI:** 10.64898/2026.01.29.702479

**Authors:** Bence Mélykúti, Gonzalo Bustos-Quevedo, Tony Prinz, Irina Nazarenko

## Abstract

Accurate and transparent characterization of extracellular vesicle (EV) preparations is essential to ensure reproducibility, comparability, and adherence to MISEV reporting standards. However, data outputs from commonly used instruments for assessing EV size, concentration, and surface charge (zeta potential) vary widely in format and structure, complicating standardized analysis and integration across platforms.

We present PHoNUPS (*Plotting the Histogram of Non-Uniform Particles’ Sizes*), free and open-source software (FOSS) developed in R, that enables unified processing, analysis, and visualization of EV characterization data. PHoNUPS computes statistics and generates standardized histograms and contour plots (for size against zeta potential) suitable for transparent reporting and cross-study comparison. The software produces high-quality, publication-ready figures. Third-party graphical editing tools allow users to refine and annotate visualizations for presentation or manuscript preparation. PHoNUPS supports multiple measurement file formats, thereby facilitating dataset integration from different instruments.

PHoNUPS was developed with extensibility at its core, providing a basis for user-driven growth. We invite the EV community—researchers, analysts, and tool developers—to use PHoNUPS, share feedback on their experience and needs, and contribute to the platform by integrating additional input data formats, analytical routines, and visualization functionalities.

**Graphical abstract:** The free software PHoNUPS processes the outputs of several different EV characterization instruments and it is extensible with further ones. It computes statistics of particle size and zeta potential distributions and it plots the corresponding histograms or contour plots.

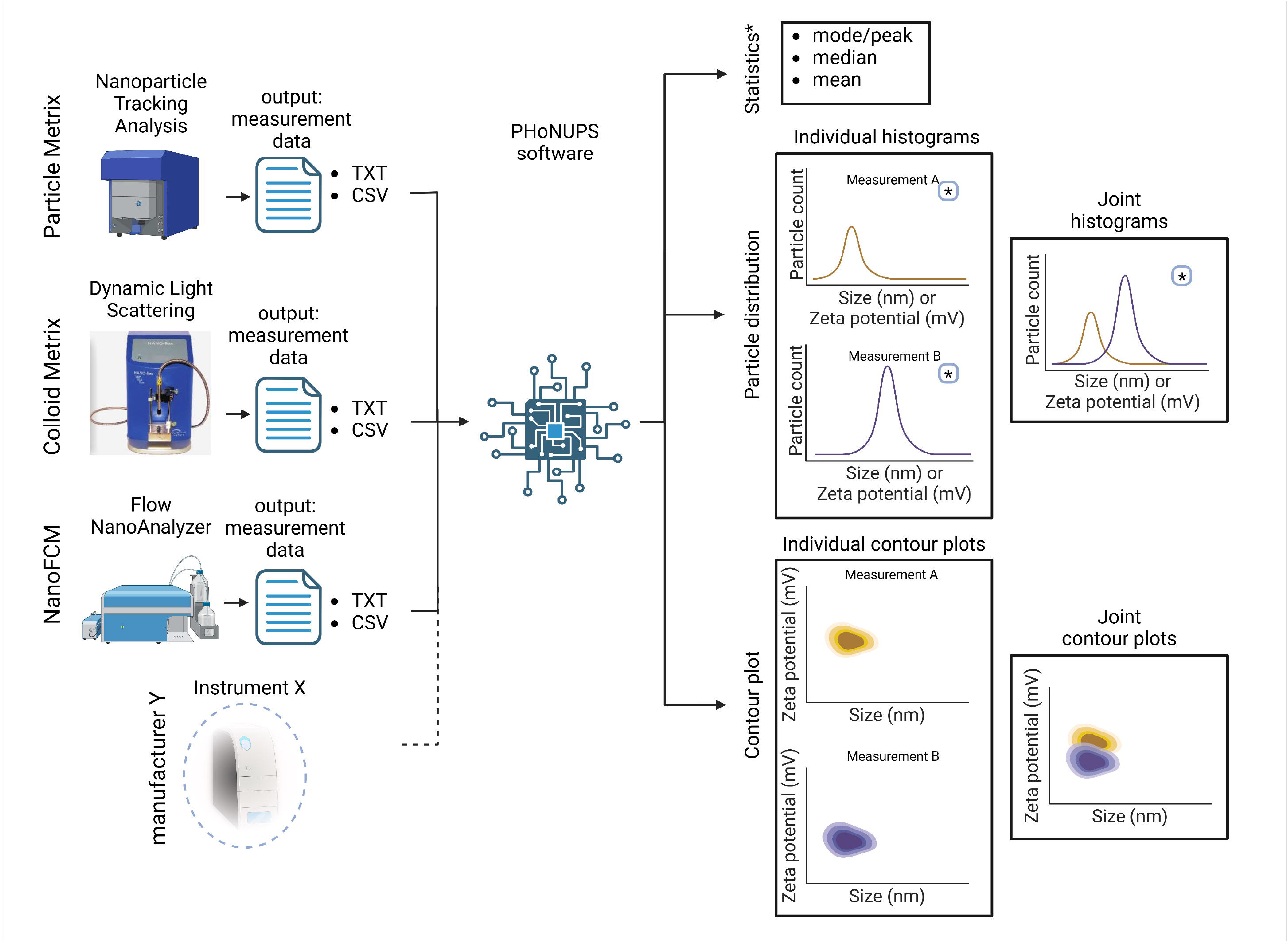

## Introduction

### The Rationale for Characterizing EV Size, Concentration, and Surface Charge

Accurate quantification of particle size and concentration is fundamental for reproducible EV research and is strongly emphasized in MISEV (Welsh et al., 2024, Section 5, EV characterization) and associated reference documents. Commonly used methods include *flow cytometry, nano-flow cytometry, nanoparticle tracking analysis* (NTA), also referred as a *single particle tracking, resistive pulse sensing* (RPS), *multi-angle light scattering*, and *dynamic light scattering* (DLS), also referred to as *photon correlation spectroscopy* (PCS) and *quasi-elastic light scattering* (QELS) (Welsh et al., 2024, Section 6). These technologies yield two essential datasets: (i) particle concentration and (ii) particle diameter distributions. Size distributions may be summarized via descriptive statistics such as the arithmetic mean, median, standard deviation, mode (the particle size with the greatest count of observations), minimum, and maximum. However, reducing EV populations to a single value or a set of summary statistics is insufficient. We emphasize the importance of visualizing and reporting the full empirical size distribution, rather than relying solely on mean or median values, as histograms uncover sample and preparation heterogeneity and improve transparency and comparability across datasets.

Beyond size, surface charge (zeta potential) is a critical yet underreported quantitative parameter of EVs. Zeta potential influences particle behavior in suspension, colloidal stability, structural integrity, biocorona formation, and interactions with cells and other particles (Németh et al., 2022; Shaw et al., 2025). In physiological systems, the surface charge of particles at neutral pH is often negative due to functional groups (e.g. carboxyl, phosphate, amine) that deprotonate, inducing a net negative surface charge. This contributes to a negative zeta potential, which is critical for colloidal stability and interactions in physiological environments (Ow et al., 2025). The surface charge affects interactions of particles with immune cells and can modulate immune system activation (Freitag et al., 2020; Lee, Choi, Webster, Kim, & Khang, 2015). Reported EV zeta potential values range between −60 mV and −10 mV in phosphate-buffered saline (PBS), with or without Tween 20, and depend on buffer composition and dilution conditions (Midekessa et al., 2020; Nguyen, Tran, Kaestner, & Bernhardt, 2022). Depending on the source of the EVs, different subpopulations of particles could potentially be distinguished by the zeta potential. In most EV isolation techniques, the co-isolation with unwanted particles like lipoproteins is unavoidable (Chou et al., 2024). Still, we anticipate that zeta potential measurements may help distinguish EV subpopulations from lipoprotein contaminants and warrant more systematic reporting in the field.

### Current Challenges in EV Data Analysis, Visualization and Existing Software Solutions

Despite the importance of EV size and surface charge for biological interpretation and quality control, data visualization and reporting practices lack standardization. Instrument control software typically generates human-readable PDF reports containing histograms of particle size; however, these plots cannot be readily exported, reformatted, or integrated into other reports. The protocol (metadata) of the measurement and the raw numerical data are often stored in one or more separate files, complicating data processing. As a result, EV size and zeta potential data are frequently published with the help of graphing software such as *Prism* (GraphPad Software, Boston, MA, USA) as separate plots containing particle size mode, particle size median and particle concentration comparisons (Ho et al., 2025; Mladenovic, Brealey, Peacock, Koort, & Zarovni, 2025), or they are copied directly as screenshots from the PDF reports (Huang et al., 2024; Liu et al., 2025; Winter et al., 2025; Zhou et al., 2024). They can be plotted manually using spreadsheet software (Ho et al., 2025), but this is time-consuming, prone to error, hardly reproducible, and produces inconsistent visual formats when data originate in different instruments that use differing histogram bin centers and bin sizes. These limitations impede cross-study comparison and standardization, both within and between laboratories.

Only a limited number of tools exist to address this challenge. Examples include *Floreada*.*io*, a free web-based, but proprietary application for plotting size distributions and calculating statistics from Flow Cytometry Standard (FCS) files (“Floreada.io,” 2020-2024; Mladenovic et al., 2025). To enhance quantitative measurements, specialized software for image-based NTA frame re-analysis has been developed to improve size estimation (Defante, Vreeland, Benkstein, & Ripple, 2018). *MPTHub* (Multiple Particle Tracking Analysis) is an open-source Python application to analyze NTA particle trajectories (Gabriel, Almeida, Avelar, Sarmento, & das Neves, 2022). Additional efforts exist for DLS-specific analysis (Salazar, Srivastav, Srivastava, & Srivastava, 2023), using the TurboCorr digital correlator, which is part of the NanoBrook DLS instrument (Brookhaven Instrument, Holtsville, NY, USA). We are unaware of a comprehensive solution for standardized visualization across EV measurement platforms, particularly for zeta potential data.

### Motivation for Developing PHoNUPS

To unlock the full potential of high-throughput EV characterization instruments, output data must be amenable to automated, reproducible, and standardized software analysis. Until instrument manufacturers adopt an interoperable data format, the community requires a tool that is capable of parsing heterogeneous output formats into a unified intermediary data structure, which is suitable for statistical analysis and visualization. A standard data file format in this field could be the FCS, which is maintained by the International Society for Advancement of Cytometry (ISAC) (Spidlen et al., 2010). This is the format that the Flow NanoAnalyzer (NanoFCM Inc., Xiamen, China) already uses.

To address the unmet need, we developed PHoNUPS (Mélykúti, 2025), a free and open-source R-based software for standardized processing and visualization of EV size and zeta potential distributions. In this first release, PHoNUPS provides an extensive coverage of software versions and their outputs for the NTA instrument ZetaView x20 Series by Particle Metrix GmbH (Inning am Ammersee, Germany), for the DLS instrument NANO-flex 180° DLS Size by Colloid Metrix GmbH (Meerbusch, Germany), and the single-EV flow cytometry instrument Flow NanoAnalyzer by NanoFCM Inc. (Xiamen, China). PHoNUPS computes descriptive statistics, plots size and zeta potential histograms, and generates two-dimensional (2D) contour plots (size versus zeta potential), which can be displayed on screen or exported in a consistent format to a publication-ready file. Expansion to additional instruments and data types is envisioned in close collaboration with the community and will be guided by user feedback, needs, and requests.

## PHoNUPS: Target Users, Core Functionality, Visualization Features

### Design Goals and Target Users

PHoNUPS was developed to enable fast and intuitive visualization of EV size and zeta potential measurements through histograms or contour plots for users with no programming experience with minimal user input. While the initial focus is on EVs, the software is applicable to any sub-micrometer particles measurable by NTA, DLS, or single-EV flow cytometry. Our aim is to provide a simple and seamless user experience, supported by efficient local execution and good engineering practices to ensure fast processing.

A core design principle is extensibility: PHoNUPS is intended to evolve with user needs, new instruments and instrument output formats. Because software updates from manufacturers frequently alter data formats, we wish to offer a framework that supports multiple versions of instrument control software and that facilitates the addition of further instrument output formats. We envision this process to occur in close interaction with the community, with new parsers written based on user feedback or contributed by the community and shared openly back into PHoNUPS.

### Core Functionalities

The user selects one or multiple measurement files as input. PHoNUPS computes descriptive statistics for each sample separately (mean, median, mode, minimum and maximum observed particle size or zeta potential).

For histogram-based visualization (either size or zeta potential only), PHoNUPS generates two complementary plot types:

1. Individual histogram panels: Each measurement is displayed in its own subplot. Each sample is annotated with its mean, median and mode both numerically and graphically. (These annotations can be turned on or off.)
2. Joint histogram plot: All selected measurements are overlaid in a single plot, each shown as a differently colored line histogram, enabling direct comparison between samples.

If the input files contain particle-by-particle size and zeta potential data, then PHoNUPS switches to a 2D visualization mode and shows size against zeta potential. To avoid overplotting of large datasets (an incomprehensibly large number of data markers), a 2D kernel density estimation (KDE) is applied, which turns the set of points into a density function. The density function is reduced to its level sets and results are visualized as contour plots instead of scatter plots.

As with histograms, PHoNUPS provides two plot types:

1. Individual contour plot panels: Each measurement gives one contour plot in a separate subplot.
2. Joint contour plot: A single chart contains all datasets, each represented by a differently colored contour plot.

For these 2D visualizations, users can choose between contour line maps or heatmap-style, filled contour band plots.

### Processing of Replicates and Data Handling

In typical workflows, technical replicates are acquired for each biological sample. Therefore PHoNUPS implements a standardized procedure for handling replicate measurements.

In basic histogram mode, replicates are averaged on a per bin basis and output as if they were a single input measurement. We explain what we mean by averaging (Figure 3A). Our instruments output binned particle size measurements, that is, the count of particles in bins of given, fixed particle sizes (respectively, zeta potentials). The sizes (resp., zeta potentials) are no longer available as continuous values but are discretized to the bin sizes. The averaging step means that we take the arithmetic mean of the particle counts across the replicates for every bin separately. All summary statistics are then derived from this aggregated distribution. This procedure is illustrated with examples in the section *Demonstration*.

For 2D contour plots, replicate datasets are concatenated, that is, the particles measured in all repeated measurements are taken together as a single union set.

The implications of this approach for users and potential pitfalls are detailed in the section *User Interaction*.

PHoNUPS supports three types of input data dimension on the y-axis of histograms for different input modalities:

- particle concentration (particle count/ml) per bin,
- particle count per bin,
- relative frequency per bin.

The x-axis uses nanometers (nm) for size and millivolts (mV) for zeta potential.

### Supported Inputs and Automatic Format Detection

In this first release, the user can select input files from different instruments and their respective, potentially different software versions, and PHoNUPS automatically detects the source format and applies the appropriate parsing routine. Heterogeneous data inputs (e.g. from different instruments or control software versions) are processed and plotted in a unified manner, combined in the joint plot, as long as they share the same data dimension (either size only or zeta potential only or particle-by-particle size and zeta potential measurements). Mixing datasets of different dimensionality is not supported.

Automatic input recognition is possible because the parsed formats are sufficiently distinct to allow the unique identification of the instrument and software version. With future harmonization of instrument output data across the field, this feature may become more challenging to maintain, and users may eventually need to specify the source of input files.

### Output Formats and Post-Processing Options

Plots may be displayed on-screen or exported in any format supported by ggsave() of the R package ggplot2: .eps, .ps, .tex, .pdf, .jpg, .tiff, .png, .bmp, .svg, .wmf. The vector graphics format .svg and the raster image format .png have been successfully tested.

The usage of PHoNUPS opens the opportunity to treat the measurement and the resulting histogram or contour plot data from different instruments in a unified manner. The use of vector graphics enables downstream batch processing and post-editing in graphical editors, allowing users to uniformly edit labels, adjust layout, fonts, colors, line thickness, or add area shading. This provides a level of flexibility that vendor-generated PDF reports lack.

### Extensibility and Future Directions

Our vision for PHoNUPS is to support high-throughput processing, large-scale dataset handling, and increasingly automated analysis workflows. To accommodate users with different levels of computational expertise, PHoNUPS will be required to develop along two complementary calling methods:

1. Interactive Mode: Designed for users with no programming experience, providing a simple, guided interface for visualizing datasets and exporting publication-ready plots.
2. Programmatic Mode (as a package): Aimed at experienced users and bioinformaticians, enabling scripted execution, batch processing, and integration into larger data-analysis pipelines. In this mode, users would be able to call PHoNUPS functions directly as an R package. The user would input a list of files in a structured (tabular or dictionary) format. At a minimum, the data structure would contain the input file paths and their corresponding output file names.

Future extensions, including support for additional instruments, data types, and visualization modes, will be developed in collaboration with the EV community and will be guided by user feedback and community contributions.

## Software Architecture

PHoNUPS is written in R (R Core Team, 2025), which is free, open-source, copyleft, easily accessible, and for this purpose fast software that integrates both basics and complex algorithms. It is a platform used worldwide.

Data reading and parsing are clearly separated from the statistical processing and visualization of the same data. Creating such a data interface ensures that new input data formats can be added with relative ease. See Figure 1 for a diagram of the software architecture.

**Figure 1.**
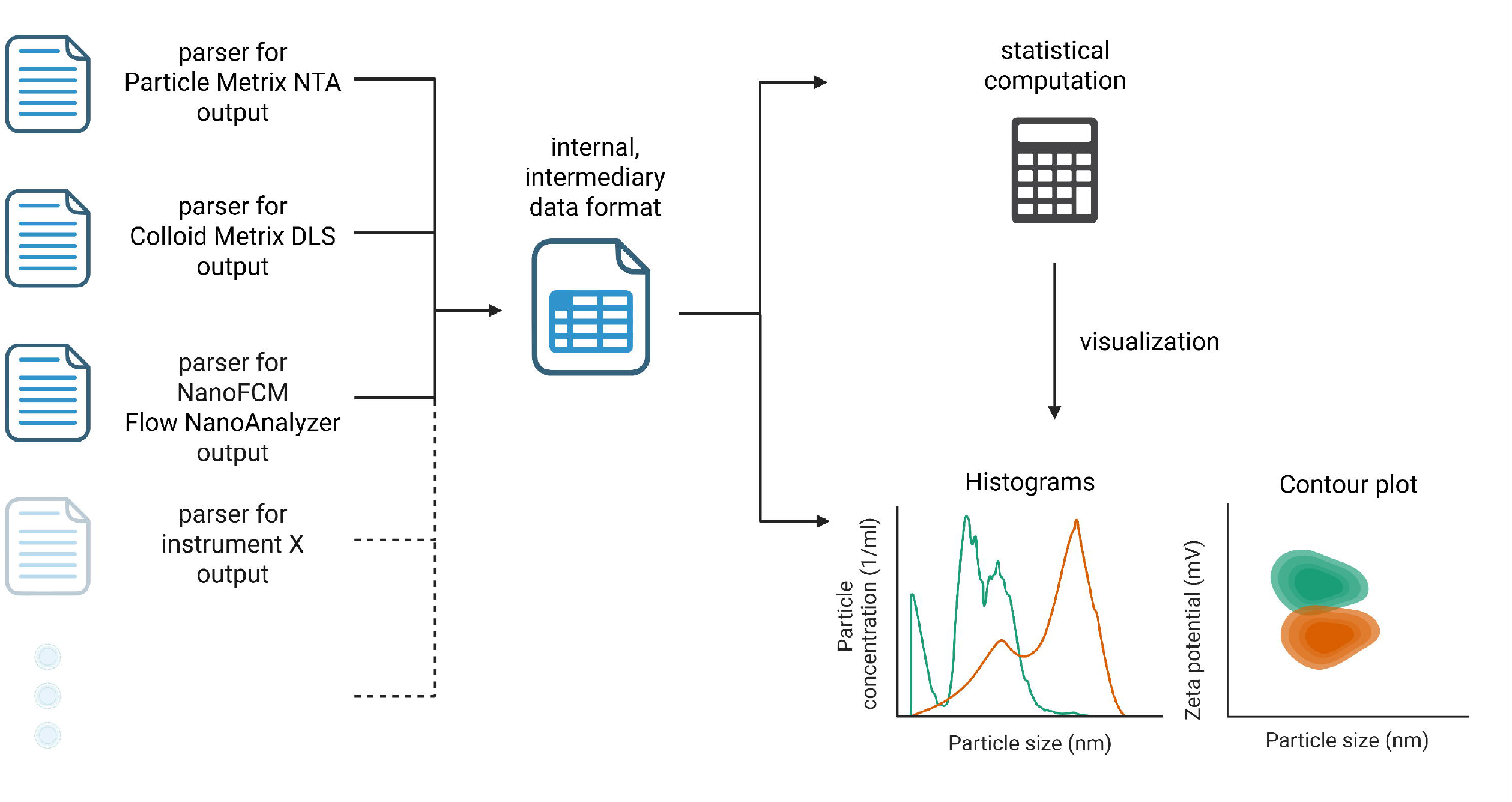
PHoNUPS software architecture. Using one of a number of measurement data parsers, PHoNUPS converts input data into a standard, internal, intermediary data format. The release version contains parsers for the instruments Nanoparticle Tracking Analysis from Particle Metrix, Dynamic Light Scattering from Colloid Metrix, and Flow NanoAnalyzer from NanoFCM. New parsers can easily be added. The standardized internal data is statistically processed and visualized such that the figures contain descriptive statistics from the statistical computation.

## User Interaction

### Installation and Setup

PHoNUPS requires the R environment, which is free and open-source software (FOSS) for statistical computing.

We distribute the software and associated sample data via a public repository (https://gitlab.com/intoev/phonups/phonups), from where users need to download PHoNUPS. Prior to first use, users must install a few R packages. Some parameter settings can be customized in the *user_settings*.*R* file.

To run PHoNUPS, the user sets in the R console the working directory to the PHoNUPS folder and executes the main script.

### Step-by-Step User Workflow

The step-by-step user interaction workflow is summarized in the flowchart in Figure 2.

**Figure 2.**
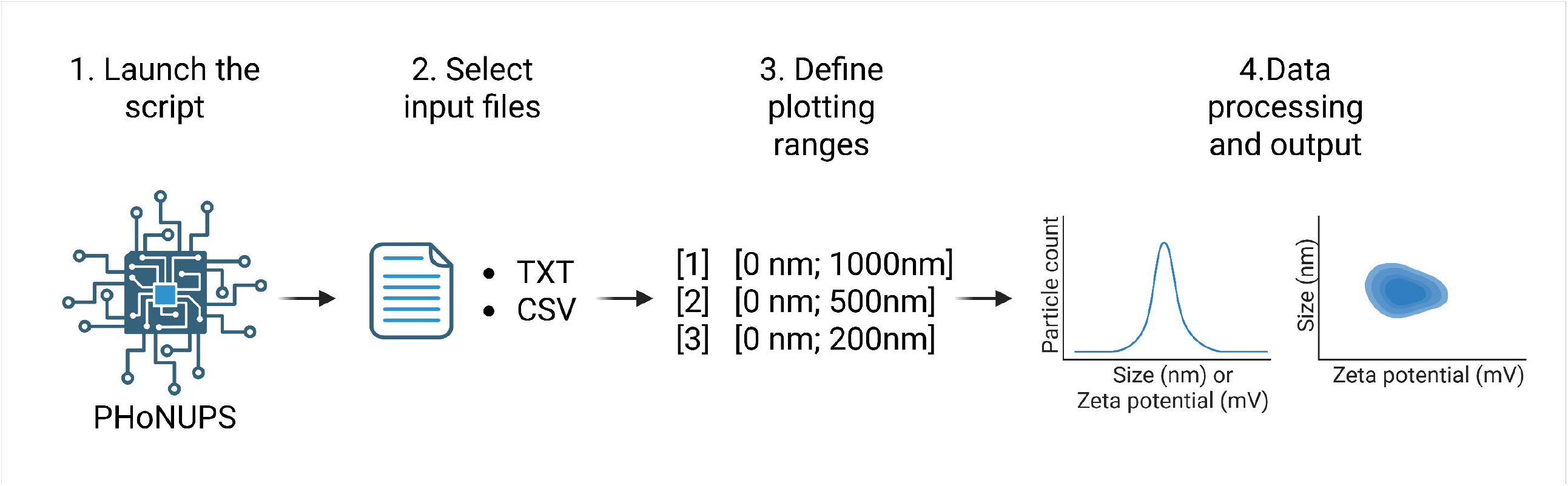
Schema of the steps to use PHoNUPS. First, the PHoNUPS script is started. Then the data input files are selected, and the plotting range is defined. Then, PHoNUPS processes the data, computes and displays descriptive statistics, and plots histograms or contour plots. The output plots can be generated as svg files, to be able to modify e.g. font size, color or transparency.

**Figure 3.**
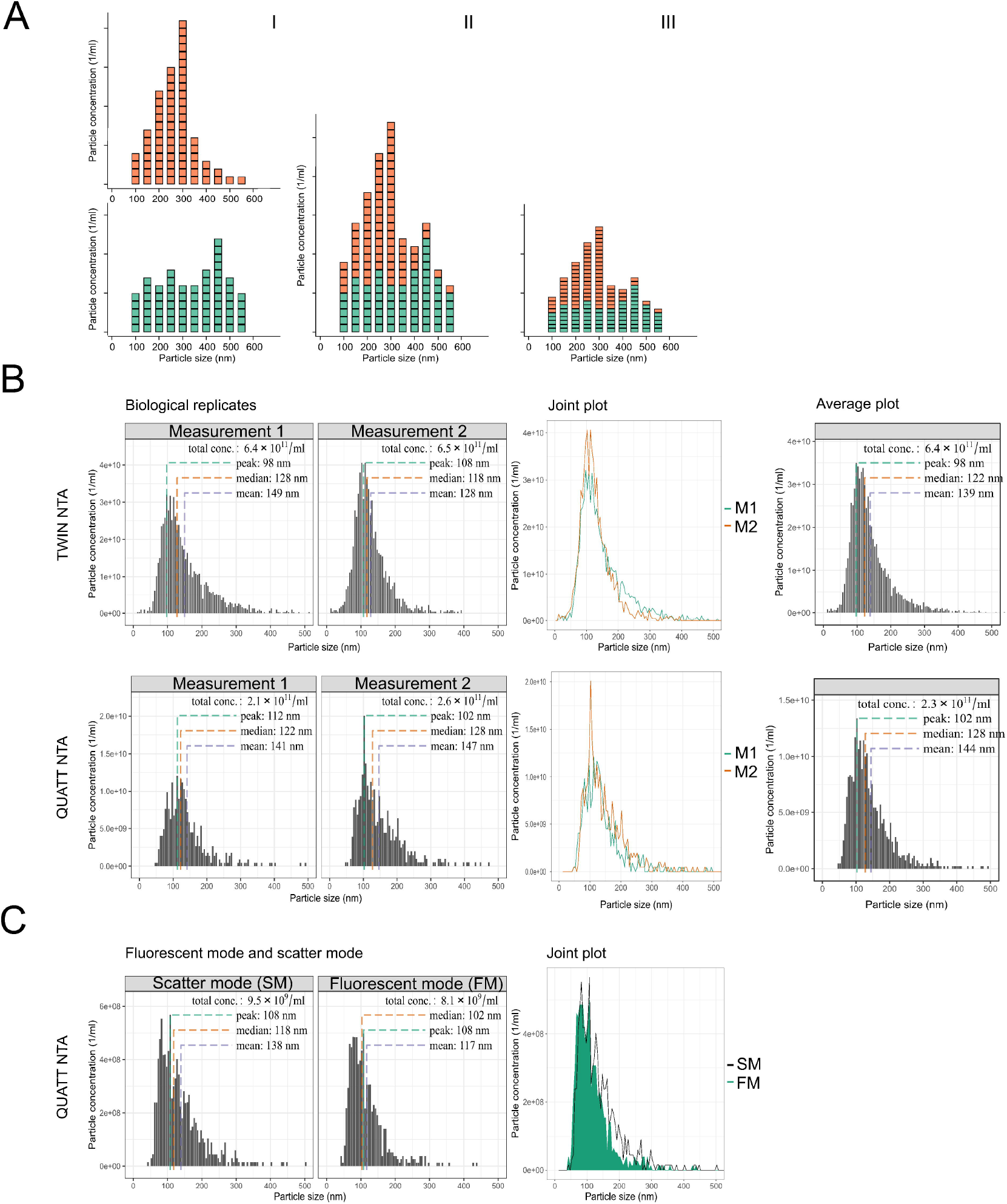
We demonstrate how PHoNUPS treats replicates by averaging and how it displays data for TWIN and QUATT NTA. A) Visualization of the averaging on the example of two technical replicates. I) The two measurements give individual distributions, which we display with different colors. II) We take the sum of the counts or concentrations in each bin separately. III) We divide the sums with the number of replicates that were summed, in this case with 2. Note that the mode in the average is not infrequently the mode of one of the technical replicates, like in this illustration. B) Representation of biological replicates of cell culture-derived EVs measured on TWIN (first row) and QUATT (second row) NTA. The first column shows the replicates as individual measurements, the middle column shows the joint plots of both measurements, and the last column displays the average of the two replicates. C) Representation of fluorescent EVs measured on the QUATT NTA in scatter and fluorescent modes. The first column shows both independent measurements. The second column displays the joint chart, showing the size distribution of the fluorescent particles with the size distribution of the particles measured in scatter mode. The partial concentrations of the fluorescent measurement are typically below those of the scatter mode measurement. The joint plot helps to visually confirm that the detected fluorescent particles correspond to the same population of particles as the one measured in scatter mode.

#### 1. Launch the PHoNUPS script

#### 2. Select input files

After launching the script, a file selector dialog box opens. The user selects one or multiple input files, with each file representing one measurement. Files may originate from different instrument models and software versions. It is in the nature of the file dialog that all files to be processed in a single PHoNUPS run must be stored in the same directory.

#### 3. Define plotting ranges

Histograms: Histograms are plotted with a shared x-axis (particle size or zeta potential) data range. In view of this, the user selects the shared x-axis range from a predefined list. In the default options for particle size, the left endpoint is always 0 nm; for zeta potential, the default intervals are symmetric around 0 mV. These default endpoints can be modified in the settings file.

Contour Plots: When particle-by-particle size and zeta potential measurements are available, users select shared axis ranges for both variables. Users may also specify in the settings file which of size and zeta potential should be on the x-axis and which on the y-axis.

#### 4. Data Processing and Output

PHoNUPS computes descriptive statistics for each sample and prints them to the console. The software then generates the individual plots and the joint plot according to the implied visualization mode (histograms or contour plots) and output format (display or save).

### Visualization Outputs

Via the user settings file, users can choose whether plots are displayed on screen or saved to files. For the display of plots, instead of the standard R interpreter, we recommend RStudio (Posit Software, PBC; Boston, MA, USA), the integrated development environment, where plots do not need to be closed in order to progress or to call the program again.

Output files are saved in the same directory where the input files are located. (Note that this is a unique directory.) The two output file names begin with the run date and time (in *YYYY-MM-DD_hh-mm-ss* format), which ensures that new runs do not overwrite existing output files.

Depending on the input, PHoNUPS visualization is in one of two modes:

- Histograms (size or zeta potential),
- 2D contour plots (size vs. zeta potential), which can be displayed as contour line maps or as heatmap-style, solid contour band plots.

In both of these two modes, in each run, PHoNUPS generates two complementary plot types:

- Individual plots: one panel per measurement,
- Joint plot: a single chart containing all measurements for direct comparison.

The panel labels in the individual plots, respectively, the dataset labels in the joint plot are the input file names without extension.

### Handling of Technical Replicates

We said that for histograms, technical replicate measurement files are averaged per bin, and for contour plots, the union of the particles in the technical replicate measurements is taken. In both cases, a single distribution is produced from which statistics are computed and plots are created.

In order to keep the user interaction simple, the user tells PHoNUPS via the file naming which files are replicates. In turn, PHoNUPS automatically detects technical replicate measurements based on file name patterns.

Typical file names could be *sample_1-1*.*txt, sample_1-2*.*txt, sample_1-3*.*txt* for the three replicates of sample 1, and *sample_2-1*.*txt, sample_2-2*.*txt, sample_2-3*.*txt* for the three replicates of sample 2. For this reason, PHoNUPS detects *-\*\*\**. (hyphen, one or multiple digits, period) in a file name as the sign for replicates, removes *-\*\*\** (hyphen, all digits until the period) from the name, and input files which now have the same name are averaged (or in the case of contour plots, merged) together and treated as one sample with this shortened name.

In summary, any file name containing a hyphen followed by one or more digits directly before the file extension (e.g. *sample_1-1*.*txt, sample_1-2*.*txt*) is interpreted as one of a number of technical replicates.

The shortened file name (with the replicate suffix removed) is used as the sample name (panel title) in the individual plot and as the dataset label in the joint plot.

Important note: This feature may unintentionally merge files with naming patterns that resemble replicates. It interferes with naming conventions that prepend the sample number with a hyphen (*mysample-1*.*txt, mysample-2*.*txt* would be treated as technical replicates of *mysample*) or contain the date (the file name *sample_2026-01-01*.*txt* would be truncated). This potential pitfall is reiterated in the documentation.

The discussed potential future programmatic calling method will need to enable the prescription of the replicate relationship between input files.

R determines the order of the input files in the output plots’ panels, respectively, legend internally by their alphabetical order. Hence users can prescribe the plotting order of their samples via prefixing file names alphabetically or numerically (e.g. *1_*, 2_*, 3_\**, … or *a_*, b_*, c_\**, …).

We recommend using the vector graphics format .svg because this enables the editing of the plots postproduction, for which we use the FOSS *Inkscape*. With its help, the user can remove the leading characters from subplot titles, respectively, the legend.

### Supported Operating Systems

PHoNUPS currently runs under Windows and Linux.

## Demonstration

To illustrate the functionalities of PHoNUPS, we applied the software to a set of representative EV datasets obtained using commonly used characterization platforms. The examples demonstrate how PHoNUPS standardizes data visualization across instruments, measurement modes, and sample types. The datasets include EVs derived from cell culture supernatants and from human plasma processed by size-exclusion chromatography (SEC) or tangential flow filtration (TFF). Measurements were acquired using Particle Metrix ZetaView x20 Series PMX-220 instruments (TWIN and QUATT configurations, Particle Metrix GmbH, Inning am Ammersee, Germany), the NANO-flex 180° DLS Size instrument (Colloid Metrix GmbH, Meerbusch, Germany), and the Flow NanoAnalyzer (NanoFCM Inc., Xiamen, China), and include both size and, for the ZetaView instruments, zeta potential outputs.

### Example 1. Histogram Visualization and Replicate Handling

In this example, we demonstrate how PHoNUPS visualizes particle size distributions and handles replicate datasets in histogram mode. To illustrate this, EVs isolated from cell culture supernatants were measured with the TWIN and QUATT NTA instruments. PHoNUPS displays each measurement first as an individual histogram, second, it enables users to overlay multiple measurements for direct comparison. This allows users to assess reproducibility and variability between replicates or measurement modes.

Figure 3A and B illustrate the default processing of technical replicates. First, individual histograms of two replicates are shown separately. Then, in Figure 3A, the two measurements are summed, and finally divided by the number of measurements, which is 2. In Figure 3B, EV isolated from the cell culture supernatants in biological replicates were measured with TWIN and QUATT NTA, and analyzed in the same manner, allowing users to compare their distributions individually, then as overlays, and finally to see the average of the replicates in a single representative histogram, from which the summary statistics are derived. In the first two columns, we withheld the information from PHoNUPS by appropriate file naming that these were replicates. Figure 3C demonstrates the comparison of measurements obtained in scatter and fluorescent modes on the QUATT instrument. PHoNUPS renders these data in a consistent format, facilitating the evaluation of mode-specific differences.

In summary, PHoNUPS provides clear visualizations of individual and combined histograms and consolidates technical replicates into a single representative distribution, simplifying interpretation and reporting. In the following, we will display histograms without guidelines for mode/peak, median and mean by changing a user setting.

### Example 2. Multi-Instrument Visualization of Size Distributions Using DLS, NTA, and NanoFCM

In this example, we demonstrate how PHoNUPS processes outputs from different measurement platforms into a unified visualization format, how it enables focused inspection of particle subpopulations using the x-axis zoom function, and that it provides compatibility with Flow NanoAnalyzer data.

EVs derived from cell culture supernatants were isolated by TFF followed by SEC, and then Fraction 1 was analyzed using DLS, two NTA instruments, and NanoFCM. PHoNUPS formats its output from disparate instruments consistently, allowing direct comparison of size distributions despite the differing data structures.

Figure 4A shows that the DLS instrument reports relative particle frequencies in the bins, whereas the NTA outputs absolute particle concentrations. The DLS measures larger particle sizes with larger bins, therefore its output has less precision. (We do not claim that the measurements are less accurate.) Another consequence is that the bin centers of the DLS are not equally spaced but follow a geometric progression with a factor of approximately 1.189, which results in a logarithmic size scale. PHoNUPS handles these differences. It preserves the native units while aligning axis scales and graphical formatting, although the bin centers differ between the DLS and the NTA instruments. All instruments revealed a dominant particle population centered around 120–140 nm, and the results obtained by different NTA generations are comparable.

**Figure 4.**
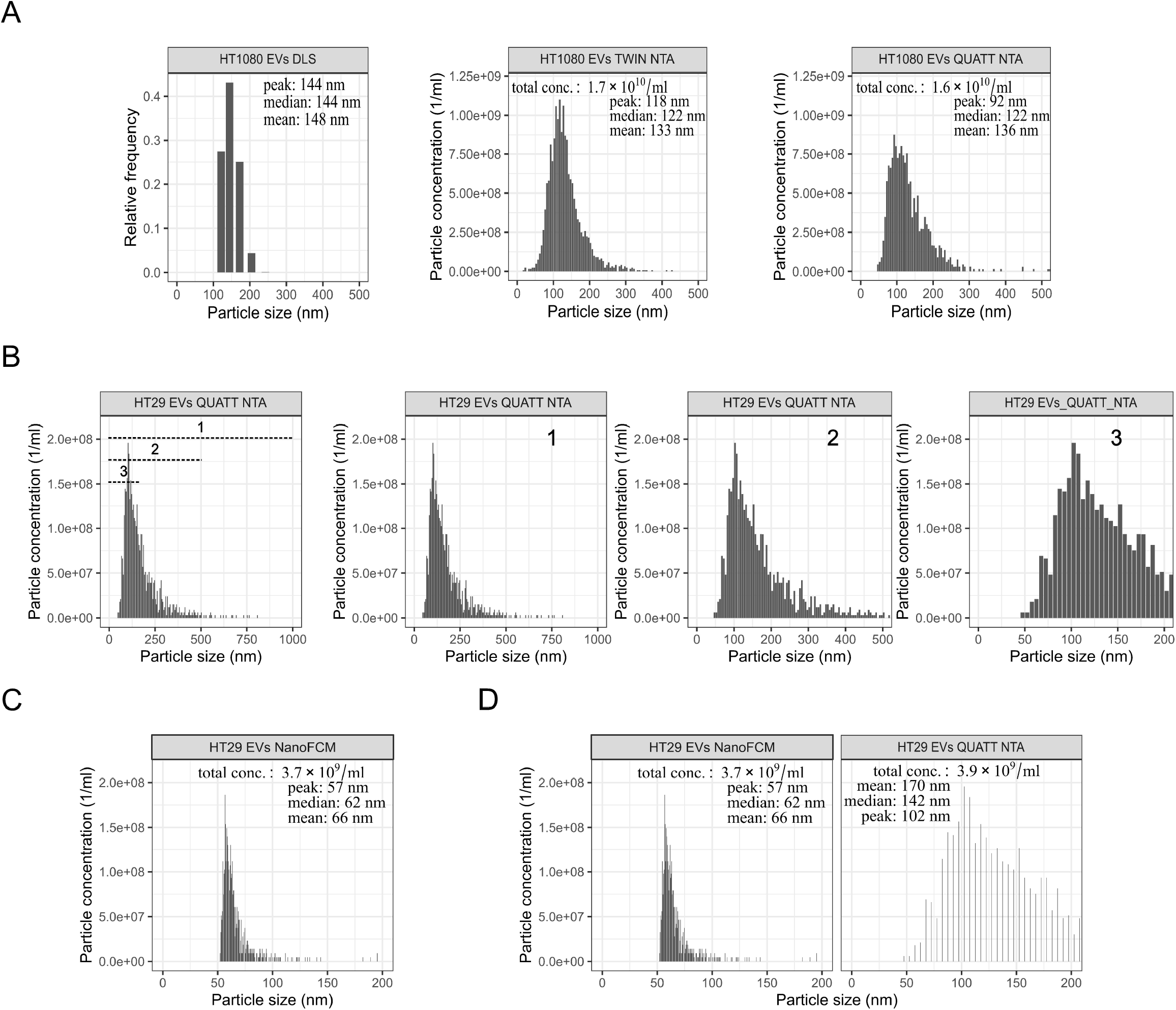
PHoNUPS displays data for comparison between DLS, TWIN and QUATT NTA. A) Representation of three different measurements performed by DLS, TWIN and QUATT NTA, respectively, of the same sample, Fraction 1 of HT1080 cell culture-derived EVs. B) PHoNUPS displays data with different endpoints on the x-axis. Representation of HT29 cell culture-derived EVs after SEC, measured by NTA. To respond to the different subpopulations that may appear in a sample, the user is able to adjust in PHoNUPS the range of displayed sizes. In this case, we opted for having three different x-axis endpoints of the same sample, which helps to provide a better visualization of the subpopulation of interest. By default, PHoNUPS plots in the ranges [0 nm, 1000 nm], [0 nm, 500 nm] and [0 nm, 200 nm]. This choice affects only the visualization; the statistics do not change. C) PHoNUPS displays the size distribution of the latter sample measured with Flow NanoAnalyzer. D) PHoNUPS displays a comparison between measurements with Flow NanoAnalyzer and NTA, having the range [0 nm, 200 nm].

We next illustrate how PHoNUPS can selectively zoom into specific size ranges to highlight features that may be less apparent at broader scales. Using the same NTA dataset, Figure 4B shows the application of the three preset x-axis ranges (0–1000 nm, 0–500 nm, 0–200 nm). Narrowing the axis range makes subtle population characteristics more visible without altering the underlying data. This feature enables users to tailor visualizations depending on the degree of sample heterogeneity or the biological question of interest. The number and the concrete set of available x-axis ranges are adjustable user settings.

Finally, PHoNUPS supports the importing and visualization of histogram exports from the Flow NanoAnalyzer control software. Figure 4C demonstrates how NanoFCM data is parsed and displayed using the same standardized layout applied to NTA and DLS. In Figure 4D, the comparison of NanoFCM and NTA profiles for the same sample shows differences in size distribution and summary metrics, reflecting intrinsic differences in measurement principles.

In conclusion, PHoNUPS can process a mix of heterogeneous input formats in a single run and display each dataset according to its dimension (in the example relative frequency and absolute concentration). It harmonizes size-distribution visualizations across platforms without altering native quantitative information. Adjustable x-axis limits enable focused inspection of particle subpopulations. Compatibility with NanoFCM extends the software’s applicability across commonly used EV characterization instruments.

### Example 3. Assessing Particle Heterogeneity in Plasma-Derived SEC Fractions

In this example, we demonstrate how PHoNUPS supports the visualization of heterogeneous particle populations across SEC fractions of plasma-derived samples, and how combining different analytical platforms provides a more comprehensive overview of the particle size landscape. Unlike cell culture-derived EV preparations, which are typically enriched for EVs, plasma fractions contain a mixture of EVs and lipoproteins of overlapping sizes. Therefore, applying complementary characterization methods such as NTA and DLS can provide synergistic information and help assess the heterogeneity of particles present in each fraction.

Figure 5 shows SEC Fractions 1, 5 and 9 analyzed using DLS and two generations of NTA instruments (TWIN and QUATT). As expected, SEC separates particles according to size, with larger particles eluting in early fractions and smaller particles in later ones. This is reflected in the DLS results, where Fraction 1 shows a broad range of particles (approximately 100–500 nm), while a subpopulation below 100 nm becomes evident in Fraction 5 and more pronounced in Fraction 9. The NTA measurements show a progressive decrease in total particle concentration across fractions, while the dominant mode size remains within a comparable range. Because NTA does not reliably detect particles below approximately 70 nm, the smaller particle populations seen in the DLS profiles are not captured. This reflects an inherent feature of the NTA technique. However, as this manuscript focuses on the PHoNUPS software rather than comparing instrument performance, we do not elaborate further on this point. Importantly, the two NTA instruments yield proportional and consistent results, indicating that data from different NTA generations can be jointly analyzed using PHoNUPS.

**Figure 5.**
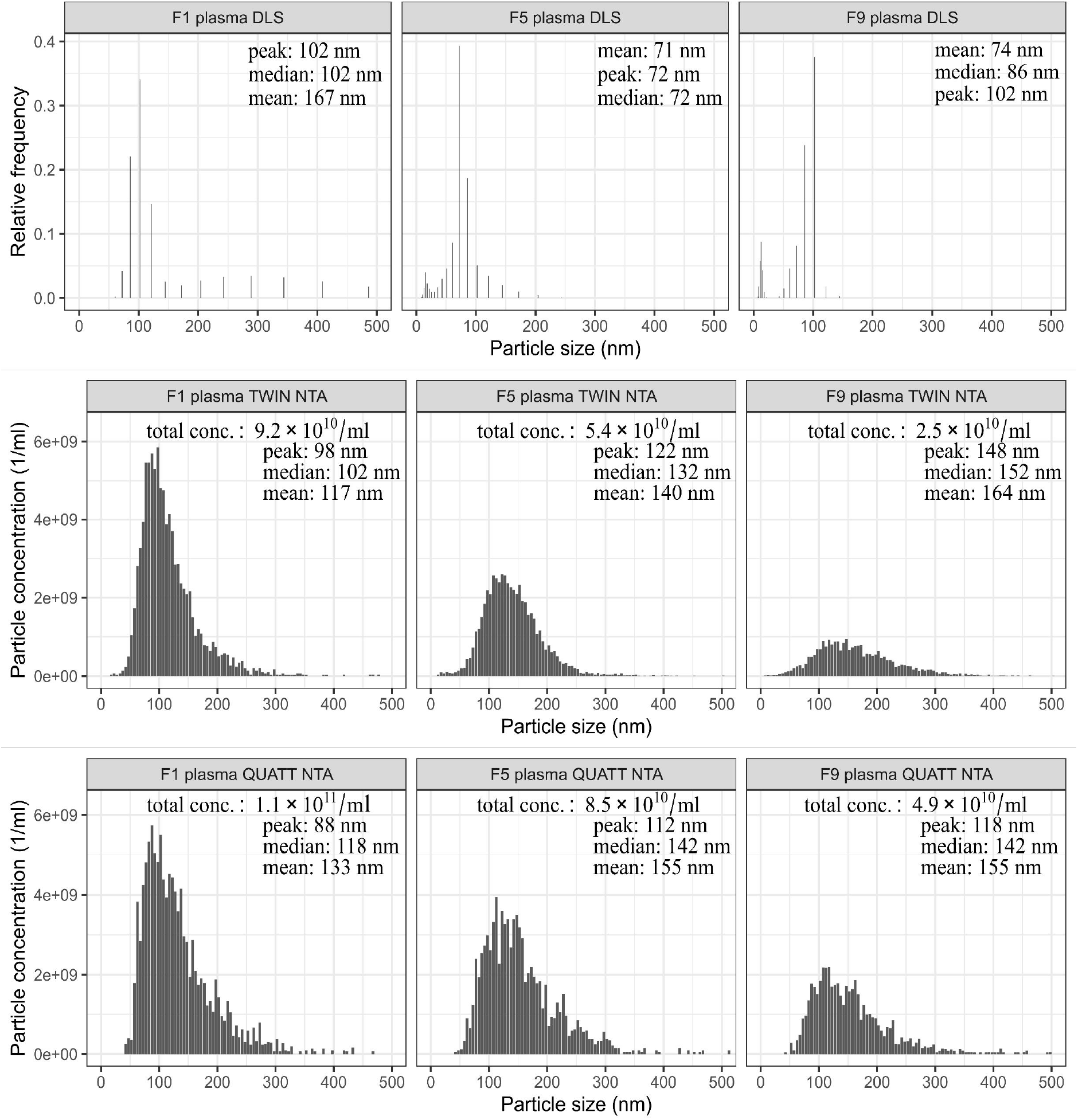
PHoNUPS displays data for the representation of plasma-derived EV Fractions 1, 5 and 9 after SEC, measured by DLS, TWIN and QUATT NTA. Fraction 1 is considered as EV enriched fraction and Fraction 9 is considered as non-EV fraction. Fraction 5 is a mixture of EVs and lipoproteins but the presence of EVs is much lower than in Fraction 1.

Similarly to Example 2, we see again that DLS reports relative particle frequencies with logarithmic binning (with bin centers following a geometric progression), while NTA reports absolute particle concentrations with linear binning. In contrast, DLS operates across a broader size range. PHoNUPS accommodates these intrinsic differences and visualizes the datasets in a unified and comparable format, preserving each instrument’s native units and dimensionality.

We conclude that PHoNUPS enables direct, side-by-side visual comparison of EV SEC fractions or different isolations across complementary platforms within a single figure, using shared axes and standardized graphical formatting.

### Example 4. Zeta Potential Measurements and Visualization Options Using Histograms and Size–Zeta Potential Contour Plots

Zeta potential reflects the electrostatic surface charge of extracellular vesicles (EVs), which is determined by their membrane lipid and protein composition, cargo exposure, and interactions with the surrounding medium. As a complementary parameter to size measurements, zeta potential provides insight into EV preparation consistency, batch-to-batch reproducibility, and overall sample quality. Therefore the standardized visualization of zeta potential distributions supports comparative assessment across EV samples and facilitates early identification of deviations in surface charge that may arise from isolation procedures, buffer conditions, or sample handling.

To illustrate how PHoNUPS visualizes zeta potential measurements, we selected EVs, which typically exhibit a moderately negative surface charge, and synthetic nanocapsules with a slightly positive surface charge to ensure a pronounced contrast for demonstration. PHoNUPS first generates zeta potential histograms with a shared scale, enabling direct comparison of surface charge distributions between particle types. As shown in Figure 6A, EVs cluster around a moderately negative mean zeta potential, whereas nanocapsules form a distinctly separate peak at positive values, demonstrating how zeta potential can be used to discriminate particle populations.

**Figure 6.**
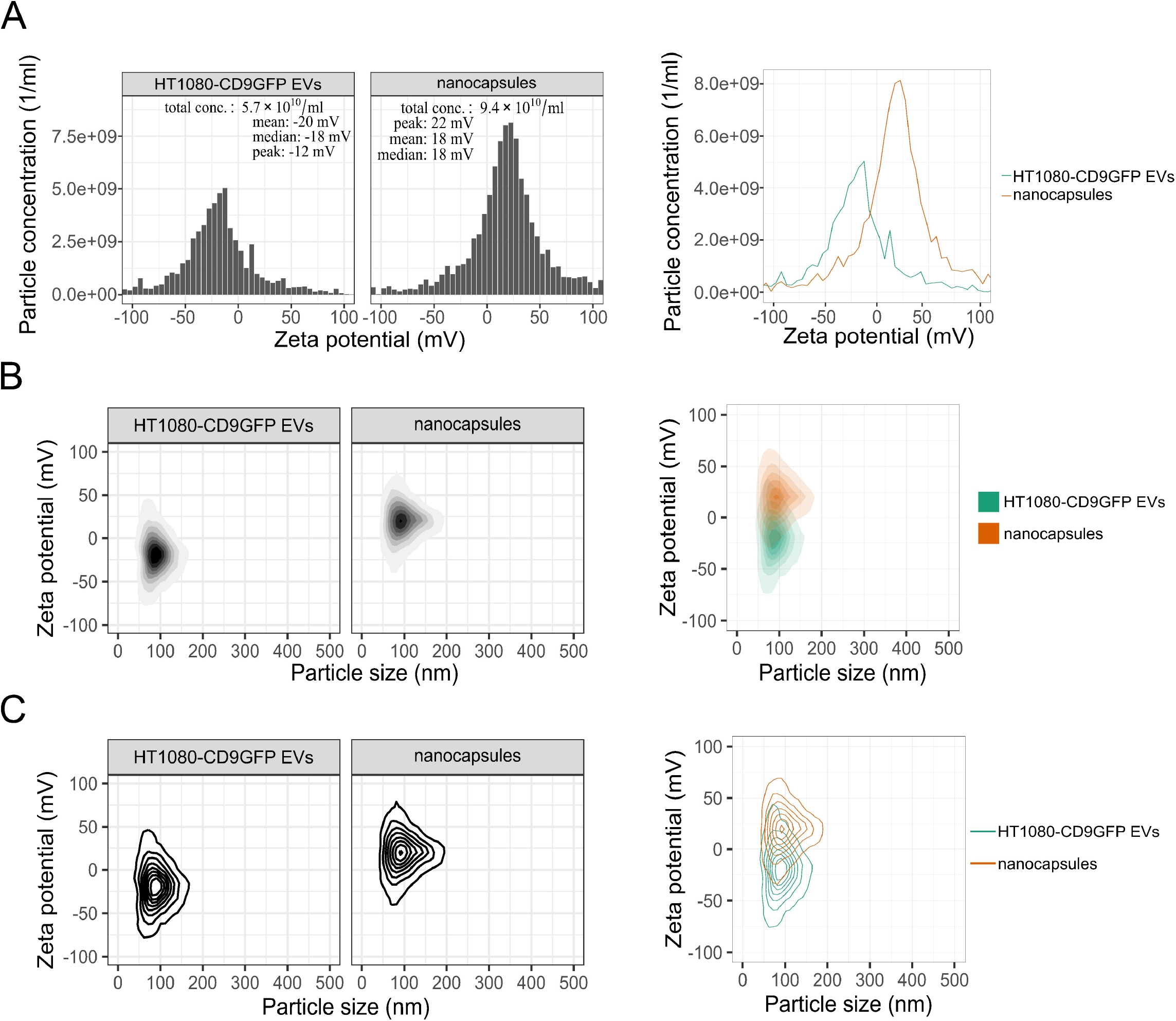
PHoNUPS displays data of zeta potential on the surface of particles. A) Histograms of HT1080-CD9GFP EVs and nanocapsules, which are mainly negatively and mainly positively charged particles, respectively. In size vs. zeta potential contour plots, the user can flip the setting whether to use in the population density visualization B) contour bands with a graded palette, or C) contour lines only.

PHoNUPS further expands this analysis by integrating size and zeta potential into a single visualization when particle-level size and zeta potential data are available. Using kernel density estimation, the software produces 2D contour plots that avoid overplotting and reveal population structure. Two visualization modes are available: filled contour plots (Figure 6B) provide an intuitive heatmap-like representation that highlights density gradients and subpopulation features, while contour line plots (Figure 6C) offer a cleaner format suitable for overlaying multiple datasets within the same figure. Both visualization styles clearly separate EVs from nanocapsules in the size–zeta potential space, demonstrating label-free discrimination between nanoparticle classes.

To summarize, PHoNUPS enables standardized visualization of zeta potential through both histograms and single-particle-based size–charge contour plots, allowing clear discrimination of differently charged particle populations and supporting surface charge-based characterization of EV subtypes.

### Example 5. Using Zeta Potential to Characterize EVs from Cell Culture Supernatants and Biofluids

Zeta potential can be used as a supplementary measurement parameter to assess the surface charge of EVs from different sources and in different media. Furthermore, its applicability extends to complex biofluids, where co-isolation of lipoproteins and other nanosized particles contributes to sample heterogeneity and influences the measured surface charge. In such cases, zeta potential offers additional information that reflects the overall composition of the isolated particle population. To demonstrate this, we present two applications of PHoNUPS: in the first case on EVs isolated from cell culture supernatants, representing a relatively well-defined matrix, and in the second case on EVs isolated from plasma, representing a complex biofluid in which zeta potential can reflect the presence of co-isolated nanoparticle populations.

We first used PHoNUPS to visualize the zeta potential distributions of EVs isolated from different cell lines. Figure 7A shows the corresponding zeta potential histograms, revealing that EVs derived from HT1080, MDA-MB-361 and THP-1 cells display highly similar surface charge profiles, with moderately negative mean zeta potential values within the expected physiological range. This standardized histogram representation allows users to quickly assess distribution shape, central tendency and potential deviations between samples.

**Figure 7.**
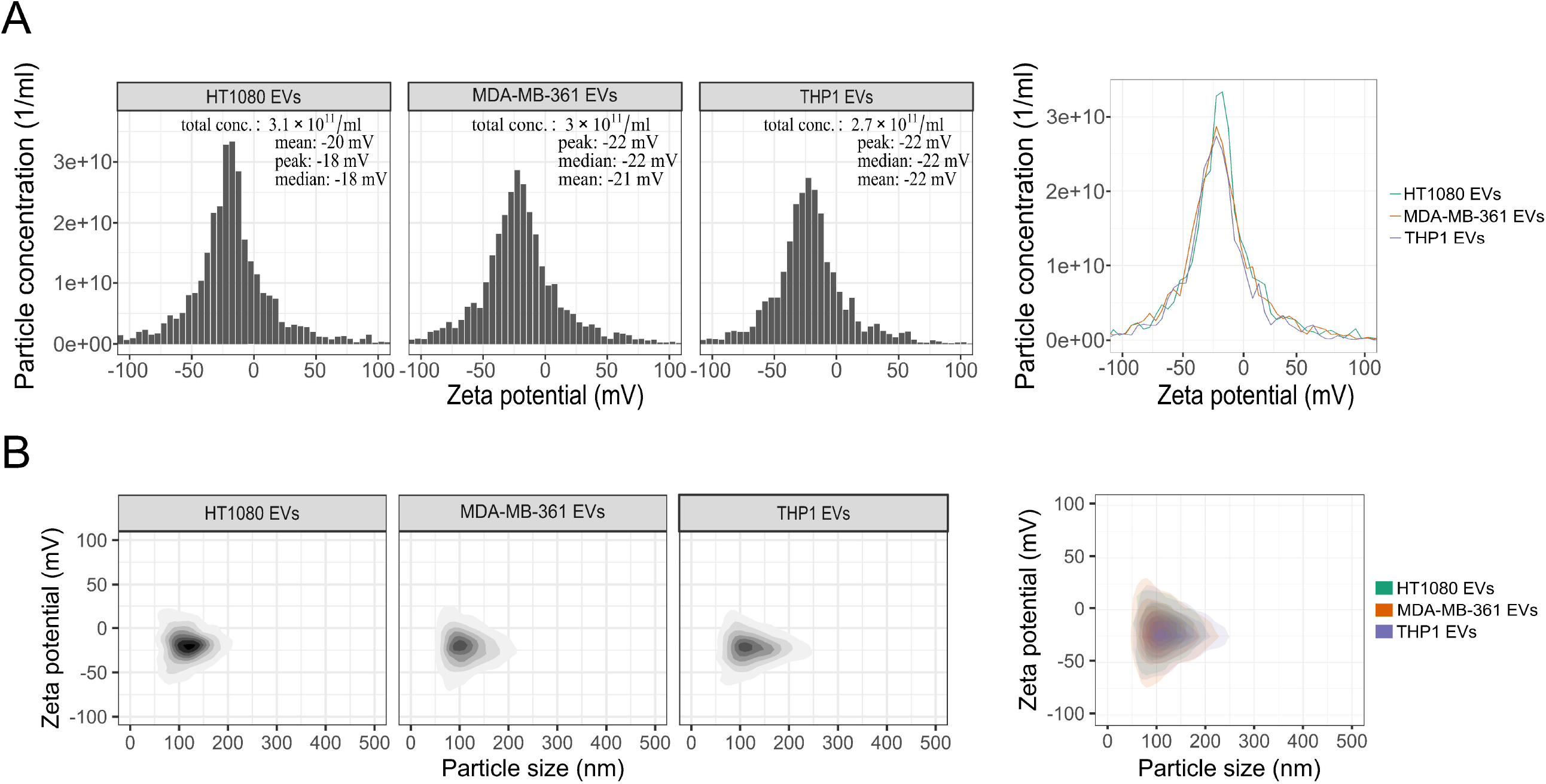
PHoNUPS displays data of zeta potential on the surface of particles in Fraction 1 from EVs derived from three different cell lines, HT1080, MDA-MB-361, and THP-1, isolated using SEC. A) Histograms of particle concentration as a function of zeta potential. B) 2D density contour plots with particle size on the x-axis and zeta potential on the y-axis. After each row of individual plots, their joint plot is shown.

Figure 7B presents the same data in the form of individual size–zeta potential contour plots, providing a 2D view that integrates particle size and charge for each EV preparation. The overlay plot further illustrates the strong similarity across cell-line EVs, demonstrating minimal variation between EV sources. Together, these standardized visual outputs enable users to verify the consistency of EV surface charge across cell-line preparations and support the use of zeta potential as a quality control metric in EV experiments.

We next applied zeta potential analysis to EVs isolated from blood plasma, a complex biofluid where heterogeneity and co-isolation of lipoproteins present a known challenge. EVs were enriched by SEC, and fractions representing three distinct particle compositions were selected: (i) EV-enriched, (ii) lipoprotein-rich with no detectable EVs, and (iii) a pure EV reference. To identify these fractions among nine SEC fractions, fluorescence-assisted SEC followed by fluorescence-NTA was performed (Figure 8A) as described in Nouvel et al. (2024). Because NTA does not reliably detect high-density lipoprotein (HDL) and low-density lipoprotein (LDL) particles (5–25 nm), ApoA1 and ApoB levels were additionally quantified by ELISA to assess lipoprotein abundance (Figure 8B). Based on these measurements, Fraction 1 (F1) was identified as EV-enriched (highest CD9 and CD63 signal with low ApoA1 and ApoB), whereas Fraction 9 (F9) contained no EVs and the highest lipoprotein content. CD9-GFP EVs isolated from HT1080-CD9-GFP cells served as a pure EV control (Supplementary Material 7).

**Figure 8.**
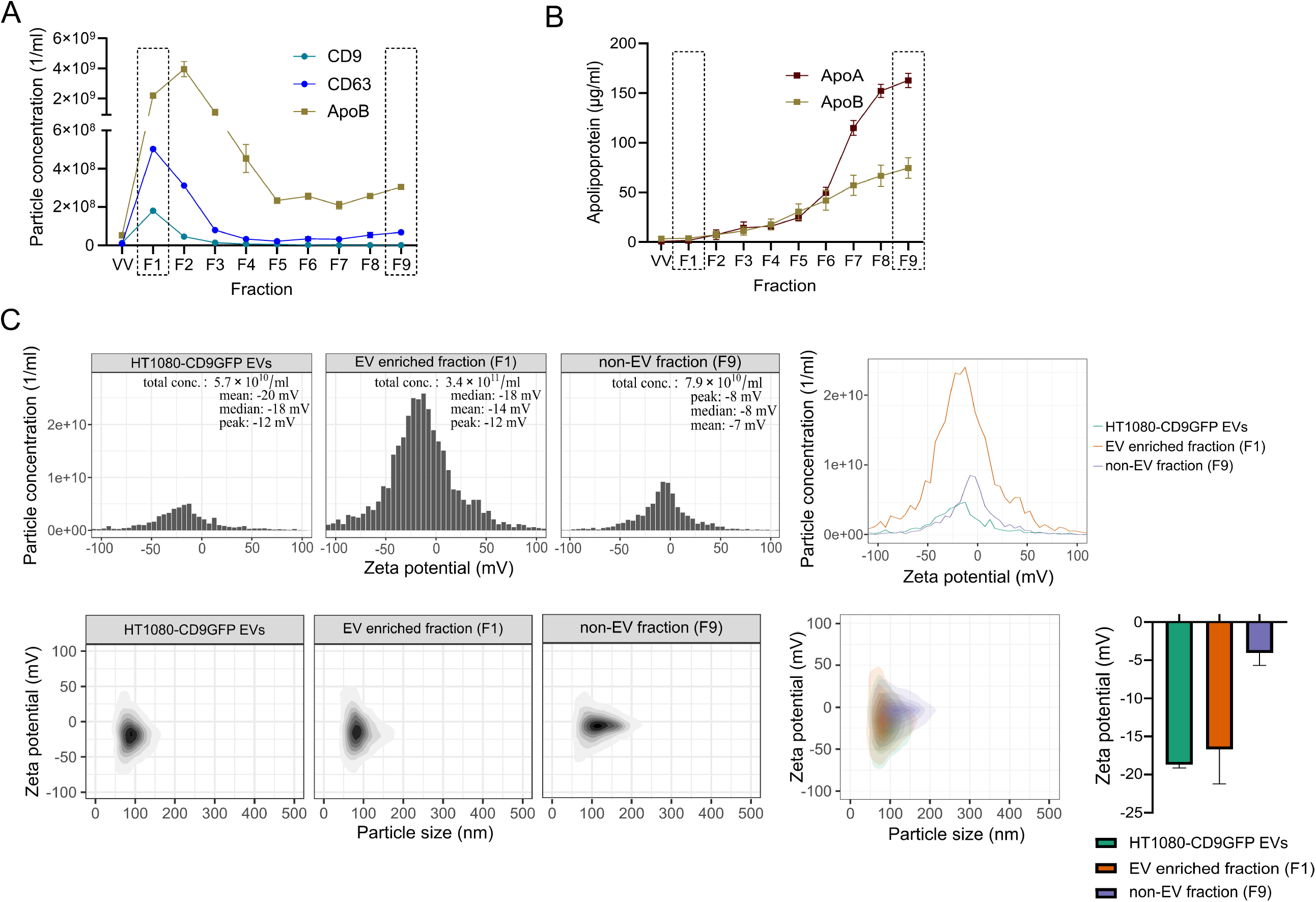
The benefit of the usage of zeta potential to assess EV purity in heterogeneous samples. A) Fluorescent particles measured by NTA of plasma incubated with fluorescent antibodies followed by SEC separation. CD9+ and CD63+ EV are distributed from Fraction 1 to Fraction 3, while lipoproteins labeled with ApoB are co-isolated from Fraction 1 to Fraction 6. B) Apolipoprotein A1 (ApoA1) and B (ApoB) measured by ELISA test in fractions isolated from plasma after SEC. The ApoA1 and ApoB are present in all the fractions, increasing in concentration from Fraction 1, and having the highest value in Fraction 9. C) PHoNUPS displays zeta potential distribution in histograms and individual particle size against zeta potential in contour plots (with the joint plot in the second column) to show the distribution of particle measurements from CD9-GFP EV derived from cell culture, from EV-enriched fraction (Fraction 1) and non-EV fraction (Fraction 9) from plasma samples. Finally, we show the mean zeta potential for the different samples. (VV = void volume)

Using these three samples, we generated zeta potential histograms with PHoNUPS (Figure 8C). Pure EVs displayed the most negative mean zeta potential (–18 mV [SD1]), the EV-enriched plasma fraction (F1) showed an intermediate distribution (–16 mV [SD5]) due to the presence of co-isolated lipoproteins, and the lipoprotein-rich F9 fraction exhibited a broader and markedly less negative profile (–4 mV [SD2]).

PHoNUPS was also used to plot size against zeta potential using single-particle data (Figure 8C). The contour plots clearly distinguished the three particle populations: pure EVs showed more negative zeta potentials (Figure 8C, left panel), the EV-enriched F1 fraction exhibited intermediate values, and the lipoprotein-rich F9 fraction showed the least negative mean charge. The combined overlay (middle panel) revealed distinct spatial distributions, demonstrating how PHoNUPS-based visualization of zeta potential supports interpretation of EV enrichment and co-isolation patterns in complex samples.

In this example, we applied our approach to EV preparations obtained from cell culture supernatants and from biofluids to demonstrate how zeta potential can enhance EV characterization across varying sample complexity. PHoNUPS enables the use of zeta potential as a practical, label-free parameter to assess EV quality and purity across cell line- and biofluid-derived samples. The integration of zeta potential with size data increases the population-level separation of EVs from co-isolated particles such as lipoproteins.

## Conclusion

With PHoNUPS, we introduce an easy-to-use software solution for the standardized processing, statistical analysis, and visualization of non-uniform particle size and zeta potential data, with a particular focus on the needs of the EV research community. By harmonizing outputs from different measurement platforms, PHoNUPS enables direct cross-instrument comparison of EV size distributions through both single-sample and multi-sample histogram visualizations. This supports more consistent reporting practices and facilitates transparent interpretation of EV size profiles across studies and laboratories.

In addition to size-based characterization, PHoNUPS incorporates zeta potential as a supplementary visualization parameter, enabling the comparison of EV surface charge profiles across samples and sample types. When particle-level size and charge data are available, PHoNUPS provides 2D contour plots that depict their joint distribution, allowing intuitive, label-free visualization of particle subpopulations and supporting the interpretation of sample composition and heterogeneity.

PHoNUPS is designed with extensibility in mind. The software already accepts data from multiple instrument types and software versions, and additional input formats can be implemented through modular parser expansion. We invite the EV community to engage with PHoNUPS by contributing new parsers, sharing feature requests, and expanding the platform to cover further analytical modalities relevant to EV research. Through community-driven development, PHoNUPS aims to evolve into a flexible and broadly applicable tool to support transparent and reproducible EV data analysis.

## Material and Methods

### Nanoparticle Tracking Analysis

Two ZetaView x20 Series NTA instruments by Particle Metrix GmbH (Inning am Ammersee, Germany) were used to measure particle concentrations and size distributions. The TWIN was a PMX220 instrument, whereas the QUATT was a PMX220 instrument built according to QUATT specification in pre-production. TWIN NTA software versions used included ZetaView 8.05.14 SP7, 8.05.16 SP3, and their successors PEX 4.2.3.0 and 4.3.2.6. QUATT NTA software versions included ZetaNavigator 1.2.8.1 and 1.4.2.1, which called PEX 4.2.3.0 and 4.3.2.6.

For NTA measurements, samples were diluted to achieve between 80-200 particles per frame for valid measurements. The settings for the TWIN NTA scatter mode were sensitivity 75, shutter 100 (that is, exposure time 1/100 s), trace length 15 s, frame rate 30, 11 positions, minimum size 10 nm, maximum size 1000 nm, laser 488 nm and temperature 25 °C. The settings for the QUATT NTA scatter mode were sensitivity 80, shutter 90, trace length 15 s, frame rate 30, 7 positions, minimum size 10 nm, maximum size 1000 nm, laser 488 nm and temperature 25 °C. On the other hand, the settings for the QUATT NTA fluorescent mode were sensitivity 95, shutter 50, trace length 12 s, frame rate 30, 7 positions, minimum size 10 nm, maximum size 1000 nm, laser 488 nm, filter 500 nm and temperature 25 °C. In the case of the zeta potential, the setting used for the QUATT NTA were sensitivity 80, shutter 90, cycle frames 30, cycle amount 4, frame rate 30, 2 positions (SL1 and SL2), dielectric constant 78.434, Henry factor 1.5, laser 488 nm and temperature 25 °C. All dimensions (size distribution, particle concentration, and zeta potential) were measured using 0.1-time Dulbecco’s phosphate-buffered saline (Gibco DPBS by Thermo Fisher Scientific, Darmstadt, Germany) in Milli-Q H_2_O from an OmniaTap xs basic system by Stakpure (Niederahr, Germany).

### Dynamic Light Scattering

NANO-flex 180° DLS Size by Colloid Metrix GmbH (Meerbusch, Germany) with software Microtrac FLEX 11.1.0.1 was used to measure particle concentrations and size distributions. After performing the set zero on purified Milli-Q H_2_O, the measurement was performed adding a drop of 5 µl on top of the 8 mm diameter probe, and the number of particles was determined to be within the measurement range. The settings were run time 30 s, number of runs 3, multi-run delay 0 min, refractive index water 1.33, low-temperature viscosity 20-1.002 and high-temperature viscosity 30-0.797, respectively. The obtained size distribution was based on the intensity signal, setting standard progression, in the range of 6540 nm as the upper edge and 0.8 nm as the lower edge.

### Flow NanoAnalyzer

Flow NanoAnalyzer (NanoFCM) by NanoFCM Inc. (Xiamen, China) was used to measure the particle concentration of cell culture-derived EVs. Briefly, the instrument was calibrated according to the manufacturer’s instructions. Quality control (QC) beads (NanoFCM Inc.) were used to align the laser and establish the particle concentration, while size beads (68-155 nm, NanoFCM Inc.) were used to calibrate the size. Dulbecco’s phosphate-buffered saline (1× Gibco DPBS, Thermo Fisher Scientific, Darmstadt, Germany) was used as the diluent and blank control, which was then subtracted in each measurement.

EVs were prepared diluted in 1× DPBS to detect between 4000 and 8000 events per measurement. The settings for acquisition were: laser power 10 mW at 488 nm, sampling pressure at 1 kPa, acquisition time of 1 minute and side scatter (SS) decay 10%. PDF and txt file containing data were extracted using the NanoFCM control software NF Profession V2.0 to determine particle concentration and size distribution.

### Enrichment of EVs derived from cell culture supernatants

EVs were isolated from MDA-MB-361, HT1080, HT1080-CD9-GFP, THP-1 and HT29 cell culture supernatant as previously described in Nouvel et al. (2024). Cells were cultured at 37 °C, 5% CO_2_, and 95% humidity in antibiotic-free medium. Briefly, MDA-MB-361 cells were cultured in Dulbecco’s Modified Eagle’s Medium (Gibco DMEM, Thermo Fisher Scientific, Darmstadt, Germany), supplemented with 10% fetal bovine serum (FBS, Thermo Fisher Scientific, Darmstadt, Germany) and 20 mM L-glutamine (Sigma-Aldrich, Schnelldorf, Germany). HT1080, HT1080-CD9-GFP and THP-1 cells were cultured in antibiotic-free Roswell Park Memorial Institute 1640 (Gibco RPMI, Thermo Fisher Scientific, Darmstadt, Germany) supplemented with 10% FBS and 20 mM L-glutamine. HT29 cells were cultured in Dulbecco’s Modified Eagle Medium: Nutrient Mixture F12 (Gibco DMEM F12, Thermo Fisher Scientific, Darmstadt, Germany) supplemented with 10% FBS.

Then, 200 million cells were grown until reaching 80% confluency in T-175 cm^2^ flasks. Cells were washed twice with Dulbecco’s phosphate-buffered saline (1× Gibco DPBS, Thermo Fisher Scientific, Darmstadt, Germany), and new medium without FBS was added. After 24 h, the supernatant was collected, and centrifuged at 800 × *g* for 5 min to remove cells and debris, followed by a second centrifugation at 5000 × *g* for 25 min to eliminate remaining cell debris and apoptotic bodies. Then the supernatant was filtered through a 0.22 µm PES syringe filter (Millex-GP filter, Merck KGaA, Darmstadt, Germany), and the EVs were concentrated using tangential flow filtration (TFF-easy, HansaBiomed Life Sciences Ltd, Tallinn, Estonia) to reduce the volume to approximately 1 ml. With the EVs concentrated, the sample was run through size exclusion chromatography (SEC qEV 35 nm Generation 2, Izon, Christchurch, New Zealand) to enrich EVs in two fractions of 400 µl after running a 2.9 ml of void volume by using an automatic fraction collector (AFC, Izon, Christchurch, New Zealand). Fractions were measured by NTA and DLS.

### Enrichment of EVs derived from plasma (plasma-derived EV, PdEV)

EVs were isolated from plasma from healthy donors. Briefly, blood was collected in 10 ml citrate tubes (S-Monovette, SARSTEDT, Nümbrecht, Germany) and then centrifuged twice at 2500 × *g* at temperature 25 °C, carefully removing the supernatant and transferring it to a new tube after each centrifugation. EVs were isolated by size exclusion chromatography (SEC qEV 35 nm Generation 2, Izon, Christchurch, New Zealand) to enrich EVs in the first fractions after running a 2.9 ml of void volume by using an automatic fraction collector (AFC, Izon, Christchurch, New Zealand). Fractions were measured by NTA and DLS.

To determine the EV content of different SEC fractions, plasma from a healthy donor was incubated 1 h at 4 °C prior to SEC with EV universal biomarker antibodies: CD9-APC (312108, BioLegend, San Diego, CA, USA) and CD63-FITC (353006, BioLegend, San Diego, CA, USA) and then the fluorescent signal was quantified by NTA immediately after running SEC.

To test the lipoprotein-enriched fractions, we tested them by ELISA and NTA. In case of the ELISA test, we used the human ApoA1 kit (894879, R&D Systems, MN, USA) to measure apolipoprotein A1, which is present in high-density lipoproteins (HDL), and the human ApoB kit (894224, R&D Systems, MN, USA) for apolipoprotein B, which is present in very-low-density lipoproteins (VLDL), intermediate-density lipoproteins (IDL), and low-density lipoproteins (LDL). The experiments were performed following the instructions of the supplier. Fractions after SEC were diluted 1:30 in the case of ApoA1 and 1:20 for ApoB, and the absorbance was measured at 450 nm on a Tecan Infinite M PLEX (Männedorf, Switzerland). In the case of fluorescence measurement on NTA, plasma samples were labeled 1 h long at 4 °C prior to SEC with ApoB-FITC antibody (ab27637, Abcam Limited, Cambridge, UK) and thereafter the fluorescent particles were immediately measured.

### Nanocapsules

Nanocapsules (Capsulon™) were donated by CapCo Bio GmbH (Freiburg, Germany). The nanocapsules were produced as previously described by Tarakanchikova et al. (2020).

### Statistical analysis

GraphPad Prism 10.4.2 version (GraphPad Software, Boston, MA, USA) was used to generate complementary plots.

## Author contributions

Bence Mélykúti: software programming, writing—original draft, writing—review and editing. Gonzalo Bustos-Quevedo: conducting experiments, data curation, investigation, writing—original draft, writing—review and editing. Tony Prinz: conceptualization. Irina Nazarenko: conceptualization, writing—review and editing, supervision, funding acquisition.

## Acknowledgements

We thank Tanja Gainey-Schleicher and Sophie Krüger for their technical assistance. We acknowledge the support of the Facility for Extracellular Vesicle Analysis and Liquid Biopsy (EV-Core), Medical Center – University of Freiburg (registered with the German Research Foundation [DFG] under RI_00612), and the financial support it received from the Faculty of Medicine – University of Freiburg. We gratefully thank Particle Metrix GmbH (Inning am Ammersee, Germany) for providing instruments.

The development reported in this publication was supported by the German Federal Ministry for Economic Affairs and Climate Action (BMWK, project Next-GenNTA KK518401), the German Federal Ministry for Research, Technology and Space (BMFTR, projects EV-Surf 13GW0605F and Nanodiag 03ZU1208CA), the State Ministry for Science, Research and Arts Baden-Württemberg (MWK, project NaPeGen MWK33-7532-50/1/3), and the European Union HORIZON-EIC-2021-TRANSITIONOPEN-01 (project Nexus 101058200).

## Supporting information

Supplement.docx

