## Supplemental text for "PHoNUPS: Open-Source Software for Standardized Analysis and Visualization of Multi-Instrument Extracellular Vesicle Measurements"

### Report formats of some instruments

The Supplementary Material shows the diverse PDF report document formats of the Particle Metrix ZetaView x20 Series PMX-220 NTA (TWIN and QUATT), the Colloid Metrix NANO-flex 180° DLS Size, and the NanoFCM Flow NanoAnalyzer instruments, and the data visualizations that they contain.

Supplementary Material 1 shows the reports corresponding to the measurements of Figure 3. The TWIN has an older software version, which displays the histogram with a line plot with a solid curve, while the newer software used by the QUATT generates bar charts for histograms, yielding higher resolution.

The statistics in the reports are close enough to the values reported by PHoNUPS, but they are not expected to be identical. The reason is that PHoNUPS uses the binned size measurements (how many particles are closest in size to a particular value from a list of bin center values), not the size measurements of each individual particle.

Supplementary Material 1 contains the reports in the following order:

- Pages 1, 2: TWIN NTA reports for Figure 3B.
- Pages 3, 4: QUATT NTA reports for Figure 3B.
- Pages 5, 6: QUATT NTA reports for Figure 3C, scatter mode, two technical replicates.
- Pages 7, 8: QUATT NTA reports for Figure 3C, fluorescent mode, two technical replicates.

Supplementary Material 2 shows the reports corresponding to the measurements of Figure 4, with two technical replicates in each case, in the following order:

- Pages 1, 2: DLS reports for Figure 4A.
- Pages 3, 4: TWIN NTA reports for Figure 4A.
- Pages 5, 6: QUATT NTA reports for Figure 4A.
- Pages 7, 8: QUATT NTA reports for Figure 4B and Figure 4D.
- Pages 9: NanoFCM report for Figure 4C and Figure 4D.

Supplementary Material 3 shows the reports corresponding to the two technical replicate measurements for each sample of Figure 5 with DLS, TWIN and QUATT NTA:

- Pages 1–6: DLS reports for Figure 5, first row.
- Pages 7–12: TWIN NTA reports for Figure 5, second row.
- Pages 13–18: QUATT NTA reports for Figure 5, third row.

In Supplementary Material 3, on Page 5, in *- Measurement Info -*, the field *Title* should read *f9_GBQ_serum_onhe_label_1* instead of *F8_GBQ_EDTA_onhe_label* for consistency, but the instrument control software did not allow this correction.

Supplementary Material 4 shows the reports corresponding to the two technical replicate measurements for both samples of Figure 6 with QUATT NTA.

- Pages 1, 2: Reports for Figure 6, EVs.
- Pages 3, 4: Reports for Figure 6, nanocapsules.

Supplementary Material 5 shows the reports corresponding to the measurements of Figure 7 with QUATT NTA, with two technical replicates in each case, in the following order:

- Pages 1, 2: Reports for Figure 7, HT1080 EVs.
- Pages 3, 4: Reports for Figure 7, MDA-MB-361 EVs.
- Pages 5, 6: Reports for Figure 7, THP-1 EVs.

Supplementary Material 6 shows the reports corresponding to the measurements of Figure 8 with QUATT NTA, with two technical replicates in each case, in the following order:

- Pages 1, 2: Reports for Figure 8C, HT1080-CD9-GFP EVs.
- Pages 3, 4: Reports for Figure 8C, plasma fraction 1.
- Pages 5, 6: Reports for Figure 8C, plasma fraction 9.

### Example 5. Using Zeta Potential to Characterize EVs from Cell Culture Supernatants and Biofluids

Supplementary Material 7 corresponds to the characterization of reference EVs derived from HT1080-CD9-GFP used in Figure 8C. A) NTA analysis of fractions obtained using size exclusion chromatography (SEC). Fractions 1 and 2 contained maximal particle number in both scatter and fluorescent modes. B) Size distribution of Fraction 1 by PHoNUPS showing that the particles quantified in scatter and fluorescent mode correspond to the same particle population.
