## Supplementary Material 1 for "PHoNUPS: Open-Source Software for Standardized Analysis and Visualization of Multi-Instrument Extracellular Vesicle Measurements"

PARTICLE METRIX **ZETAVIEW®** www.particle-metrix.com

Electrophoresis & Brownian Motion
Video Analysis
Laser Scattering Microscopy

Video Operator: ZetaUser

Operator (Report): ZetaUser

**Sample Parameters**
Sample Name:
Sample Info 1:
Sample Info 2:
Sample Info 3:
Electrolyte: $H_2O$
Temperature: 24.66 °C sensed
pH 7.0 entered
Conductivity: 1307.00 µS/cm sensed

**Instrument Parameters**
Laser Wavelength: 488 nm
Filter Wavelength: Scatter
Sensitivity: 75
Shutter: 100

**SOP: SOP100_ExTu**
Size Distribution 1 Cycle 11 Positions
Description:

**Result (sizes in nm)**

| | Number | Concentration | Volume |
|---|---|---|---|
| Median (X50) | 126.5 | 126.5 | 261.5 |
| StdDev | 73.1 | 73.1 | 134.7 |

Concentration: 6.5E+7 Particles / mL
Dilution Factor: 8000
Original Concentration: 5.2E+11 Particles / mL

**Quality**
Average Counted Particles per Frame: 178
Number of Traced Particles: 1956
2 Positions Removed for Analysis

**Analysis Parameters**
Max Area 1000, Min Area 10, Min Brightness 20, nm/Class: 5

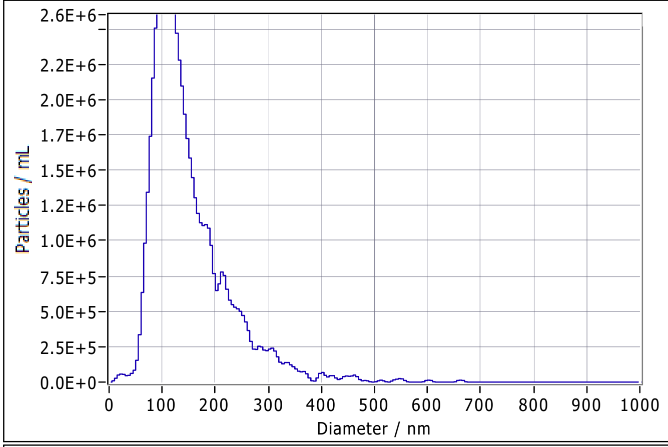


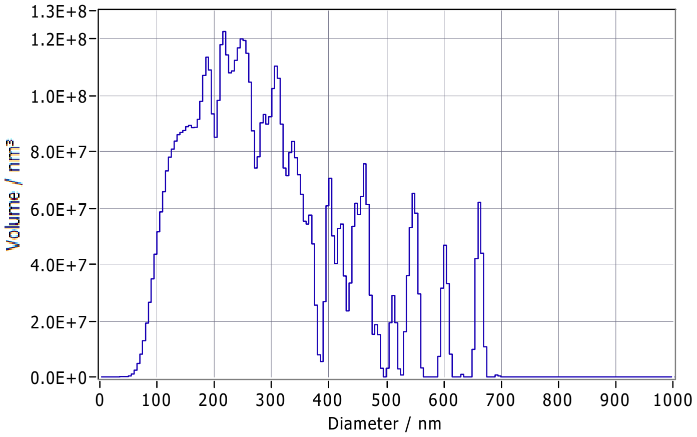


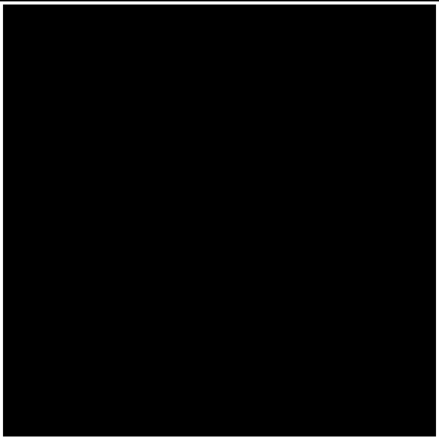

**Peak Analysis (Concentration)**

| Diameter / nm | Particles/mL | FWHM / nm | Percentage |
|---|---|---|---|
| 105.3 | 3.0E+6 | 80.4 | 100.0 |

**X Values** (all sizes are given in nm)

| | Number | Concentration | Volume |
|---|---|---|---|
| X10 | 79.6 | 79.6 | 133.3 |
| X50 | 126.5 | 126.5 | 261.5 |
| X90 | 242.0 | 242.0 | 466.7 |
| Span | 1.3 | 1.3 | 1.3 |
| Mean | 149.4 | 149.4 | 291.2 |
| StdDev | 73.1 | 73.1 | 134.7 |

**Comment**

(Signature)

Analyzed Video: Z:\Tanja\20240702\20240702_0014_iso1F1_1_8000_size_488.avi

Experiment: 2024-07-02 10:10 ZetaView S/N 20-559, Software ZetaView (version 8.05.16 SP3)

Report: 2024-07-02 10:12 Software ZetaView (version 8.05.16 SP3)

### ZETAVIEW®

PARTICLE METRIX

www.particle-metrix.com

Electrophoresis & Brownian Motion
Video Analysis
Laser Scattering Microscopy

Video Operator: ZetaUser

Operator (Report): ZetaUser

**Sample Parameters**
Sample Name:
Sample Info 1:
Sample Info 2:
Sample Info 3:
Electrolyte: $H_2O$
Temperature: 24.70 °C sensed
pH 7.0 entered
Conductivity: 1304.00 µS/cm sensed

**Instrument Parameters**
Laser Wavelength: 488 nm
Filter Wavelength: Scatter
Sensitivity: 75
Shutter: 100

**SOP: SOP100_ExTu**
Size Distribution 1 Cycle 11 Positions
Description:

**Result (sizes in nm)**

| | Number | Concentration | Volume |
|---|---|---|---|
| Median (X50) | 115.2 | 115.2 | 186.4 |
| StdDev | 52.0 | 52.0 | 106.3 |

Concentration: 6.6E+7 Particles / mL
Dilution Factor: 8000
Original Concentration: 5.2E+11 Particles / mL

**Quality**
Average Counted Particles per Frame: 180
Number of Traced Particles: 1466
3 Positions Removed for Analysis

**Analysis Parameters**
Max Area 1000, Min Area 10, Min Brightness 20, nm/Class: 5

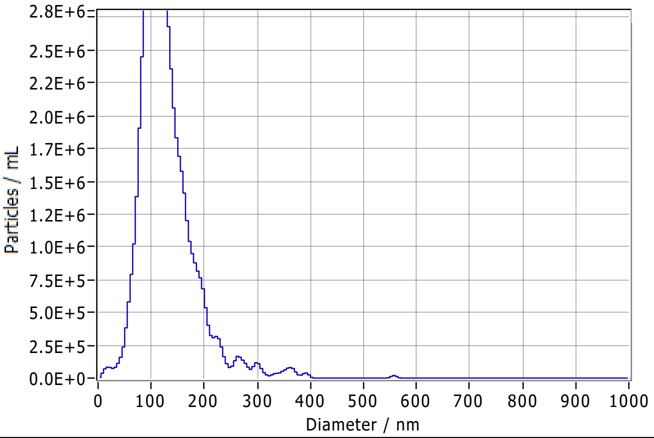


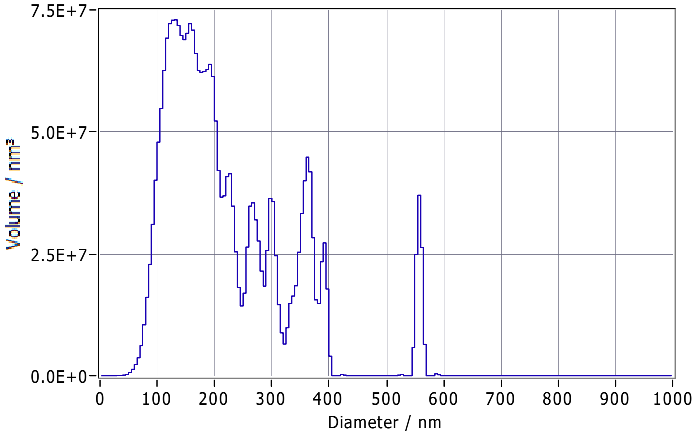


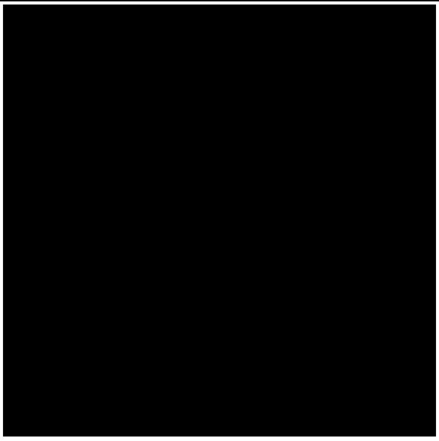

**Peak Analysis (Concentration)**

| Diameter / nm | Particles/mL | FWHM / nm | Percentage |
|---|---|---|---|
| 107.4 | 3.8E+6 | 68.6 | 100.0 |

**X Values** (all sizes are given in nm)

| | Number | Concentration | Volume |
|---|---|---|---|
| X10 | 76.7 | 76.7 | 109.7 |
| X50 | 115.2 | 115.2 | 186.4 |
| X90 | 186.3 | 186.3 | 362.5 |
| Span | 1.0 | 1.0 | 1.4 |
| Mean | 128.2 | 128.2 | 219.2 |
| StdDev | 52.0 | 52.0 | 106.3 |

**Comment**

(Signature)

Analyzed Video: Z:\Tanja\20240702\20240702_0020_iso2F1_1_8000_size_488.avi

Experiment: 2024-07-02 10:31 ZetaView S/N 20-559, Software ZetaView (version 8.05.16 SP3)

Report: 2024-07-02 10:33 Software ZetaView (version 8.05.16 SP3)

e_id: 2888, ZetaScript: SOP new fluoro/coloco/GREEN/scatter 100, Sample name: F1_CD9gfp_hamburg_23.06.202

Measurement created: 2024-07-02 11:00:20 Report created: 2024-07-02 11:29:45 User: ZetaUser

#### Measurement specification: SOP100(1)

##### Capture settings

| | | | |
|---|---|---|---|
| Temperature / °C | 25.0 | Laser wave length / nm | 488 |
| Dilution | 8000 | Filter wave length / nm | 0 |
| Viscosity / mPa · s | 0.899 | Sensitivity | 80 |
| Positions | None | Shutter | 90 |
| Video length | 30 | | |
| Frame rate / 1/s | 30.303 | | |
| Experiment type | Size distribution | | |

##### Analysis settings

| | |
|---|---|
| Concentration correction | 0.55 |
| Concentration calibration | 636400 |
| Min size /nm | 10 |
| Max size / nm | 1000 |
| Min area / px | 1 |
| Max area / px | 1000 |
| Trace length | 15 |

Comment:

#### Graphics:

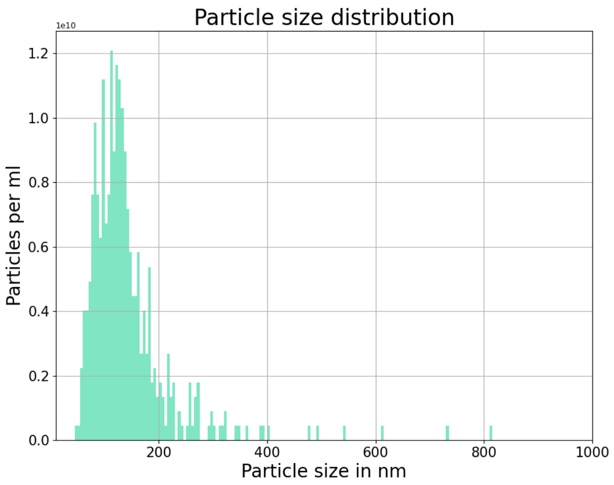


#### Results:

| | Peak /nm | Median /nm | #Particles | Avg. Num Det. | FWHM /nm | Concentration /1/mL | Position |
|---|---|---|---|---|---|---|---|
| | **115.4 (± 14.2)** | **124.0 (± 10.2)** | **468** | **74.8 (± 7.5)** | **93.6 (± 26.9)** | **2.10E+11 (± 2.1E+10)** | |
| 1 | 115.0 | 123.8 | 70 | 74.0 | 83.6 | 2.07E+11 | 0.30 |
| 2 | 110.1 | 117.1 | 74 | 85.0 | 102.9 | 2.38E+11 | 0.40 |
| 3 | 85.1 | 104.8 | 66 | 78.9 | 92.3 | 2.21E+11 | 0.50 |
| 4 | 117.9 | 122.4 | 66 | 75.6 | 75.5 | 2.12E+11 | 0.60 |
| 5 | 126.7 | 137.7 | 69 | 75.2 | 152.3 | 2.11E+11 | 0.70 |
| 6 | 119.6 | 128.2 | 75 | 76.8 | 87.8 | 2.15E+11 | 0.80 |
| 7 | 133.1 | 134.0 | 48 | 58.4 | 61.0 | 1.63E+11 | 0.85 |

e_id: 2892, ZetaScript: SOP new fluoro/coloco/GREEN/scatter 100, Sample name: F1_iso2_CD9gfp_hamburg_23.06.2[illegible]

Measurement created: 2024-07-02 11:14:01 Report created: 2024-07-02 11:29:47 User: ZetaUser

#### Measurement specification: SOP100(1)

##### Capture settings

| | | | |
|---|---|---|---|
| Temperature / °C | 25.1 | Laser wave length / nm | 488 |
| Dilution | 8000 | Filter wave length / nm | 0 |
| Viscosity / mPa · s | 0.897 | Sensitivity | 80 |
| Positions | None | Shutter | 90 |
| Video length | 30 | | |
| Frame rate / 1/s | 30.303 | | |
| Experiment type | Size distribution | | |

##### Analysis settings

| | |
|---|---|
| Concentration correction | 0.55 |
| Concentration calibration | 636400 |
| Min size /nm | 10 |
| Max size / nm | 1000 |
| Min area / px | 1 |
| Max area / px | 1000 |
| Trace length | 15 |

Comment:

#### Graphics:

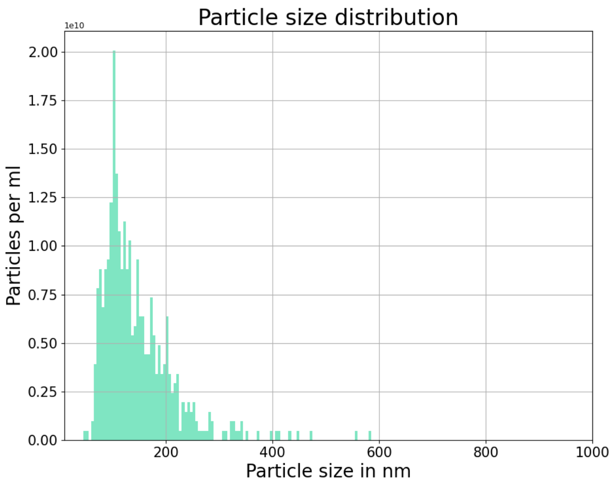


#### Results:

| | Peak /nm | Median /nm | #Particles | Avg. Num Det. | FWHM /nm | Concentration /1/mL | Position |
|---|---|---|---|---|---|---|---|
| | **109.3 (± 5.0)** | **128.6 (± 5.5)** | **525** | **91.8 (± 8.5)** | **113.3 (± 22.7)** | **2.57E+11 (± 2.4E+10)** | |
| 1 | 107.3 | 119.6 | 83 | 89.5 | 83.6 | 2.51E+11 | 0.30 |
| 2 | 108.4 | 130.3 | 69 | 89.4 | 124.9 | 2.50E+11 | 0.40 |
| 3 | 105.9 | 133.7 | 76 | 107.3 | 144.1 | 3.00E+11 | 0.50 |
| 4 | 111.2 | 125.3 | 75 | 90.4 | 83.2 | 2.53E+11 | 0.60 |
| 5 | 104.0 | 123.9 | 58 | 76.4 | 118.2 | 2.14E+11 | 0.70 |
| 6 | 107.6 | 136.2 | 78 | 95.4 | 137.3 | 2.67E+11 | 0.80 |
| 7 | 120.4 | 131.3 | 86 | 93.9 | 101.9 | 2.63E+11 | 0.85 |

### e_id: 3309, ZetaScript: SOP FM GBQ, Sample name: CD9-GFP_nexus_isolation_after_isolation_1_500

Measurement created: 2024-09-25 19:11:52 Report created: 2024-09-25 19:26:00 User: ZetaUser

#### Measurement specification: SOP100(1)

##### Capture settings

| | | | |
|---|---|---|---|
| Temperature / °C | 25.1 | Laser wave length / nm | 488 |
| Dilution | 500 | Filter wave length / nm | 0 |
| Viscosity / mPa · s | 0.898 | Sensitivity | 80 |
| Positions | None | Shutter | 90 |
| Video length | 30 | | |
| Frame rate / 1/s | 30.303 | | |
| Experiment type | Size distribution | | |

##### Analysis settings

| | |
|---|---|
| Concentration correction | 0.55 |
| Concentration calibration | 636400 |
| Min size /nm | 10 |
| Max size / nm | 1000 |
| Min area / px | 1 |
| Max area / px | 1000 |
| Trace length | 15 |

Comment:

#### Graphics:

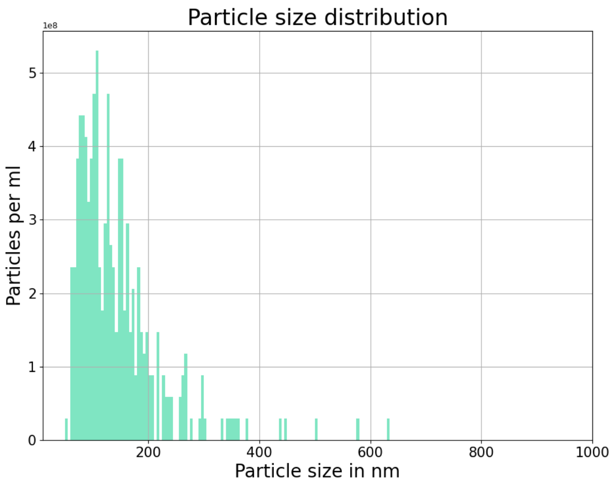


#### Results:

| | Peak /nm | Median /nm | #Particles | Avg. Num Det. | FWHM /nm | Concentration /1/mL | Position |
|---|---|---|---|---|---|---|---|
| | **112.4 (± 18.3)** | **125.2 (± 13.8)** | **320** | **53.9 (± 6.4)** | **121.0 (± 19.7)** | **9.43E+09 (± 1.1E+09)** | |
| 1 | 108.4 | 125.5 | 34 | 41.7 | 115.9 | 7.29E+09 | 0.30 |
| 2 | 108.2 | 136.1 | 52 | 62.1 | 129.7 | 1.09E+10 | 0.40 |
| 3 | 154.1 | 147.7 | 45 | 50.7 | 123.9 | 8.88E+09 | 0.50 |
| 4 | 118.8 | 132.2 | 44 | 50.2 | 135.3 | 8.79E+09 | 0.60 |
| 5 | 99.2 | 119.4 | 48 | 55.6 | 112.0 | 9.74E+09 | 0.70 |
| 6 | 99.2 | 107.6 | 47 | 58.8 | 148.6 | 1.03E+10 | 0.80 |
| 7 | 98.9 | 107.6 | 50 | 58.0 | 81.5 | 1.01E+10 | 0.85 |

### e_id: 3309, ZetaScript: SOP FM GBQ, Sample name: CD9-GFP_nexus_isolation_after_isolation_1_500

Measurement created: 2024-09-25 19:11:52 Report created: 2024-09-25 19:26:00 User: ZetaUser

#### Measurement specification: SOP100(2)

##### Capture settings

| | | | |
|---|---|---|---|
| Temperature / °C | 25.1 | Laser wave length / nm | 488 |
| Dilution | 500 | Filter wave length / nm | 0 |
| Viscosity / mPa · s | 0.898 | Sensitivity | 80 |
| Positions | None | Shutter | 90 |
| Video length | 30 | | |
| Frame rate / 1/s | 30.303 | | |
| Experiment type | Size distribution | | |

##### Analysis settings

| | |
|---|---|
| Concentration correction | 0.55 |
| Concentration calibration | 636400 |
| Min size /nm | 10 |
| Max size / nm | 1000 |
| Min area / px | 1 |
| Max area / px | 1000 |
| Trace length | 15 |

Comment:

#### Graphics:

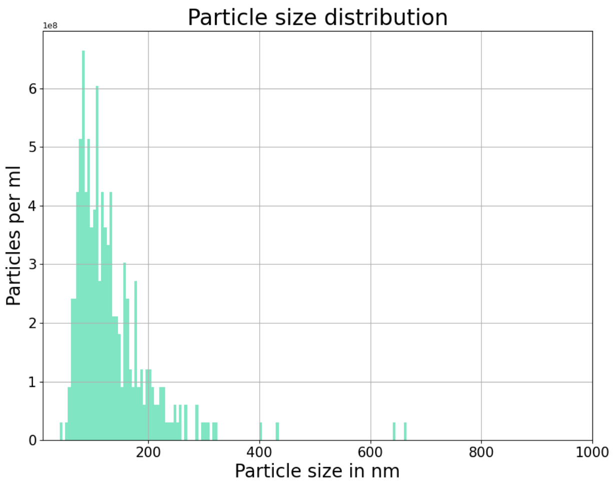


#### Results:

| | Peak /nm | Median /nm | #Particles | Avg. Num Det. | FWHM /nm | Concentration /1/mL | Position |
|---|---|---|---|---|---|---|---|
| | **100.4 (± 9.6)** | **112.7 (± 10.2)** | **318** | **54.9 (± 6.3)** | **104.9 (± 16.8)** | **9.61E+09 (± 1.1E+09)** | |
| 1 | 103.9 | 118.8 | 47 | 48.4 | 83.0 | 8.47E+09 | 0.30 |
| 2 | 85.4 | 94.5 | 44 | 51.5 | 102.7 | 9.01E+09 | 0.40 |
| 3 | 120.1 | 129.6 | 36 | 47.1 | 130.7 | 8.24E+09 | 0.50 |
| 4 | 97.6 | 107.5 | 46 | 57.9 | 91.7 | 1.01E+10 | 0.60 |
| 5 | 97.3 | 115.7 | 54 | 63.9 | 115.1 | 1.12E+10 | 0.70 |
| 6 | 98.7 | 107.3 | 47 | 63.3 | 88.9 | 1.11E+10 | 0.80 |
| 7 | 99.9 | 115.8 | 44 | 52.4 | 122.3 | 9.17E+09 | 0.85 |

### e_id: 3309, ZetaScript: SOP FM GBQ, Sample name: CD9-GFP_nexus_isolation_after_isolation_1_500

Measurement created: 2024-09-25 19:11:52 Report created: 2024-09-25 19:25:59 User: ZetaUser

#### Measurement specification: TP_1_GREEN_SOP_100_95int_sh50_4

##### Capture settings

| | | | |
|---|---|---|---|
| Temperature / °C | 25.0 | Laser wave length / nm | 488 |
| Dilution | 500 | Filter wave length / nm | 500 |
| Viscosity / mPa · s | 0.9 | Sensitivity | 95 |
| Positions | None | Shutter | 50 |
| Video length | 30 | | |
| Frame rate / 1/s | 30.303 | | |
| Experiment type | Size distribution | | |

##### Analysis settings

| | |
|---|---|
| Concentration correction | 0.55 |
| Concentration calibration | 636400 |
| Min size /nm | 10 |
| Max size / nm | 1000 |
| Min area / px | 1 |
| Max area / px | 1000 |
| Trace length | 12 |

Comment:

#### Graphics:

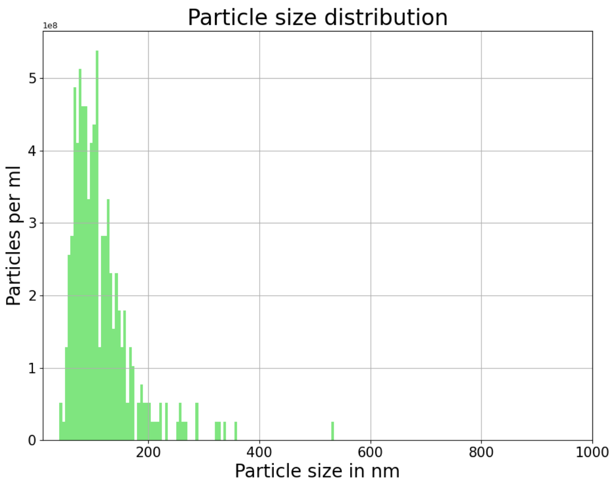


#### Results:

| | Peak /nm | Median /nm | #Particles | Avg. Num Det. | FWHM /nm | Concentration /1/mL | Position |
|---|---|---|---|---|---|---|---|
| | **88.0 (± 4.2)** | **103.0 (± 4.8)** | **311** | **45.6 (± 4.4)** | **93.1 (± 12.3)** | **7.98E+09 (± 7.7E+08)** | |
| 1 | 86.1 | 102.3 | 43 | 44.1 | 104.2 | 7.72E+09 | 0.20 |
| 2 | 87.0 | 98.9 | 43 | 42.8 | 81.6 | 7.48E+09 | 0.30 |
| 3 | 87.4 | 105.3 | 64 | 54.9 | 103.7 | 9.61E+09 | 0.40 |
| 4 | 96.8 | 109.2 | 34 | 47.1 | 111.7 | 8.24E+09 | 0.50 |
| 5 | 85.2 | 107.6 | 37 | 39.7 | 90.7 | 6.95E+09 | 0.60 |
| 6 | 83.0 | 94.1 | 46 | 46.0 | 80.1 | 8.05E+09 | 0.70 |
| 7 | 90.7 | 104.0 | 44 | 44.5 | 79.9 | 7.79E+09 | 0.80 |

e_id: 3309, ZetaScript: SOP FM GBQ, Sample name: CD9-GFP_nexus_isolation_after_isolation_1_500

Measurement created: 2024-09-25 19:11:52    Report created: 2024-09-25 19:25:59    User: ZetaUser

#### Measurement specification: TP_1_GREEN_SOP_100_95int_sh50_3

##### Capture settings

| | | | |
|---|---|---|---|
| Temperature / °C | 25.1 | Laser wave length / nm | 488 |
| Dilution | 500 | Filter wave length / nm | 500 |
| Viscosity / mPa · s | 0.898 | Sensitivity | 95 |
| Positions | None | Shutter | 50 |
| Video length | 30 | | |
| Frame rate / 1/s | 30.303 | | |
| Experiment type | Size distribution | | |

##### Analysis settings

| | |
|---|---|
| Concentration correction | 0.55 |
| Concentration calibration | 636400 |
| Min size /nm | 10 |
| Max size / nm | 1000 |
| Min area / px | 1 |
| Max area / px | 1000 |
| Trace length | 12 |

Comment:

#### Graphics:

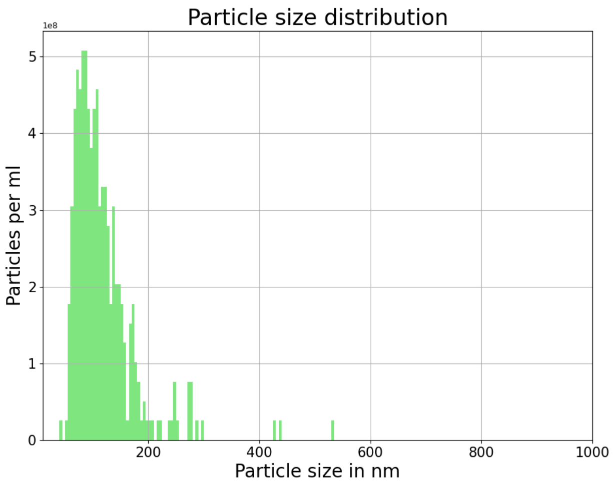


#### Results:

| | Peak /nm | Median /nm | #Particles | Avg. Num Det. | FWHM /nm | Concentration /1/mL | Position |
|---|---|---|---|---|---|---|---|
| | **92.8 (± 7.2)** | **104.8 (± 6.2)** | **324** | **47.0 (± 3.7)** | **89.9 (± 12.9)** | **8.23E+09 (± 6.5E+08)** | |
| 1 | 95.3 | 105.5 | 44 | 41.9 | 79.5 | 7.34E+09 | 0.20 |
| 2 | 81.3 | 94.9 | 47 | 45.5 | 92.3 | 7.96E+09 | 0.30 |
| 3 | 90.0 | 110.7 | 45 | 48.1 | 87.9 | 8.41E+09 | 0.40 |
| 4 | 105.1 | 111.8 | 51 | 52.9 | 107.1 | 9.25E+09 | 0.50 |
| 5 | 97.4 | 103.9 | 48 | 45.5 | 76.0 | 7.96E+09 | 0.60 |
| 6 | 93.8 | 110.0 | 42 | 43.9 | 109.8 | 7.69E+09 | 0.70 |
| 7 | 86.5 | 96.9 | 47 | 51.6 | 77.0 | 9.02E+09 | 0.80 |
