## Supplementary Material 2 for "PHoNUPS: Open-Source Software for Standardized Analysis and Visualization of Multi-Instrument Extracellular Vesicle Measurements"

Microtrac
Total Solutions in Particle Characterization

### - Particle Size Analysis -

C:\Microtrac\FLEX 11.1.0.1\Databases\Gonzalo\28.8.23.MDB

#### -Tabular Data -

| d(nm) | qr(%) | Qr(%P) | d(nm) | qr(%) | Qr(%P) |
|---|---|---|---|---|---|
| 6540 | 0,00 | 100,00 | 15,19 | 0,00 | 0,00 |
| 5500 | 0,00 | 100,00 | 12,77 | 0,00 | 0,00 |
| 4620 | 0,00 | 100,00 | 10,74 | 0,00 | 0,00 |
| 3890 | 0,00 | 100,00 | 9,03 | 0,00 | 0,00 |
| 3270 | 0,00 | 100,00 | 7,60 | 0,00 | 0,00 |
| 2750 | 0,00 | 100,00 | 6,39 | 0,00 | 0,00 |
| 2312 | 0,00 | 100,00 | 5,37 | 0,00 | 0,00 |
| 1944 | 0,00 | 100,00 | 4,52 | 0,00 | 0,00 |
| 1635 | 0,00 | 100,00 | 3,80 | 0,00 | 0,00 |
| 1375 | 0,00 | 100,00 | 3,19 | 0,00 | 0,00 |
| 1156 | 0,00 | 100,00 | 2,690 | 0,00 | 0,00 |
| 972,0 | 0,00 | 100,00 | 2,260 | 0,00 | 0,00 |
| 818,0 | 0,00 | 100,00 | 1,900 | 0,00 | 0,00 |
| 687,0 | 0,00 | 100,00 | 1,600 | 0,00 | 0,00 |
| 578,0 | 0,00 | 100,00 | 1,340 | 0,00 | 0,00 |
| 486,0 | 0,00 | 100,00 | 1,130 | 0,00 | 0,00 |
| 409,0 | 0,00 | 100,00 | 0,950 | 0,00 | 0,00 |
| 344,0 | 0,00 | 100,00 | | | |
| 289,0 | 0,00 | 100,00 | | | |
| 243,0 | 53,41 | 100,00 | | | |
| 204,4 | 46,59 | 46,59 | | | |
| 171,9 | 0,00 | 0,00 | | | |
| 144,5 | 0,00 | 0,00 | | | |
| 121,5 | 0,00 | 0,00 | | | |
| 102,2 | 0,00 | 0,00 | | | |
| 85,90 | 0,00 | 0,00 | | | |
| 72,30 | 0,00 | 0,00 | | | |
| 60,80 | 0,00 | 0,00 | | | |
| 51,10 | 0,00 | 0,00 | | | |
| 43,00 | 0,00 | 0,00 | | | |
| 36,10 | 0,00 | 0,00 | | | |
| 30,40 | 0,00 | 0,00 | | | |
| 25,55 | 0,00 | 0,00 | | | |
| 21,48 | 0,00 | 0,00 | | | |
| 18,06 | 0,00 | 0,00 | | | |

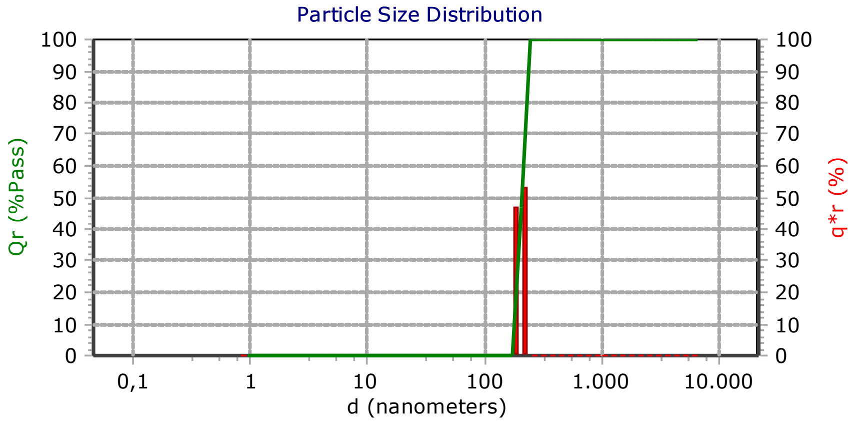


#### - Measurement Info -

| Title | |
|---|---|
| iso1 ht1080 f3 onh dilution | |
| **Identifiers** | |
| - | |
| - | |
| Database Record | 20 |
| Run Number | 2 of 3 |
| Date | 10.10.2024 |
| Time | 12:41 |
| Acquired Date | 10.10.2024 |
| Acquired Time | 12:41 |
| Serial Number | W3422 |
| **Calculated Data** | |
| Above Residual | 0 |
| Below Residual | 0 |
| Loading Index | 0,040 |
| Conc. Index | 0,02661 |
| RMS Residual | 4,043% |
| Cell Temp (C) | 22,47 |
| Viscosity(cp) | 0,9440 |
| Reflected Pwr (uW) | 2,90 |
| **User Defined Calculations** | |
| Name | Value |
| **Recalculation Status** | |
| DB-Meas : : Original : | |

#### -SOP Info-

| EV_ANNA(*) | |
|---|---|
| **Timing** | |
| Setzero Time | 30 (sec) |
| Run Time | 30 (sec) |
| Number of Runs | 3 |
| Multi-Run Delay | 0 (min) |
| Delay First Meas. | Disabled |
| **Analysis** | |
| Russ | |
| Refractive Index | N/A |
| Transparency | Absorb |
| Shape | Irregular |
| WATER | |
| Refractive Index | 1,33 |
| Low Temperature | 20,0 |
| Low Temp. Visc. | 1,002 |
| High Temperature | 30,0 |
| High Temp. Visc. | 0,797 |
| Options: | |
| Analysis Type | Mode |
| **Perspective** | |
| Progression | Standard |
| Distribution | Intensity |
| Upper Edge(nm) | 6540 |
| Lower Edge(nm) | 0,8 |
| Residuals | Disabled |

| Summary | |
|---|---|
| Data | Value |
| MI(nm): | 206,5 |
| MN(nm): | 201,6 |
| MA(nm): | 204,9 |
| CS: | 29,29 |
| SD: | 21,04 |
| PDI: | 0,0536 |
| Mz: | 206,5 |
| σι: | 19,63 |
| Ski: | 18,84 |
| Kg: | 820,8 |

| Percentiles | |
|---|---|
| d(i) | d(nm) |
| 10,00 | 181,4 |
| 20,00 | 188,2 |
| 30,00 | 194,5 |
| 40,00 | 200,5 |
| 50,00 | 206,4 |
| 60,00 | 212,1 |
| 70,00 | 218,3 |
| 80,00 | 224,8 |
| 90,00 | 231,9 |
| 95,00 | 237,2 |

Mode Summary

| d(nm) | Pct | Width | C(I) | C(V):cc/ml |
|---|---|---|---|---|
| 202,2 | 100,0 | 3,97E+01 | 3,97E-02 | 8,2E-08 |

10.10.2024 12:54

**FLEX**

11.1.0.1

#### - Notes -

-

Microtrac
Total Solutions in Particle Characterization

### - Particle Size Analysis -

C:\Microtrac\FLEX 11.1.0.1\Databases\Gonzalo\28.8.23.MDB

#### -Tabular Data -

| d(nm) | qr(%) | Qr(%P) | d(nm) | qr(%) | Qr(%P) |
|---|---|---|---|---|---|
| 6540 | 0,00 | 100,00 | 15,19 | 0,00 | 0,00 |
| 5500 | 0,00 | 100,00 | 12,77 | 0,00 | 0,00 |
| 4620 | 0,00 | 100,00 | 10,74 | 0,00 | 0,00 |
| 3890 | 0,00 | 100,00 | 9,03 | 0,00 | 0,00 |
| 3270 | 0,00 | 100,00 | 7,60 | 0,00 | 0,00 |
| 2750 | 0,00 | 100,00 | 6,39 | 0,00 | 0,00 |
| 2312 | 0,00 | 100,00 | 5,37 | 0,00 | 0,00 |
| 1944 | 0,00 | 100,00 | 4,52 | 0,00 | 0,00 |
| 1635 | 0,00 | 100,00 | 3,80 | 0,00 | 0,00 |
| 1375 | 0,00 | 100,00 | 3,19 | 0,00 | 0,00 |
| 1156 | 0,00 | 100,00 | 2,690 | 0,00 | 0,00 |
| 972,0 | 0,00 | 100,00 | 2,260 | 0,00 | 0,00 |
| 818,0 | 0,00 | 100,00 | 1,900 | 0,00 | 0,00 |
| 687,0 | 0,00 | 100,00 | 1,600 | 0,00 | 0,00 |
| 578,0 | 0,00 | 100,00 | 1,340 | 0,00 | 0,00 |
| 486,0 | 0,00 | 100,00 | 1,130 | 0,00 | 0,00 |
| 409,0 | 0,00 | 100,00 | 0,950 | 0,00 | 0,00 |
| 344,0 | 0,00 | 100,00 | | | |
| 289,0 | 0,00 | 100,00 | | | |
| 243,0 | 0,00 | 100,00 | | | |
| 204,4 | 0,00 | 100,00 | | | |
| 171,9 | 4,16 | 100,00 | | | |
| 144,5 | 87,14 | 95,84 | | | |
| 121,5 | 8,70 | 8,70 | | | |
| 102,2 | 0,00 | 0,00 | | | |
| 85,90 | 0,00 | 0,00 | | | |
| 72,30 | 0,00 | 0,00 | | | |
| 60,80 | 0,00 | 0,00 | | | |
| 51,10 | 0,00 | 0,00 | | | |
| 43,00 | 0,00 | 0,00 | | | |
| 36,10 | 0,00 | 0,00 | | | |
| 30,40 | 0,00 | 0,00 | | | |
| 25,55 | 0,00 | 0,00 | | | |
| 21,48 | 0,00 | 0,00 | | | |
| 18,06 | 0,00 | 0,00 | | | |

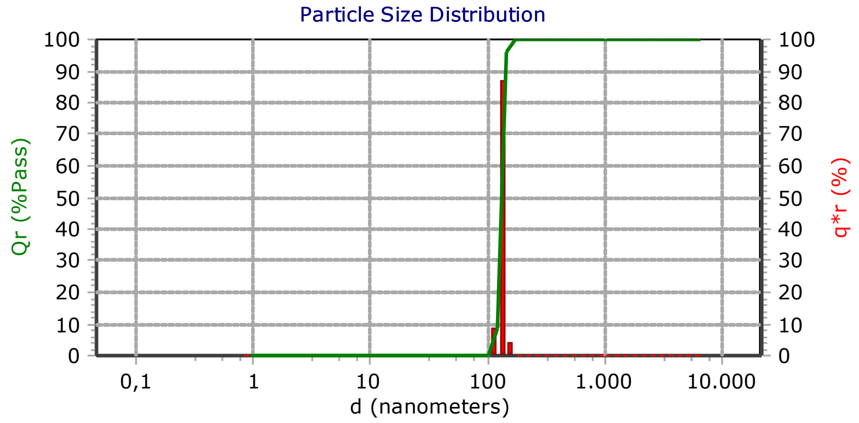


#### - Measurement Info -

| Title | |
|---|---|
| iso1 ht1080 f3 onh dilution | |
| Identifiers | |
| - | |
| - | |
| Database Record | 21 |
| Run Number | 3 of 3 |
| Date | 10.10.2024 |
| Time | 12:42 |
| Acquired Date | 10.10.2024 |
| Acquired Time | 12:42 |
| Serial Number | W3422 |
| Calculated Data | |
| Above Residual | 0 |
| Below Residual | 0 |
| Loading Index | 0,251 |
| Conc. Index | 0,1977 |
| RMS Residual | 4,552% |
| Cell Temp (C) | 22,53 |
| Viscosity(cp) | 0,9430 |
| Reflected Pwr (uW) | 2,90 |
| User Defined Calculations | |
| Name | Value |
| Recalculation Status | |
| DB-Meas : : Original : | |

#### -SOP Info-

| EV_ANNA(*) | |
|---|---|
| Timing | |
| Setzero Time | 30 (sec) |
| Run Time | 30 (sec) |
| Number of Runs | 3 |
| Multi-Run Delay | 0 (min) |
| Delay First Meas. | Disabled |
| Analysis | |
| Russ | |
| Refractive Index | N/A |
| Transparency | Absorb |
| Shape | Irregular |
| WATER | |
| Refractive Index | 1,33 |
| Low Temperature | 20,0 |
| Low Temp. Visc. | 1,002 |
| High Temperature | 30,0 |
| High Temp. Visc. | 0,797 |
| Options: | |
| Analysis Type | Mode |
| Perspective | |
| Progression | Standard |
| Distribution | Intensity |
| Upper Edge(nm) | 6540 |
| Lower Edge(nm) | 0,8 |
| Residuals | Disabled |

| Summary | |
|---|---|
| Data | Value |
| MI(nm): | 131,9 |
| MN(nm): | 130,5 |
| MA(nm): | 131,4 |
| CS: | 45,65 |
| SD: | 8,48 |
| PDI: | 0,0426 |
| Mz: | 132,0 |
| σι: | 8,11 |
| Ski: | -8,70945 |
| Kg: | 899,4 |

| Percentiles | |
|---|---|
| d(i) | d(nm) |
| 10,00 | 121,9 |
| 20,00 | 124,7 |
| 30,00 | 127,4 |
| 40,00 | 129,7 |
| 50,00 | 131,9 |
| 60,00 | 134,2 |
| 70,00 | 136,6 |
| 80,00 | 139,2 |
| 90,00 | 142,5 |
| 95,00 | 144,2 |

Mode Summary

| d(nm) | Pct | Width | C(I) | C(V):cc/ml |
|---|---|---|---|---|
| 128,7 | 100,0 | 2,51E+02 | 2,51E-01 | -2,33E-06 |

10.10.2024 12:57

FLEX

11.1.0.1

#### - Notes -

-

PARTICLE METRIX

ZETAVIEW®

www.particle-metrix.com

Electrophoresis & Brownian Motion
Video Analysis
Laser Scattering Microscopy

Video Operator: ZetaUser

Operator (Report): ZetaUser

**Sample Parameters**
Sample Name:
Sample Info 1:
Sample Info 2:
Sample Info 3:
Electrolyte: H2O
Temperature: 24.66 °C sensed
pH 7.0 entered
Conductivity: 1387.00 µS/cm sensed

**Instrument Parameters**
Laser Wavelength: 488 nm
Filter Wavelength: Scatter
Sensitivity: 75
Shutter: 100

**SOP: SOP100_ExTu**
Size Distribution 1 Cycle 11 Positions
Description:

**Result (sizes in nm)**

| | Number | Concentration | Volume |
|---|---|---|---|
| Median (X50) | 117.4 | 117.4 | 166.6 |
| StdDev | 44.2 | 44.1 | 79.0 |

Concentration: 2.5E+7 Particles / mL
Dilution Factor: 500
Original Concentration: 1.2E+10 Particles / mL

**Quality**
Average Counted Particles per Frame: 69
Number of Traced Particles: 945

**Analysis Parameters**
Max Area 1000, Min Area 10, Min Brightness 20, nm/Class: 5

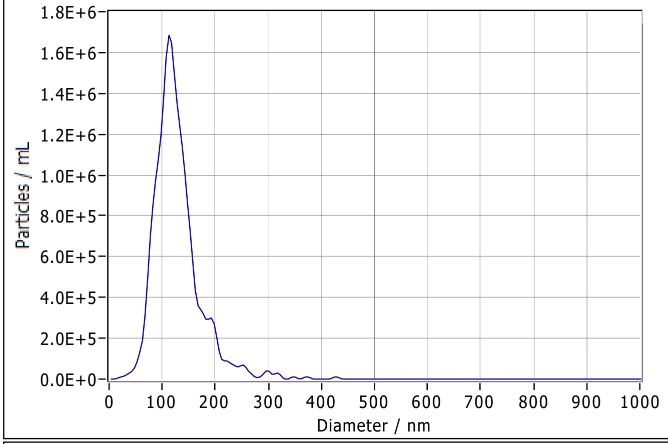


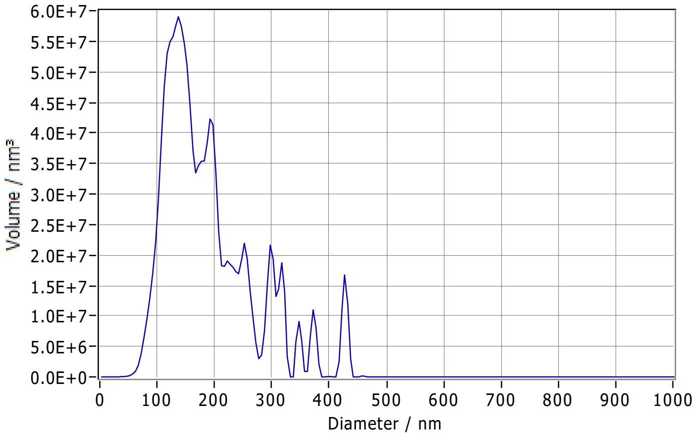


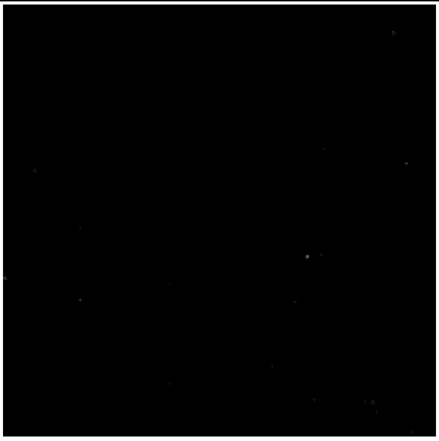

**Peak Analysis (Concentration)**

| Diameter / nm | Particles/mL | FWHM / nm | Percentage |
|---|---|---|---|
| 114.6 | 1.7E+6 | 66.0 | 100.0 |

**X Values** (all sizes are given in nm)

| | Number | Concentration | Volume |
|---|---|---|---|
| X10 | 79.7 | 79.7 | 107.8 |
| X50 | 117.4 | 117.4 | 166.6 |
| X90 | 179.8 | 179.8 | 301.3 |
| Span | 0.9 | 0.9 | 1.2 |
| Mean | 127.7 | 127.7 | 189.3 |
| StdDev | 44.2 | 44.1 | 79.0 |

**Comment**

(Signature)

Analyzed Video: C:\Users\ZetaView\Desktop\g\comparison\20241010_0001_HT1080_ISO1_F3_a_size_488.avi

Experiment: 2024-10-10 13:57 ZetaView S/N 20-559, Software ZetaView (version 8.05.16 SP3)

Report: 2024-10-10 14:07 Software ZetaView (version 8.05.16 SP3)

PARTICLE METRIX

### ZETAVIEW®

www.particle-metrix.com

Electrophoresis & Brownian Motion
Video Analysis
Laser Scattering Microscopy

Video Operator: ZetaUser

Operator (Report): ZetaUser

**Sample Parameters**
Sample Name:
Sample Info 1:
Sample Info 2:
Sample Info 3:
Electrolyte: H2O
Temperature: 25.05 °C sensed
pH 7.0 entered
Conductivity: 1310.00 µS/cm sensed

**Instrument Parameters**
Laser Wavelength: 488 nm
Filter Wavelength: Scatter
Sensitivity: 75
Shutter: 100

**SOP: SOP100_ExTu**
Size Distribution 1 Cycle 11 Positions
Description:

**Result (sizes in nm)**

| | Number | Concentration | Volume |
|---|---|---|---|
| Median (X50) | 125.6 | 125.6 | 188.3 |
| StdDev | 51.1 | 51.0 | 128.1 |

Concentration: 3.1E+7 Particles / mL
Dilution Factor: 500
Original Concentration: 1.6E+10 Particles / mL

**Quality**
Average Counted Particles per Frame: 86
Number of Traced Particles: 1146

**Analysis Parameters**
Max Area 1000, Min Area 10, Min Brightness 20, nm/Class: 5

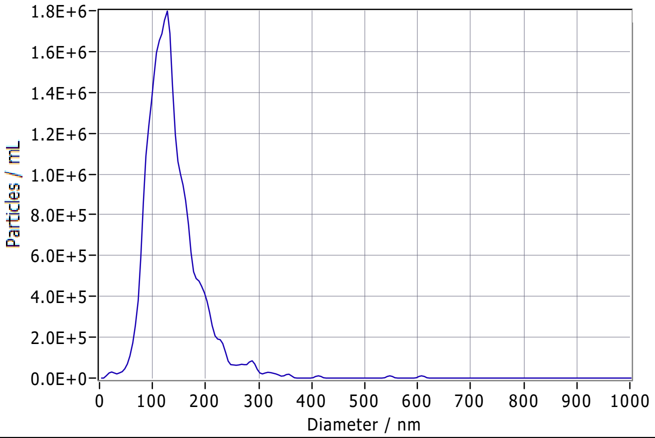


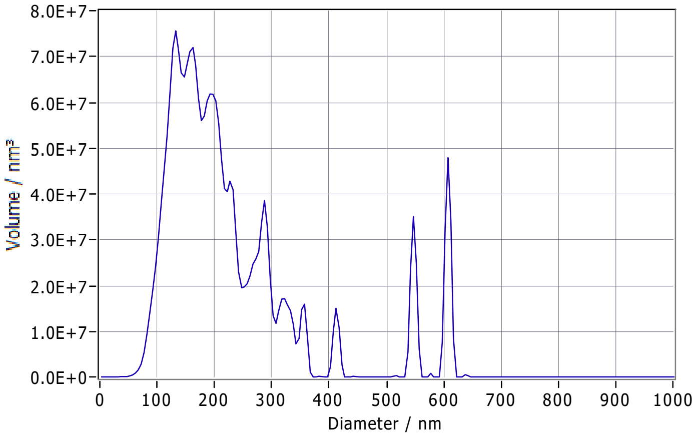


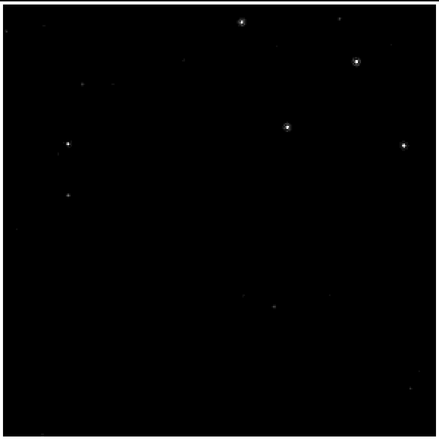

**Peak Analysis (Concentration)**

| Diameter / nm | Particles/mL | FWHM / nm | Percentage |
|---|---|---|---|
| 123.0 | 1.8E+6 | 77.2 | 100.0 |

**X Values** (all sizes are given in nm)

| | Number | Concentration | Volume |
|---|---|---|---|
| X10 | 84.5 | 84.5 | 116.6 |
| X50 | 125.6 | 125.6 | 188.3 |
| X90 | 195.9 | 195.9 | 407.9 |
| Span | 0.9 | 0.9 | 1.5 |
| Mean | 137.4 | 137.4 | 229.4 |
| StdDev | 51.1 | 51.0 | 128.1 |

**Comment**

(Signature)

Analyzed Video: C:\Users\ZetaView\Desktop\g\comparison\20241010_0002_HT1080_ISO1_F3_b_size_488.avi

Experiment: 2024-10-10 14:08 ZetaView S/N 20-559, Software ZetaView (version 8.05.16 SP3)

Report: 2024-10-10 14:09 Software ZetaView (version 8.05.16 SP3)

### e_id: 3455, ZetaScript: SOP100, Sample name: HT1080 ISO1 F3 dls nta comparison

Measurement created: 2024-10-10 14:12:02 Report created: 2024-10-10 14:17:34 User: ZetaUser

#### Measurement specification: SOP100(1)

##### Capture settings

| | | | |
|---|---|---|---|
| Temperature / °C | 25.0 | Laser wave length / nm | 488 |
| Dilution | 500 | Filter wave length / nm | 0 |
| Viscosity / mPa · s | 0.899 | Sensitivity | 80 |
| Positions | None | Shutter | 90 |
| Video length | 30 | | |
| Frame rate / 1/s | 30.303 | | |
| Experiment type | Size distribution | | |

##### Analysis settings

| | |
|---|---|
| Concentration correction | 0.55 |
| Concentration calibration | 636400 |
| Min size /nm | 10 |
| Max size / nm | 1000 |
| Min area / px | 1 |
| Max area / px | 1000 |
| Trace length | 15 |

Comment:

#### Graphics:

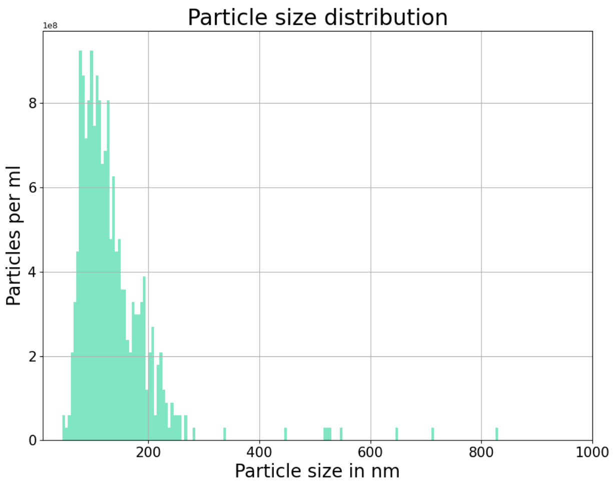


#### Results:

| | Peak /nm | Median /nm | #Particles | Avg. Num Det. | FWHM /nm | Concentration /1/mL | Position |
|---|---|---|---|---|---|---|---|
| | **106.6 (± 8.9)** | **121.2 (± 10.2)** | **559** | **95.4 (± 15.8)** | **102.9 (± 23.0)** | **1.67E+10 (± 2.8E+09)** | |
| 1 | 120.2 | 140.8 | 66 | 74.6 | 143.3 | 1.31E+10 | 0.30 |
| 2 | 111.8 | 127.4 | 81 | 98.2 | 132.5 | 1.72E+10 | 0.40 |
| 3 | 98.3 | 115.8 | 90 | 101.4 | 88.8 | 1.77E+10 | 0.50 |
| 4 | 93.2 | 106.6 | 100 | 128.3 | 93.0 | 2.25E+10 | 0.60 |
| 5 | 109.3 | 119.5 | 80 | 86.3 | 98.2 | 1.51E+10 | 0.70 |
| 6 | 100.1 | 114.0 | 74 | 93.7 | 85.3 | 1.64E+10 | 0.80 |
| 7 | 113.2 | 124.1 | 68 | 85.1 | 79.0 | 1.49E+10 | 0.85 |

### e_id: 3455, ZetaScript: SOP100, Sample name: HT1080 ISO1 F3 dls nta comparison

Measurement created: 2024-10-10 14:12:02 Report created: 2024-10-10 14:17:34 User: ZetaUser

#### Measurement specification: SOP100(2)

##### Capture settings

| | | | |
|---|---|---|---|
| Temperature / °C | 25.0 | Laser wave length / nm | 488 |
| Dilution | 500 | Filter wave length / nm | 0 |
| Viscosity / mPa · s | 0.899 | Sensitivity | 80 |
| Positions | None | Shutter | 90 |
| Video length | 30 | | |
| Frame rate / 1/s | 30.303 | | |
| Experiment type | Size distribution | | |

##### Analysis settings

| | |
|---|---|
| Concentration correction | 0.55 |
| Concentration calibration | 636400 |
| Min size /nm | 10 |
| Max size / nm | 1000 |
| Min area / px | 1 |
| Max area / px | 1000 |
| Trace length | 15 |

Comment:

#### Graphics:

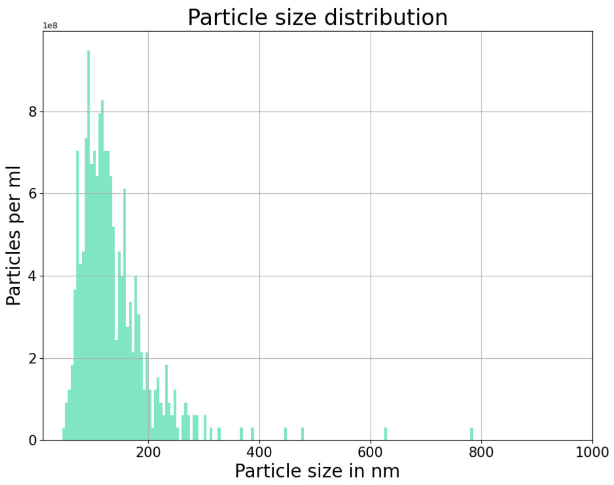


#### Results:

| | Peak /nm | Median /nm | #Particles | Avg. Num Det. | FWHM /nm | Concentration /1/mL | Position |
|---|---|---|---|---|---|---|---|
| | **110.0 (± 8.2)** | **121.6 (± 8.3)** | **516** | **90.2 (± 10.8)** | **102.0 (± 18.6)** | **1.58E+10 (± 1.9E+09)** | |
| 1 | 107.7 | 121.0 | 64 | 74.5 | 89.6 | 1.30E+10 | 0.30 |
| 2 | 107.9 | 121.2 | 79 | 97.8 | 101.6 | 1.71E+10 | 0.40 |
| 3 | 115.9 | 128.2 | 83 | 92.1 | 118.0 | 1.61E+10 | 0.50 |
| 4 | 105.3 | 113.7 | 75 | 109.0 | 81.6 | 1.91E+10 | 0.60 |
| 5 | 111.6 | 131.9 | 69 | 81.0 | 127.1 | 1.42E+10 | 0.70 |
| 6 | 96.5 | 106.7 | 79 | 94.3 | 76.3 | 1.65E+10 | 0.80 |
| 7 | 124.9 | 128.4 | 67 | 82.7 | 119.7 | 1.45E+10 | 0.85 |

### e_id: 2764, ZetaScript: SM50-100, Sample name: HT29 after italy nd try

Measurement created: 2024-06-07 16:41:40 Report created: 2025-07-16 10:24:56 User: ZetaUser

#### Measurement specification: SOP100(1)

##### Capture settings

| | | | |
|---|---|---|---|
| Temperature / °C | 25.2 | Laser wave length / nm | 488 |
| Dilution | 100 | Filter wave length / nm | 0 |
| Viscosity / mPa · s | 0.895 | Sensitivity | 80 |
| Positions | None | Shutter | 90 |
| Video length | 30 | | |
| Frame rate / 1/s | 30.303 | | |
| Experiment type | Size distribution | | |

##### Analysis settings

| | |
|---|---|
| Concentration correction | 0.55 |
| Concentration calibration | 636400 |
| Min size /nm | 10 |
| Max size / nm | 1000 |
| Min area / px | 1 |
| Max area / px | 1000 |
| Trace length | 15 |

Comment:

#### Graphics:

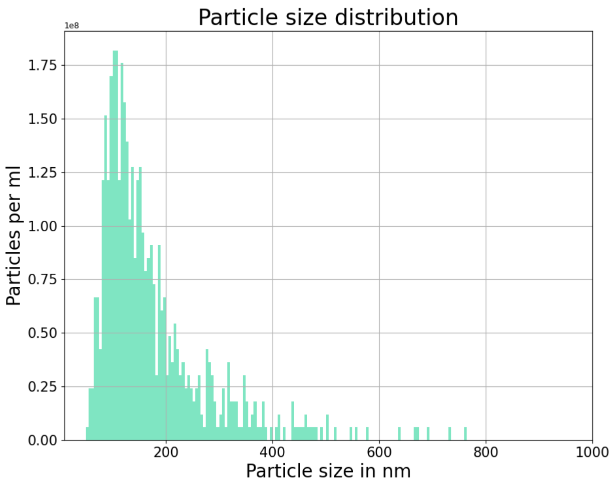


#### Results:

| | Peak /nm | Median /nm | #Particles | Avg. Num Det. | FWHM /nm | Concentration /1/mL | Position |
|---|---|---|---|---|---|---|---|
| | **118.1 (± 10.0)** | **143.1 (± 10.9)** | **664** | **115.0 (± 8.5)** | **132.5 (± 22.9)** | **4.02E+09 (± 3.0E+08)** | |
| 1 | 136.1 | 156.8 | 96 | 121.7 | 136.6 | 4.26E+09 | 0.30 |
| 2 | 112.0 | 135.4 | 108 | 121.0 | 117.0 | 4.24E+09 | 0.40 |
| 3 | 109.4 | 128.7 | 83 | 106.0 | 96.5 | 3.71E+09 | 0.50 |
| 4 | 123.8 | 149.9 | 113 | 128.5 | 134.0 | 4.50E+09 | 0.60 |
| 5 | 124.2 | 150.7 | 88 | 110.5 | 146.8 | 3.87E+09 | 0.70 |
| 6 | 104.2 | 128.8 | 93 | 113.5 | 121.8 | 3.97E+09 | 0.80 |
| 7 | 116.7 | 151.6 | 83 | 103.5 | 174.8 | 3.62E+09 | 0.85 |

### e_id: 2764, ZetaScript: SM50-100, Sample name: HT29 after italy nd try

Measurement created: 2024-06-07 16:41:40 Report created: 2025-07-16 10:24:57 User: ZetaUser

#### Measurement specification: SOP100(2)

##### Capture settings

| | | | |
|---|---|---|---|
| Temperature / °C | 25.1 | Laser wave length / nm | 488 |
| Dilution | 100 | Filter wave length / nm | 0 |
| Viscosity / mPa · s | 0.897 | Sensitivity | 80 |
| Positions | None | Shutter | 90 |
| Video length | 30 | | |
| Frame rate / 1/s | 30.303 | | |
| Experiment type | Size distribution | | |

##### Analysis settings

| | |
|---|---|
| Concentration correction | 0.55 |
| Concentration calibration | 636400 |
| Min size /nm | 10 |
| Max size / nm | 1000 |
| Min area / px | 1 |
| Max area / px | 1000 |
| Trace length | 15 |

Comment:

#### Graphics:

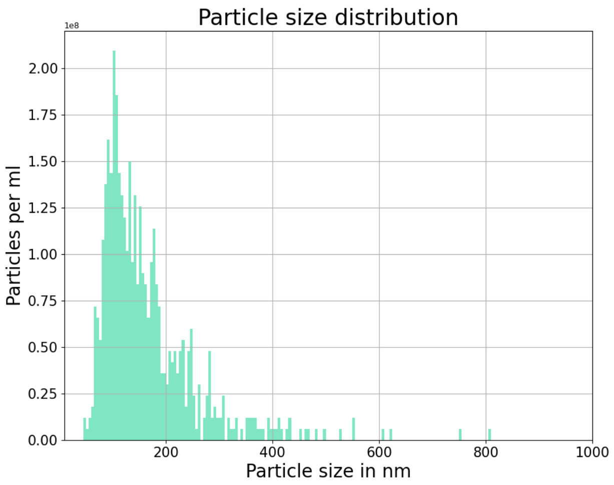


#### Results:

| | Peak /nm | Median /nm | #Particles | Avg. Num Det. | FWHM /nm | Concentration /1/mL | Position |
|---|---|---|---|---|---|---|---|
| | **121.0 (± 13.0)** | **140.5 (± 12.0)** | **642** | **109.9 (± 9.4)** | **126.7 (± 17.7)** | **3.85E+09 (± 3.3E+08)** | |
| 1 | 104.4 | 117.5 | 103 | 126.7 | 95.9 | 4.44E+09 | 0.30 |
| 2 | 119.4 | 146.1 | 98 | 112.9 | 145.8 | 3.95E+09 | 0.40 |
| 3 | 145.9 | 157.6 | 91 | 101.2 | 137.9 | 3.54E+09 | 0.50 |
| 4 | 133.3 | 149.5 | 89 | 111.1 | 133.7 | 3.89E+09 | 0.60 |
| 5 | 116.0 | 133.0 | 81 | 97.4 | 104.1 | 3.41E+09 | 0.70 |
| 6 | 113.9 | 137.6 | 89 | 103.4 | 128.6 | 3.62E+09 | 0.80 |
| 7 | 113.8 | 142.0 | 91 | 116.5 | 140.7 | 4.08E+09 | 0.85 |

#### Size & Concentration Report

ht29 unstained new

| Data File | 20240607 ht29 unstained new 26.nfa | Population | Total |
|---|---|---|---|

SN: FNAU30E23122748
Software: V2.0
Sample Pressure: 1.0Kpa

Laser: 6/50 mW 488
SS Decay: 10%
Threshold/sub: 58.9 7.9 5.5 1/0 0 0 0
Min Width: 0.3 ms

##### Total Size Information

| | | |
|---|---|---|
| All Events | 837 | |
| Gating Events | 837 | |
| % of all | 100.00 | |
| Median | 61.8 | nm |
| Mean | 66.0 | nm |
| Std Dev. | 15.7 | nm |

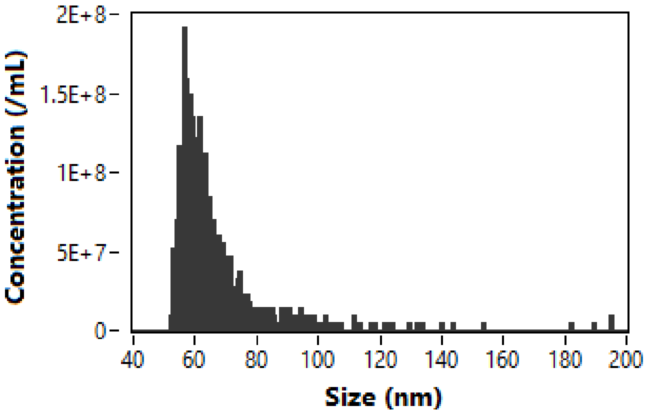


##### Total Concentration Information

| | Particle Number | Dilution Factor |
|---|---|---|
| STD | 4591 | 100 |
| Blank | 0 | —— |
| Sample | 837 | 100 |

| | | |
|---|---|---|
| STD Con. | 2.14E+10 | Particles/mL |
| Sample Flow Rate | 21.45 | nL/min |
| Sample Con. | 3.90E+9 | Particles/mL |

Corrected Ratio: 837/837 100.0%

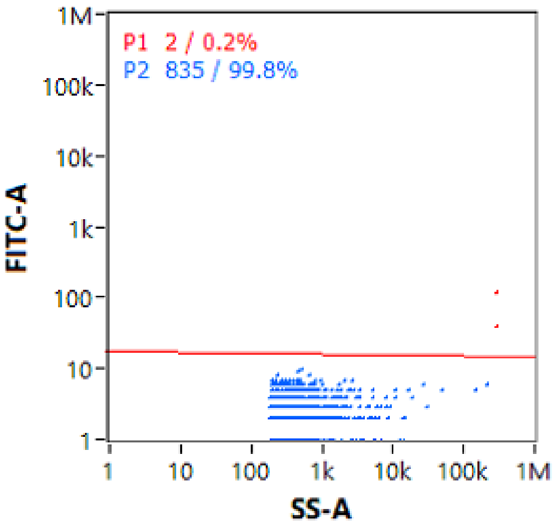


Report By :

7/16/2025 12:35 PM
