## Supplementary Material 3 for "PHoNUPS: Open-Source Software for Standardized Analysis and Visualization of Multi-Instrument Extracellular Vesicle Measurements"

### - Particle Size Analysis -

C:\Microtrac\FLEX 11.1.0.1\Databases\Gonzalo\28.8.23.MDB

#### -Tabular Data -

| d(nm) | qr(%) | Qr(%P) | d(nm) | qr(%) | Qr(%P) |
|---|---|---|---|---|---|
| 6540 | 0,00 | 100,00 | 15,19 | 0,00 | 0,00 |
| 5500 | 0,00 | 100,00 | 12,77 | 0,00 | 0,00 |
| 4620 | 0,00 | 100,00 | 10,74 | 0,00 | 0,00 |
| 3890 | 0,00 | 100,00 | 9,03 | 0,00 | 0,00 |
| 3270 | 0,00 | 100,00 | 7,60 | 0,00 | 0,00 |
| 2750 | 0,00 | 100,00 | 6,39 | 0,00 | 0,00 |
| 2312 | 1,71 | 100,00 | 5,37 | 0,00 | 0,00 |
| 1944 | 4,08 | 98,29 | 4,52 | 0,00 | 0,00 |
| 1635 | 3,54 | 94,21 | 3,80 | 0,00 | 0,00 |
| 1375 | 1,32 | 90,67 | 3,19 | 0,00 | 0,00 |
| 1156 | 0,10 | 89,35 | 2,690 | 0,00 | 0,00 |
| 972,0 | 0,00 | 89,25 | 2,260 | 0,00 | 0,00 |
| 818,0 | 0,18 | 89,25 | 1,900 | 0,00 | 0,00 |
| 687,0 | 0,47 | 89,07 | 1,600 | 0,00 | 0,00 |
| 578,0 | 4,68 | 88,60 | 1,340 | 0,00 | 0,00 |
| 486,0 | 4,48 | 83,92 | 1,130 | 0,00 | 0,00 |
| 409,0 | 4,32 | 79,44 | 0,950 | 0,00 | 0,00 |
| 344,0 | 6,91 | 75,12 | | | |
| 289,0 | 9,43 | 68,21 | | | |
| 243,0 | 11,20 | 58,78 | | | |
| 204,4 | 11,68 | 47,58 | | | |
| 171,9 | 10,45 | 35,90 | | | |
| 144,5 | 8,25 | 25,45 | | | |
| 121,5 | 5,77 | 17,20 | | | |
| 102,2 | 3,68 | 11,43 | | | |
| 85,90 | 2,29 | 7,75 | | | |
| 72,30 | 1,53 | 5,46 | | | |
| 60,80 | 1,16 | 3,93 | | | |
| 51,10 | 0,95 | 2,77 | | | |
| 43,00 | 0,75 | 1,82 | | | |
| 36,10 | 0,54 | 1,07 | | | |
| 30,40 | 0,34 | 0,53 | | | |
| 25,55 | 0,19 | 0,19 | | | |
| 21,48 | 0,00 | 0,00 | | | |
| 18,06 | 0,00 | 0,00 | | | |

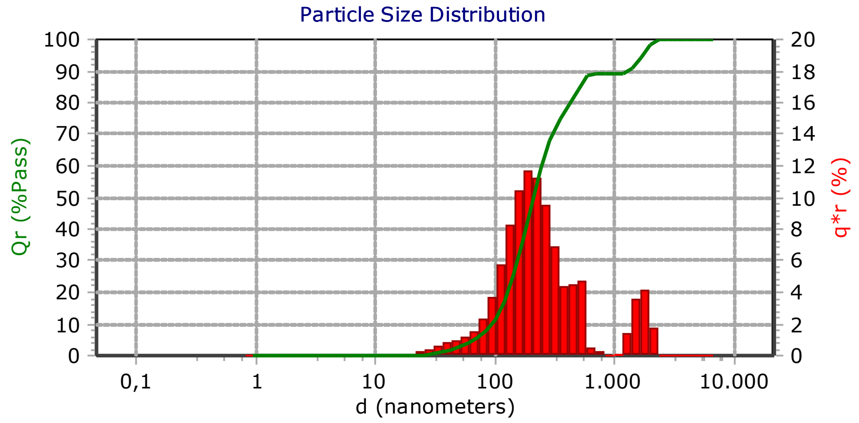


#### - Measurement Info -

| Title | |
|---|---|
| f1_GBQ_serum_onhe_label_1 | |
| Identifiers | |
| - | |
| - | |
| Database Record | 77 |
| Run Number | 2 of 3 |
| Date | 28.10.2024 |
| Time | 11:34 |
| Acquired Date | 28.10.2024 |
| Acquired Time | 11:34 |
| Serial Number | W3422 |
| Calculated Data | |
| Above Residual | 0 |
| Below Residual | 0 |
| Loading Index | 0,223 |
| Conc. Index | 0,1788 |
| RMS Residual | 0,479% |
| Cell Temp (C) | 22,58 |
| Viscosity(cp) | 0,9420 |
| Reflected Pwr (uW) | 2,80 |
| User Defined Calculations | |
| Name | Value |
| Recalculation Status | |
| DB-Meas : : Original : | |

#### -SOP Info-

| EV_ANNA(*) | |
|---|---|
| Timing | |
| Setzero Time | 30 (sec) |
| Run Time | 30 (sec) |
| Number of Runs | 3 |
| Multi-Run Delay | 0 (min) |
| Delay First Meas. | Disabled |
| Analysis | |
| Russ | |
| Refractive Index | N/A |
| Transparency | Absorb |
| Shape | Irregular |
| WATER | |
| Refractive Index | 1,33 |
| Low Temperature | 20,0 |
| Low Temp. Visc. | 1,002 |
| High Temperature | 30,0 |
| High Temp. Visc. | 0,797 |
| Options: | |
| Analysis Type | Mode |
| Perspective | |
| Progression | Standard |
| Distribution | Intensity |
| Upper Edge(nm) | 6540 |
| Lower Edge(nm) | 0,8 |
| Residuals | Disabled |

| Summary | |
|---|---|
| Data | Value |
| MI(nm): | 380,0 |
| MN(nm): | 58,00 |
| MA(nm): | 173,6 |
| CS: | 34,56 |
| SD: | 184,7 |
| PDI: | 11,02 |
| Mz: | 272,3 |
| σι: | 337,5 |
| Ski: | 657,1 |
| Kg: | 3324 |

| Percentiles | |
|---|---|
| d(i) | d(nm) |
| 10,00 | 96,40 |
| 20,00 | 129,6 |
| 30,00 | 156,4 |
| 40,00 | 182,8 |
| 50,00 | 211,9 |
| 60,00 | 248,1 |
| 70,00 | 301,0 |
| 80,00 | 419,0 |
| 90,00 | 1294 |
| 95,00 | 1687 |

| Mode Summary | | | | |
|---|---|---|---|---|
| d(nm) | Pct | Width | C(I) | C(V):cc/ml |
| 51,81 | 5,89 | 1,32E+01 | 1,32E-02 | 3,17E-08 |
| 192,9 | 77,93 | 1,74E+02 | 1,74E-01 | 2,32E-07 |
| 493,0 | 5,43 | 1,21E+01 | 1,21E-02 | 1,98E-08 |
| 1.635 | 10,74 | 2,4E+01 | 2,4E-02 | 7,61E-08 |

16.01.2026 10:47

FLEX

11.1.0.1

#### - Notes -

-

Microtrac
Total Solutions in Particle Characterization

### - Particle Size Analysis -

C:\Microtrac\FLEX 11.1.0.1\Databases\Gonzalo\28.8.23.MDB

#### -Tabular Data -

| d(nm) | qr(%) | Qr(%P) | d(nm) | qr(%) | Qr(%P) |
|---|---|---|---|---|---|
| 6540 | 0,00 | 100,00 | 15,19 | 0,00 | 0,00 |
| 5500 | 0,00 | 100,00 | 12,77 | 0,00 | 0,00 |
| 4620 | 0,00 | 100,00 | 10,74 | 0,00 | 0,00 |
| 3890 | 0,00 | 100,00 | 9,03 | 0,00 | 0,00 |
| 3270 | 0,00 | 100,00 | 7,60 | 0,00 | 0,00 |
| 2750 | 0,00 | 100,00 | 6,39 | 0,00 | 0,00 |
| 2312 | 0,00 | 100,00 | 5,37 | 0,00 | 0,00 |
| 1944 | 0,00 | 100,00 | 4,52 | 0,00 | 0,00 |
| 1635 | 0,00 | 100,00 | 3,80 | 0,00 | 0,00 |
| 1375 | 0,00 | 100,00 | 3,19 | 0,00 | 0,00 |
| 1156 | 0,00 | 100,00 | 2,690 | 0,00 | 0,00 |
| 972,0 | 0,00 | 100,00 | 2,260 | 0,00 | 0,00 |
| 818,0 | 0,00 | 100,00 | 1,900 | 0,00 | 0,00 |
| 687,0 | 0,00 | 100,00 | 1,600 | 0,00 | 0,00 |
| 578,0 | 0,00 | 100,00 | 1,340 | 0,00 | 0,00 |
| 486,0 | 0,00 | 100,00 | 1,130 | 0,00 | 0,00 |
| 409,0 | 0,00 | 100,00 | 0,950 | 0,00 | 0,00 |
| 344,0 | 0,00 | 100,00 | | | |
| 289,0 | 0,00 | 100,00 | | | |
| 243,0 | 0,00 | 100,00 | | | |
| 204,4 | 3,73 | 100,00 | | | |
| 171,9 | 69,39 | 96,27 | | | |
| 144,5 | 26,88 | 26,88 | | | |
| 121,5 | 0,00 | 0,00 | | | |
| 102,2 | 0,00 | 0,00 | | | |
| 85,90 | 0,00 | 0,00 | | | |
| 72,30 | 0,00 | 0,00 | | | |
| 60,80 | 0,00 | 0,00 | | | |
| 51,10 | 0,00 | 0,00 | | | |
| 43,00 | 0,00 | 0,00 | | | |
| 36,10 | 0,00 | 0,00 | | | |
| 30,40 | 0,00 | 0,00 | | | |
| 25,55 | 0,00 | 0,00 | | | |
| 21,48 | 0,00 | 0,00 | | | |
| 18,06 | 0,00 | 0,00 | | | |

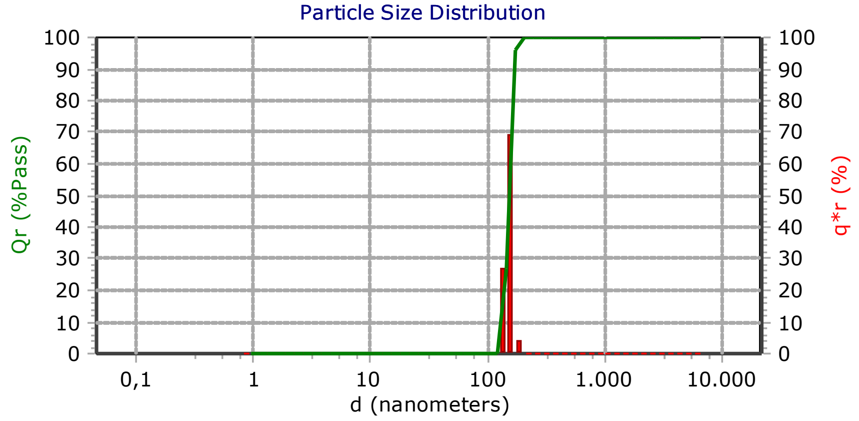


#### - Measurement Info -

| Title | |
|---|---|
| f1_GBQ_serum_onhe_label_1 | |
| Identifiers | |
| - | |
| - | |
| Database Record | 76 |
| Run Number | 1 of 3 |
| Date | 28.10.2024 |
| Time | 11:33 |
| Acquired Date | 28.10.2024 |
| Acquired Time | 11:33 |
| Serial Number | W3422 |
| Calculated Data | |
| Above Residual | 0 |
| Below Residual | 0 |
| Loading Index | 0,433 |
| Conc. Index | 0,347 |
| RMS Residual | 2,416% |
| Cell Temp (C) | 22,56 |
| Viscosity(cp) | 0,9420 |
| Reflected Pwr (uW) | 2,80 |
| User Defined Calculations | |
| Name | Value |
| Recalculation Status | |
| DB-Meas : : Original : | |

#### -SOP Info-

| EV_ANNA(*) | |
|---|---|
| Timing | |
| Setzero Time | 30 (sec) |
| Run Time | 30 (sec) |
| Number of Runs | 3 |
| Multi-Run Delay | 0 (min) |
| Delay First Meas. | Disabled |
| Analysis | |
| Russ | |
| Refractive Index | N/A |
| Transparency | Absorb |
| Shape | Irregular |
| WATER | |
| Refractive Index | 1,33 |
| Low Temperature | 20,0 |
| Low Temp. Visc. | 1,002 |
| High Temperature | 30,0 |
| High Temp. Visc. | 0,797 |
| Options: | |
| Analysis Type | Mode |
| Perspective | |
| Progression | Standard |
| Distribution | Intensity |
| Upper Edge(nm) | 6540 |
| Lower Edge(nm) | 0,8 |
| Residuals | Disabled |

| Summary | |
|---|---|
| Data | Value |
| MI(nm): | 152,1 |
| MN(nm): | 148,8 |
| MA(nm): | 151,0 |
| CS: | 39,73 |
| SD: | 13,36 |
| PDI: | 0,887 |
| Mz: | 152,3 |
| σI: | 13,06 |
| Ski: | -101,40493 |
| Kg: | 944,2 |

| Percentiles | |
|---|---|
| d(i) | d(nm) |
| 10,00 | 133,9 |
| 20,00 | 140,8 |
| 30,00 | 145,7 |
| 40,00 | 149,4 |
| 50,00 | 152,9 |
| 60,00 | 156,3 |
| 70,00 | 159,9 |
| 80,00 | 163,6 |
| 90,00 | 168,5 |
| 95,00 | 171,2 |

| Mode Summary | | | | |
|---|---|---|---|---|
| d(nm) | Pct | Width | C(l) | C(V):cc/ml |
| 147,8 | 100,0 | 4,33E+02 | 4,33E-01 | 2,75E-07 |

16.01.2026 10:46

FLEX
11.1.0.1

#### - Notes -

-

### - Particle Size Analysis -

C:\Microtrac\FLEX 11.1.0.1\Databases\Gonzalo\28.8.23.MDB

#### -Tabular Data -

| d(nm) | qr(%) | Qr(%P) | d(nm) | qr(%) | Qr(%P) |
|---|---|---|---|---|---|
| 6540 | 0,00 | 100,00 | 15,19 | 1,16 | 2,87 |
| 5500 | 0,00 | 100,00 | 12,77 | 0,46 | 1,71 |
| 4620 | 0,00 | 100,00 | 10,74 | 0,15 | 1,25 |
| 3890 | 0,00 | 100,00 | 9,03 | 0,13 | 1,10 |
| 3270 | 0,00 | 100,00 | 7,60 | 0,80 | 0,97 |
| 2750 | 0,00 | 100,00 | 6,39 | 0,17 | 0,17 |
| 2312 | 0,00 | 100,00 | 5,37 | 0,00 | 0,00 |
| 1944 | 0,00 | 100,00 | 4,52 | 0,00 | 0,00 |
| 1635 | 0,00 | 100,00 | 3,80 | 0,00 | 0,00 |
| 1375 | 0,00 | 100,00 | 3,19 | 0,00 | 0,00 |
| 1156 | 0,00 | 100,00 | 2,690 | 0,00 | 0,00 |
| 972,0 | 0,00 | 100,00 | 2,260 | 0,00 | 0,00 |
| 818,0 | 0,00 | 100,00 | 1,900 | 0,00 | 0,00 |
| 687,0 | 0,00 | 100,00 | 1,600 | 0,00 | 0,00 |
| 578,0 | 0,00 | 100,00 | 1,340 | 0,00 | 0,00 |
| 486,0 | 0,00 | 100,00 | 1,130 | 0,00 | 0,00 |
| 409,0 | 0,00 | 100,00 | 0,950 | 0,00 | 0,00 |
| 344,0 | 0,00 | 100,00 | | | |
| 289,0 | 0,00 | 100,00 | | | |
| 243,0 | 0,00 | 100,00 | | | |
| 204,4 | 0,42 | 100,00 | | | |
| 171,9 | 2,67 | 99,58 | | | |
| 144,5 | 8,89 | 96,91 | | | |
| 121,5 | 15,83 | 88,02 | | | |
| 102,2 | 14,86 | 72,19 | | | |
| 85,90 | 7,84 | 57,33 | | | |
| 72,30 | 3,29 | 49,49 | | | |
| 60,80 | 3,08 | 46,20 | | | |
| 51,10 | 4,78 | 43,12 | | | |
| 43,00 | 6,72 | 38,34 | | | |
| 36,10 | 7,90 | 31,62 | | | |
| 30,40 | 7,75 | 23,72 | | | |
| 25,55 | 6,33 | 15,97 | | | |
| 21,48 | 4,32 | 9,64 | | | |
| 18,06 | 2,45 | 5,32 | | | |

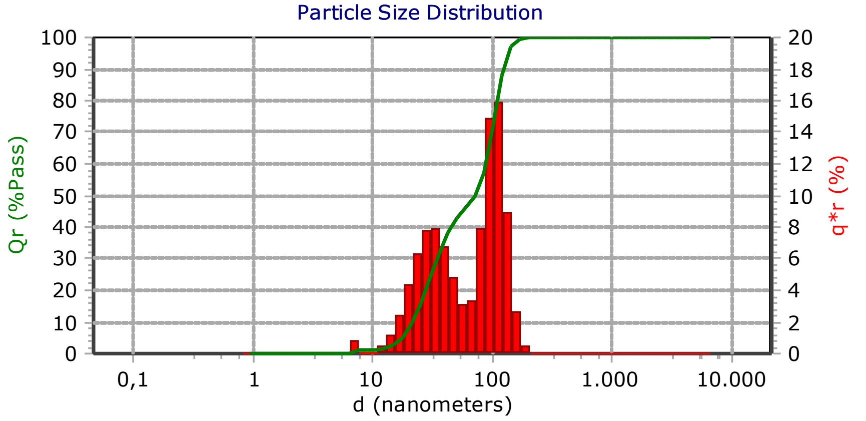


#### - Measurement Info -

| Title | |
|---|---|
| f5_GBQ_serum_onhe_label_1 | |
| Identifiers | |
| - | |
| - | |
| Database Record | 92 |
| Run Number | 1 of 3 |
| Date | 28.10.2024 |
| Time | 11:55 |
| Acquired Date | 28.10.2024 |
| Acquired Time | 11:55 |
| Serial Number | W3422 |
| Calculated Data | |
| Above Residual | 0 |
| Below Residual | 0 |
| Loading Index | 0,062 |
| Conc. Index | 0,0493 |
| RMS Residual | 0,147% |
| Cell Temp (C) | 21,87 |
| Viscosity(cp) | 0,9580 |
| Reflected Pwr (uW) | 2,80 |
| User Defined Calculations | |
| Name | Value |
| Recalculation Status | |
| DB-Meas : : Original : | |

#### -SOP Info-

| EV_ANNA(*) | |
|---|---|
| Timing | |
| Setzero Time | 30 (sec) |
| Run Time | 30 (sec) |
| Number of Runs | 3 |
| Multi-Run Delay | 0 (min) |
| Delay First Meas. | Disabled |
| Analysis | |
| Russ | |
| Refractive Index | N/A |
| Transparency | Absorb |
| Shape | Irregular |
| WATER | |
| Refractive Index | 1,33 |
| Low Temperature | 20,0 |
| Low Temp. Visc. | 1,002 |
| High Temperature | 30,0 |
| High Temp. Visc. | 0,797 |
| Options: | |
| Analysis Type | Mode |
| Perspective | |
| Progression | Standard |
| Distribution | Intensity |
| Upper Edge(nm) | 6540 |
| Lower Edge(nm) | 0,8 |
| Residuals | Disabled |

| Summary | |
|---|---|
| Data | Value |
| MI(nm): | 71,00 |
| MN(nm): | 14,35 |
| MA(nm): | 43,20 |
| CS: | 138,8 |
| SD: | 45,10 |
| PDI: | 7,43 |
| Mz: | 71,61 |
| σι: | 40,66 |
| Ski: | 2,580 |
| Kg: | 662,3 |

| Percentiles | |
|---|---|
| d(i) | d(nm) |
| 10,00 | 21,72 |
| 20,00 | 28,02 |
| 30,00 | 34,80 |
| 40,00 | 45,30 |
| 50,00 | 73,50 |
| 60,00 | 89,10 |
| 70,00 | 99,90 |
| 80,00 | 110,8 |
| 90,00 | 125,1 |
| 95,00 | 137,3 |

| Mode Summary | | | | |
|---|---|---|---|---|
| d(nm) | Pct | Width | C(I) | C(V):cc/ml |
| 6,78 | 1,067 | 6,6E-01 | 6,6E-04 | 4,84E-07 |
| 30,44 | 47,14 | 2,92E+01 | 2,92E-02 | 2,5E-07 |
| 102,3 | 51,79 | 3,2E+01 | 3,2E-02 | 4,41E-08 |

16.01.2026 10:49

**FLEX**

**11.1.0.1**

#### - Notes -

-

Microtrac
Total Solutions in Particle Characterization

### - Particle Size Analysis -

C:\Microtrac\FLEX 11.1.0.1\Databases\Gonzalo\28.8.23.MDB

#### -Tabular Data -

| d(nm) | qr(%) | Qr(%P) | d(nm) | qr(%) | Qr(%P) |
|---|---|---|---|---|---|
| 6540 | 0,00 | 100,00 | 15,19 | 3,11 | 5,29 |
| 5500 | 0,00 | 100,00 | 12,77 | 1,58 | 2,18 |
| 4620 | 0,00 | 100,00 | 10,74 | 0,50 | 0,60 |
| 3890 | 0,00 | 100,00 | 9,03 | 0,10 | 0,10 |
| 3270 | 1,59 | 100,00 | 7,60 | 0,00 | 0,00 |
| 2750 | 4,47 | 98,41 | 6,39 | 0,00 | 0,00 |
| 2312 | 4,22 | 93,94 | 5,37 | 0,00 | 0,00 |
| 1944 | 1,54 | 89,72 | 4,52 | 0,00 | 0,00 |
| 1635 | 0,12 | 88,18 | 3,80 | 0,00 | 0,00 |
| 1375 | 0,00 | 88,06 | 3,19 | 0,00 | 0,00 |
| 1156 | 0,00 | 88,06 | 2,690 | 0,00 | 0,00 |
| 972,0 | 0,00 | 88,06 | 2,260 | 0,00 | 0,00 |
| 818,0 | 0,00 | 88,06 | 1,900 | 0,00 | 0,00 |
| 687,0 | 0,00 | 88,06 | 1,600 | 0,00 | 0,00 |
| 578,0 | 0,10 | 88,06 | 1,340 | 0,00 | 0,00 |
| 486,0 | 0,25 | 87,96 | 1,130 | 0,00 | 0,00 |
| 409,0 | 0,55 | 87,71 | 0,950 | 0,00 | 0,00 |
| 344,0 | 1,05 | 87,16 | | | |
| 289,0 | 1,75 | 86,11 | | | |
| 243,0 | 2,58 | 84,36 | | | |
| 204,4 | 3,38 | 81,78 | | | |
| 171,9 | 4,06 | 78,40 | | | |
| 144,5 | 4,65 | 74,34 | | | |
| 121,5 | 5,32 | 69,69 | | | |
| 102,2 | 6,21 | 64,37 | | | |
| 85,90 | 7,20 | 58,16 | | | |
| 72,30 | 7,94 | 50,96 | | | |
| 60,80 | 8,12 | 43,02 | | | |
| 51,10 | 7,18 | 34,90 | | | |
| 43,00 | 5,68 | 27,72 | | | |
| 36,10 | 4,00 | 22,04 | | | |
| 30,40 | 2,78 | 18,04 | | | |
| 25,55 | 2,62 | 15,26 | | | |
| 21,48 | 3,42 | 12,64 | | | |
| 18,06 | 3,93 | 9,22 | | | |

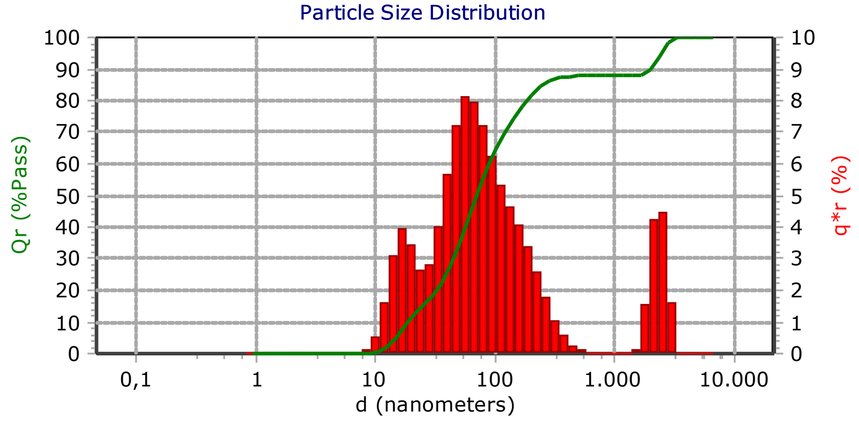


#### - Measurement Info -

| Title | |
|---|---|
| f5_GBQ_serum_onhe_label_1 | |
| Identifiers | |
| - | |
| - | |
| Database Record | 93 |
| Run Number | 2 of 3 |
| Date | 28.10.2024 |
| Time | 11:56 |
| Acquired Date | 28.10.2024 |
| Acquired Time | 11:56 |
| Serial Number | W3422 |
| Calculated Data | |
| Above Residual | 0 |
| Below Residual | 0 |
| Loading Index | 0,071 |
| Conc. Index | 0,0564 |
| RMS Residual | 0,161% |
| Cell Temp (C) | 22,10 |
| Viscosity(cp) | 0,9530 |
| Reflected Pwr (uW) | 2,80 |
| User Defined Calculations | |
| Name | Value |
| Recalculation Status | |
| DB-Meas : : Original : | |

#### -SOP Info-

| EV_ANNA(*) | |
|---|---|
| Timing | |
| Setzero Time | 30 (sec) |
| Run Time | 30 (sec) |
| Number of Runs | 3 |
| Multi-Run Delay | 0 (min) |
| Delay First Meas. | Disabled |
| Analysis | |
| Russ | |
| Refractive Index | N/A |
| Transparency | Absorb |
| Shape | Irregular |
| WATER | |
| Refractive Index | 1,33 |
| Low Temperature | 20,0 |
| Low Temp. Visc. | 1,002 |
| High Temperature | 30,0 |
| High Temp. Visc. | 0,797 |
| Options: | |
| Analysis Type | Mode |
| Perspective | |
| Progression | Standard |
| Distribution | Intensity |
| Upper Edge(nm) | 6540 |
| Lower Edge(nm) | 0,8 |
| Residuals | Disabled |

| Summary | |
|---|---|
| Data | Value |
| MI(nm): | 353,0 |
| MN(nm): | 17,30 |
| MA(nm): | 49,10 |
| CS: | 122,3 |
| SD: | 104,8 |
| PDI: | 180,9 |
| Mz: | 111,4 |
| σι: | 413,8 |
| Ski: | 767,2 |
| Kg: | 8996 |

| Percentiles | |
|---|---|
| d(i) | d(nm) |
| 10,00 | 18,73 |
| 20,00 | 33,30 |
| 30,00 | 45,60 |
| 40,00 | 57,00 |
| 50,00 | 70,80 |
| 60,00 | 90,20 |
| 70,00 | 122,9 |
| 80,00 | 185,7 |
| 90,00 | 1972 |
| 95,00 | 2400 |

| Mode Summary | | | | |
|---|---|---|---|---|
| d(nm) | Pct | Width | C(I) | C(V):cc/ml |
| 16,10 | 13,65 | 9,64E+00 | 9,64E-03 | 5,95E-07 |
| 55,98 | 51,80 | 3,66E+01 | 3,66E-02 | 7,92E-08 |
| 152,3 | 22,59 | 1,6E+01 | 1,6E-02 | 2,19E-08 |
| 2.280 | 11,95 | 8,44E+00 | 8,44E-03 | 3,21E-08 |

16.01.2026 10:50

FLEX
11.1.0.1

#### - Notes -

-

#### -Tabular Data -

| d(nm) | qr(%) | Qr(%P) | d(nm) | qr(%) | Qr(%P) |
|---|---|---|---|---|---|
| 6540 | 0,00 | 100,00 | 15,19 | 0,24 | 2,53 |
| 5500 | 0,00 | 100,00 | 12,77 | 2,29 | 2,29 |
| 4620 | 0,00 | 100,00 | 10,74 | 0,00 | 0,00 |
| 3890 | 0,00 | 100,00 | 9,03 | 0,00 | 0,00 |
| 3270 | 0,00 | 100,00 | 7,60 | 0,00 | 0,00 |
| 2750 | 0,00 | 100,00 | 6,39 | 0,00 | 0,00 |
| 2312 | 0,00 | 100,00 | 5,37 | 0,00 | 0,00 |
| 1944 | 0,00 | 100,00 | 4,52 | 0,00 | 0,00 |
| 1635 | 0,00 | 100,00 | 3,80 | 0,00 | 0,00 |
| 1375 | 0,00 | 100,00 | 3,19 | 0,00 | 0,00 |
| 1156 | 0,00 | 100,00 | 2,690 | 0,00 | 0,00 |
| 972,0 | 0,00 | 100,00 | 2,260 | 0,00 | 0,00 |
| 818,0 | 0,00 | 100,00 | 1,900 | 0,00 | 0,00 |
| 687,0 | 0,00 | 100,00 | 1,600 | 0,00 | 0,00 |
| 578,0 | 0,00 | 100,00 | 1,340 | 0,00 | 0,00 |
| 486,0 | 0,00 | 100,00 | 1,130 | 0,00 | 0,00 |
| 409,0 | 0,00 | 100,00 | 0,950 | 0,00 | 0,00 |
| 344,0 | 0,00 | 100,00 | | | |
| 289,0 | 0,00 | 100,00 | | | |
| 243,0 | 0,00 | 100,00 | | | |
| 204,4 | 0,00 | 100,00 | | | |
| 171,9 | 0,00 | 100,00 | | | |
| 144,5 | 0,00 | 100,00 | | | |
| 121,5 | 0,48 | 100,00 | | | |
| 102,2 | 65,75 | 99,52 | | | |
| 85,90 | 31,24 | 33,77 | | | |
| 72,30 | 0,00 | 2,53 | | | |
| 60,80 | 0,00 | 2,53 | | | |
| 51,10 | 0,00 | 2,53 | | | |
| 43,00 | 0,00 | 2,53 | | | |
| 36,10 | 0,00 | 2,53 | | | |
| 30,40 | 0,00 | 2,53 | | | |
| 25,55 | 0,00 | 2,53 | | | |
| 21,48 | 0,00 | 2,53 | | | |
| 18,06 | 0,00 | 2,53 | | | |

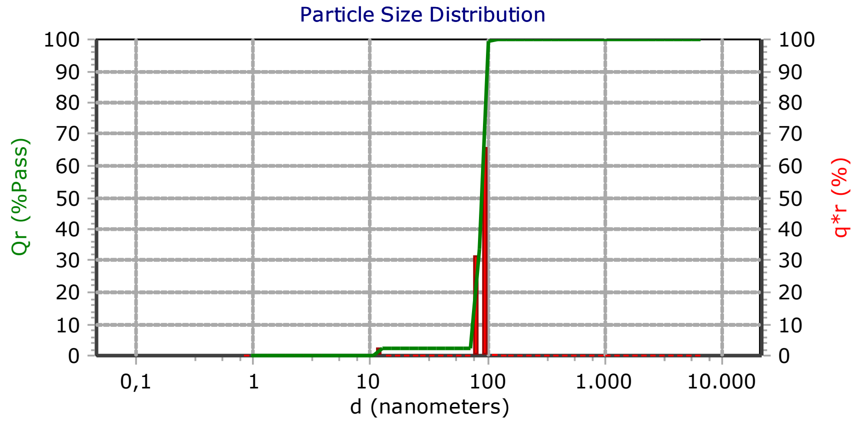


#### - Measurement Info -

| Title | |
|---|---|
| F8_GBQ_EDTA_onhe_label | |
| Identifiers | |
| - | |
| - | |
| Database Record | 63 |
| Run Number | PSD Avg Of 3 |
| Date | 11.10.2024 |
| Time | 12:15 |
| Acquired Date | 11.10.2024 |
| Acquired Time | 12:15 |
| Serial Number | W3422 |
| Calculated Data | |
| Above Residual | 0 |
| Below Residual | 0 |
| Loading Index | 0,165 |
| Conc. Index | 0,1318 |
| RMS Residual | 1,407% |
| Cell Temp (C) | 22,51 |
| Viscosity(cp) | 0,9430 |
| Reflected Pwr (uW) | 2,80 |
| User Defined Calculations | |
| Name | Value |
| Recalculation Status | |
| DB-Meas : : Original : | |

#### -SOP Info-

| EV_ANNA(*) | |
|---|---|
| Timing | |
| Setzero Time | 30 (sec) |
| Run Time | 30 (sec) |
| Number of Runs | 3 |
| Multi-Run Delay | 0 (min) |
| Delay First Meas. | Disabled |
| Analysis | |
| Russ | |
| Refractive Index | N/A |
| Transparency | Absorb |
| Shape | Irregular |
| WATER | |
| Refractive Index | 1,33 |
| Low Temperature | 20,0 |
| Low Temp. Visc. | 1,002 |
| High Temperature | 30,0 |
| High Temp. Visc. | 0,797 |
| Options: | |
| Analysis Type | Mode |
| Perspective | |
| Progression | Standard |
| Distribution | Intensity |
| Upper Edge(nm) | 6540 |
| Lower Edge(nm) | 0,8 |
| Residuals | Disabled |

| Summary | |
|---|---|
| Data | Value |
| MI(nm): | 87,30 |
| MN(nm): | 18,21 |
| MA(nm): | 76,10 |
| CS: | 78,82 |
| SD: | 8,55 |
| PDI: | 0,0528 |
| Mz: | 88,89 |
| σι: | 8,25 |
| Ski: | -134,66514 |
| Kg: | 919,1 |

| Percentiles | |
|---|---|
| d(i) | d(nm) |
| 10,00 | 77,30 |
| 20,00 | 81,60 |
| 30,00 | 84,80 |
| 40,00 | 87,30 |
| 50,00 | 89,50 |
| 60,00 | 91,70 |
| 70,00 | 93,80 |
| 80,00 | 96,20 |
| 90,00 | 98,80 |
| 95,00 | 100,6 |

Mode Summary

| d(nm) | Pct | Width | C(I) | C(V):cc/ml |
|---|---|---|---|---|
| 11,51 | 2,636 | 4,36E+00 | 4,36E-03 | 6,32E-07 |
| 87,76 | 97,36 | 1,61E+02 | 1,61E-01 | 2,21E-07 |

16.01.2026 14:51

**FLEX**

11.1.0.1

#### - Notes -

-

### - Particle Size Analysis -

C:\Microtrac\FLEX 11.1.0.1\Databases\Gonzalo\28.8.23.MDB

#### -Tabular Data -

| d(nm) | qr(%) | Qr(%P) | d(nm) | qr(%) | Qr(%P) |
|---|---|---|---|---|---|
| 6540 | 0,00 | 100,00 | 15,19 | 0,00 | 16,26 |
| 5500 | 0,00 | 100,00 | 12,77 | 0,00 | 16,26 |
| 4620 | 0,00 | 100,00 | 10,74 | 1,10 | 16,26 |
| 3890 | 0,00 | 100,00 | 9,03 | 14,35 | 15,16 |
| 3270 | 0,00 | 100,00 | 7,60 | 0,81 | 0,81 |
| 2750 | 0,00 | 100,00 | 6,39 | 0,00 | 0,00 |
| 2312 | 0,00 | 100,00 | 5,37 | 0,00 | 0,00 |
| 1944 | 0,00 | 100,00 | 4,52 | 0,00 | 0,00 |
| 1635 | 0,00 | 100,00 | 3,80 | 0,00 | 0,00 |
| 1375 | 0,00 | 100,00 | 3,19 | 0,00 | 0,00 |
| 1156 | 0,00 | 100,00 | 2,690 | 0,00 | 0,00 |
| 972,0 | 0,00 | 100,00 | 2,260 | 0,00 | 0,00 |
| 818,0 | 0,00 | 100,00 | 1,900 | 0,00 | 0,00 |
| 687,0 | 0,00 | 100,00 | 1,600 | 0,00 | 0,00 |
| 578,0 | 0,00 | 100,00 | 1,340 | 0,00 | 0,00 |
| 486,0 | 0,00 | 100,00 | 1,130 | 0,00 | 0,00 |
| 409,0 | 0,00 | 100,00 | 0,950 | 0,00 | 0,00 |
| 344,0 | 0,00 | 100,00 | | | |
| 289,0 | 0,00 | 100,00 | | | |
| 243,0 | 0,15 | 100,00 | | | |
| 204,4 | 1,58 | 99,85 | | | |
| 171,9 | 8,26 | 98,27 | | | |
| 144,5 | 21,39 | 90,01 | | | |
| 121,5 | 27,76 | 68,62 | | | |
| 102,2 | 17,93 | 40,86 | | | |
| 85,90 | 5,76 | 22,93 | | | |
| 72,30 | 0,91 | 17,17 | | | |
| 60,80 | 0,00 | 16,26 | | | |
| 51,10 | 0,00 | 16,26 | | | |
| 43,00 | 0,00 | 16,26 | | | |
| 36,10 | 0,00 | 16,26 | | | |
| 30,40 | 0,00 | 16,26 | | | |
| 25,55 | 0,00 | 16,26 | | | |
| 21,48 | 0,00 | 16,26 | | | |
| 18,06 | 0,00 | 16,26 | | | |

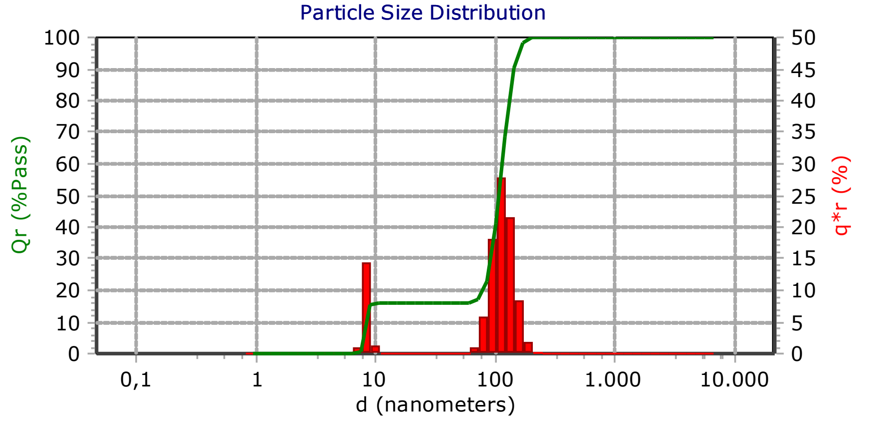


#### - Measurement Info -

| Title | |
|---|---|
| f9_GBQ_serum_onhe_label_1 | |
| Identifiers | |
| - | |
| - | |
| Database Record | 109 |
| Run Number | 2 of 3 |
| Date | 28.10.2024 |
| Time | 13:33 |
| Acquired Date | 28.10.2024 |
| Acquired Time | 13:33 |
| Serial Number | W3422 |
| Calculated Data | |
| Above Residual | 0 |
| Below Residual | 0 |
| Loading Index | 0,092 |
| Conc. Index | 0,0737 |
| RMS Residual | 1,153% |
| Cell Temp (C) | 21,37 |
| Viscosity(cp) | 0,9690 |
| Reflected Pwr (uW) | 2,80 |
| User Defined Calculations | |
| Name | Value |
| Recalculation Status | |
| DB-Meas : : Original : | |

#### -SOP Info-

| EV_ANNA(*) | |
|---|---|
| Timing | |
| Setzero Time | 30 (sec) |
| Run Time | 30 (sec) |
| Number of Runs | 3 |
| Multi-Run Delay | 0 (min) |
| Delay First Meas. | Disabled |
| Analysis | |
| Russ | |
| Refractive Index | N/A |
| Transparency | Absorb |
| Shape | Irregular |
| WATER | |
| Refractive Index | 1,33 |
| Low Temperature | 20,0 |
| Low Temp. Visc. | 1,002 |
| High Temperature | 30,0 |
| High Temp. Visc. | 0,797 |
| Options: | |
| Analysis Type | Mode |
| Perspective | |
| Progression | Standard |
| Distribution | Intensity |
| Upper Edge(nm) | 6540 |
| Lower Edge(nm) | 0,8 |
| Residuals | Disabled |

| Summary | |
|---|---|
| Data | Value |
| MI(nm): | 98,80 |
| MN(nm): | 8,46 |
| MA(nm): | 36,90 |
| CS: | 162,7 |
| SD: | 63,40 |
| PDI: | 38,20 |
| Mz: | 84,69 |
| σι: | 54,19 |
| Ski: | -457,38457 |
| Kg: | 1576 |

| Percentiles | |
|---|---|
| d(i) | d(nm) |
| 10,00 | 8,46 |
| 20,00 | 81,00 |
| 30,00 | 93,40 |
| 40,00 | 101,6 |
| 50,00 | 108,4 |
| 60,00 | 115,2 |
| 70,00 | 122,7 |
| 80,00 | 131,7 |
| 90,00 | 144,5 |
| 95,00 | 156,4 |

Mode Summary

| d(nm) | Pct | Width | C(I) | C(V):cc/ml |
|---|---|---|---|---|
| 8,10 | 16,27 | 1,5E+01 | 1,5E-02 | 6,87E-06 |
| 112,1 | 83,73 | 7,72E+01 | 7,72E-02 | 1,1E-07 |

16.01.2026 10:53

FLEX

11.1.0.1

#### - Notes -

-

Electrophoresis & Brownian Motion
Video Analysis
Laser Scattering Microscopy

ZETAVIEW®
www.particle-metrix.com

Video Operator: ZetaUser

Operator (Report): ZetaUser

**Sample Parameters**
Sample Name:
Sample Info 1:
Sample Info 2:
Sample Info 3:
Electrolyte: $H_2O$
Temperature: 25.32 °C sensed
pH 7.0 entered
Conductivity: 1130.00 µS/cm sensed

**Instrument Parameters**
Laser Wavelength: 488 nm
Filter Wavelength: Scatter
Sensitivity: 75
Shutter: 100

**SOP: SOP100_ExTu**
Size Distribution 1 Cycle 11 Positions
Description:

**Result (sizes in nm)**

| | Number | Concentration | Volume |
|---|---|---|---|
| Median (X50) | 104.0 | 104.0 | 187.1 |
| StdDev | 51.0 | 51.0 | 146.3 |

Concentration: 3.7E+7 Particles / mL
Dilution Factor: 2000
Original Concentration: 7.3E+10 Particles / mL

**Quality**
Average Counted Particles per Frame: 101
Number of Traced Particles: 1396
1 Positions Removed for Analysis

**Analysis Parameters**
Max Area 1000, Min Area 10, Min Brightness 20, nm/Class: 5

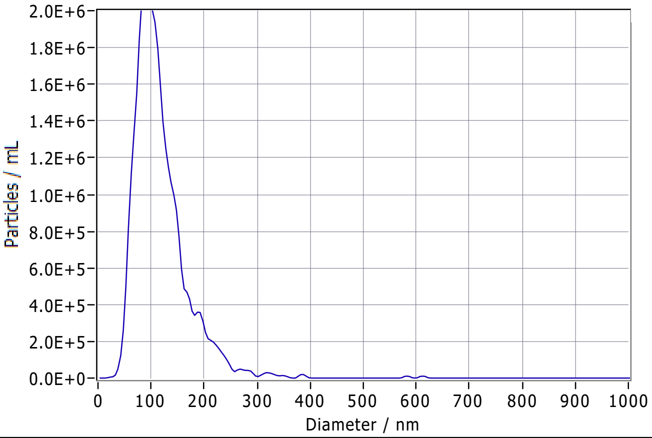


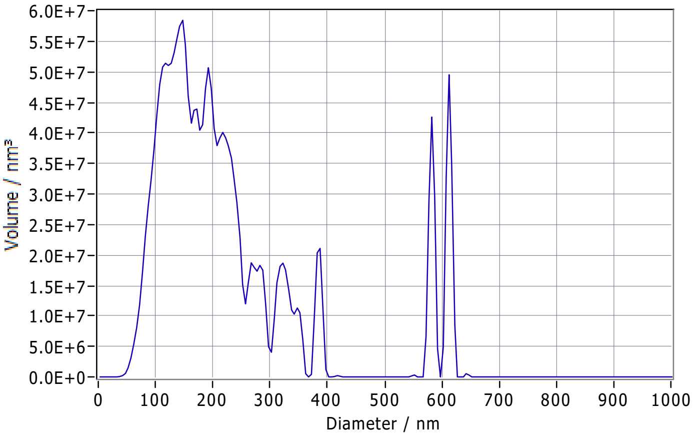


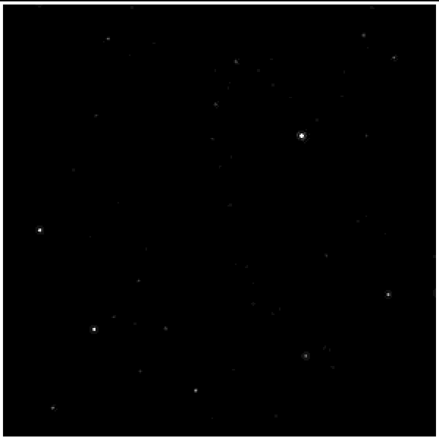

**Peak Analysis (Concentration)**

| Diameter / nm | Particles/mL | FWHM / nm | Percentage |
|---|---|---|---|
| 91.7 | 2.1E+6 | 77.3 | 100.0 |

**X Values** (all sizes are given in nm)

| | Number | Concentration | Volume |
|---|---|---|---|
| X10 | 65.2 | 65.2 | 102.3 |
| X50 | 104.0 | 104.0 | 187.1 |
| X90 | 178.6 | 178.6 | 578.3 |
| Span | 1.1 | 1.1 | 2.5 |
| Mean | 117.2 | 117.2 | 233.3 |
| StdDev | 51.0 | 51.0 | 146.3 |

**Comment**

(Signature)

Analyzed Video: C:\Users\ZetaView\Desktop\g\EDTA GBQ ONHE LABEL\20241011_0006_F1_gbq_edta_onhe_laBEL_1_size_488.avi

Experiment: 2024-10-11 10:57 ZetaView S/N 20-559, Software ZetaView (version 8.05.16 SP3)
Report: 2024-10-11 10:58 Software ZetaView (version 8.05.16 SP3)

Electrophoresis & Brownian Motion
Video Analysis
Laser Scattering Microscopy

Video Operator: ZetaUser

Operator (Report): ZetaUser

**Sample Parameters**
Sample Name:
Sample Info 1:
Sample Info 2:
Sample Info 3:
Electrolyte: $H_2O$
Temperature: 25.04 °C sensed
pH 7.0 entered
Conductivity: 1171.00 µS/cm sensed

**Instrument Parameters**
Laser Wavelength: 488 nm
Filter Wavelength: Scatter
Sensitivity: 75
Shutter: 100

**SOP: SOP100_ExTu**
Size Distribution 1 Cycle 11 Positions
Description:

**Result (sizes in nm)**

| | Number | Concentration | Volume |
|---|---|---|---|
| Median (X50) | 100.2 | 100.2 | 197.9 |
| StdDev | 52.7 | 52.7 | 145.5 |

Concentration: 3.8E+7 Particles / mL
Dilution Factor: 2000
Original Concentration: 7.6E+10 Particles / mL

**Quality**
Average Counted Particles per Frame: 104
Number of Traced Particles: 1324
1 Positions Removed for Analysis

**Analysis Parameters**
Max Area 1000, Min Area 10, Min Brightness 20, nm/Class: 5

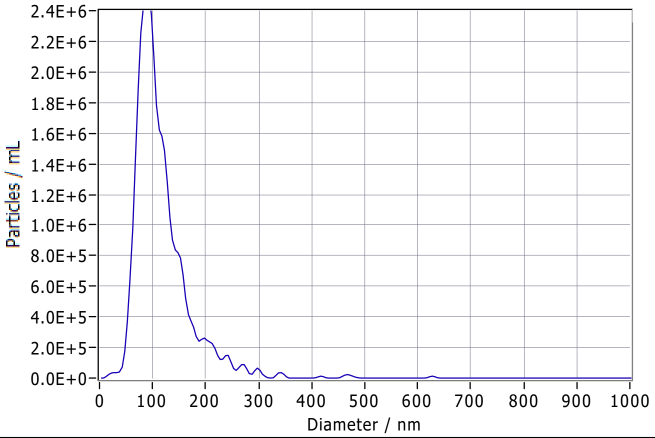


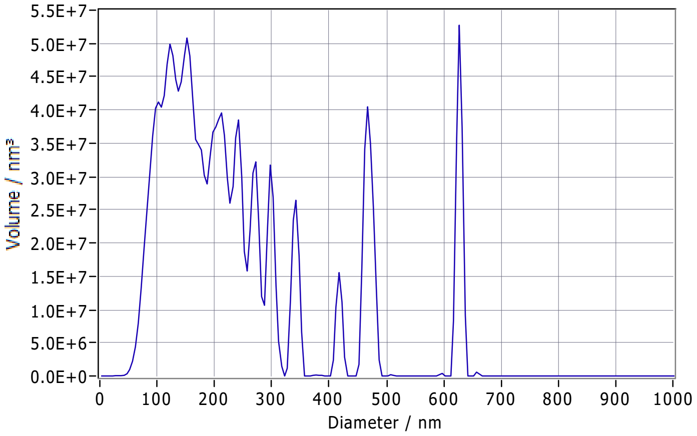


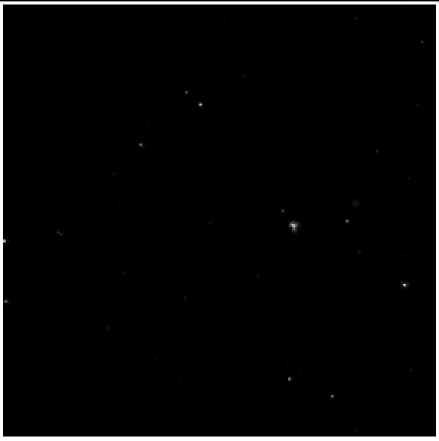

**Peak Analysis (Concentration)**

| Diameter / nm | Particles/mL | FWHM / nm | Percentage |
|---|---|---|---|
| 88.8 | 2.5E+6 | 62.4 | 100.0 |

**X Values** (all sizes are given in nm)

| | Number | Concentration | Volume |
|---|---|---|---|
| X10 | 67.3 | 67.3 | 100.0 |
| X50 | 100.2 | 100.2 | 197.9 |
| X90 | 175.0 | 175.0 | 466.0 |
| Span | 1.1 | 1.1 | 1.8 |
| Mean | 116.3 | 116.3 | 241.9 |
| StdDev | 52.7 | 52.7 | 145.5 |

**Comment**

(Signature)

Analyzed Video: C:\Users\ZetaView\Desktop\g\EDTA GBQ ONHE LABEL\20241011_0007_F1_gbq_edta_onhe_laBEL_2_size_488.avi

Experiment: 2024-10-11 10:59 ZetaView S/N 20-559, Software ZetaView (version 8.05.16 SP3)
Report: 2024-10-11 11:00 Software ZetaView (version 8.05.16 SP3)

PARTICLE METRIX

### ZETAVIEW®

www.particle-metrix.com

Electrophoresis & Brownian Motion
Video Analysis
Laser Scattering Microscopy

Video Operator: ZetaUser

Operator (Report): ZetaUser

**Sample Parameters**
Sample Name:
Sample Info 1:
Sample Info 2:
Sample Info 3:
Electrolyte: $H_2O$
Temperature: 25.15 °C sensed
pH 7.0 entered
Conductivity: 1180.00 µS/cm sensed

**Instrument Parameters**
Laser Wavelength: 488 nm
Filter Wavelength: Scatter
Sensitivity: 75
Shutter: 100

**SOP: SOP100_ExTu**
Size Distribution 1 Cycle 11 Positions
Description:

**Result (sizes in nm)**

| | Number | Concentration | Volume |
|---|---|---|---|
| Median (X50) | 130.1 | 130.1 | 182.7 |
| StdDev | 49.4 | 49.4 | 74.1 |

Concentration: 8.3E+7 Particles / mL
Dilution Factor: 500
Original Concentration: 4.1E+10 Particles / mL

**Quality**
Average Counted Particles per Frame: 227
Number of Traced Particles: 2847

**Analysis Parameters**
Max Area 1000, Min Area 10, Min Brightness 20, nm/Class: 5

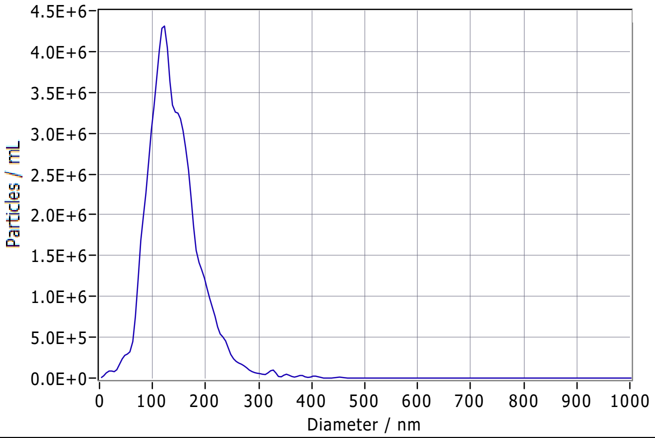


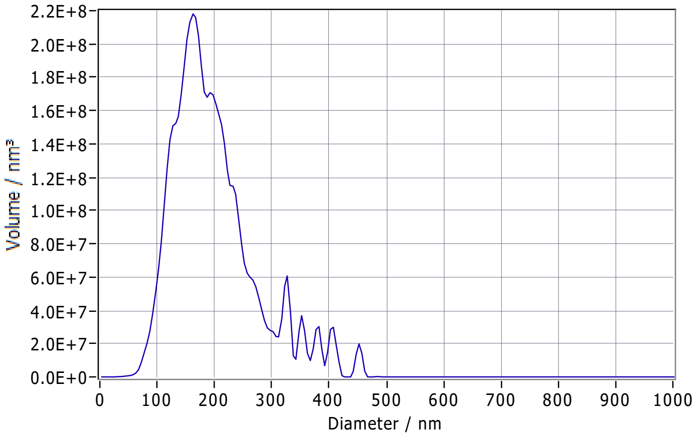


**Peak Analysis (Concentration)**

| Diameter / nm | Particles/mL | FWHM / nm | Percentage |
|---|---|---|---|
| 120.0 | 4.3E+6 | 87.8 | 100.0 |

**X Values** (all sizes are given in nm)

| | Number | Concentration | Volume |
|---|---|---|---|
| X10 | 83.3 | 83.3 | 119.0 |
| X50 | 130.1 | 130.1 | 182.7 |
| X90 | 199.5 | 199.5 | 311.7 |
| Span | 0.9 | 0.9 | 1.1 |
| Mean | 139.9 | 139.9 | 200.3 |
| StdDev | 49.4 | 49.4 | 74.1 |

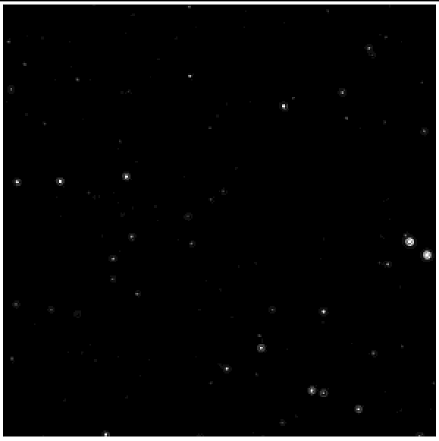

**Comment**

(Signature)

Analyzed Video: C:\Users\ZetaView\Desktop\g\EDTA GBQ ONHE LABEL\20241011_0022_F5_gbq_edta_onhe_laBEL_1_size_488.avi

Experiment: 2024-10-11 11:40 ZetaView S/N 20-559, Software ZetaView (version 8.05.16 SP3)

Report: 2024-10-11 11:43 Software ZetaView (version 8.05.16 SP3)

PARTICLE METRIX

### ZETAVIEW®

www.particle-metrix.com

Electrophoresis & Brownian Motion
Video Analysis
Laser Scattering Microscopy

Video Operator: ZetaUser

Operator (Report): ZetaUser

**Sample Parameters**
Sample Name:
Sample Info 1:
Sample Info 2:
Sample Info 3:
Electrolyte: $H_2O$
Temperature: 24.98 °C sensed
pH 7.0 entered
Conductivity: 1135.00 µS/cm sensed

**Instrument Parameters**
Laser Wavelength: 488 nm
Filter Wavelength: Scatter
Sensitivity: 75
Shutter: 100

**SOP: SOP100_ExTu**
Size Distribution 1 Cycle 11 Positions
Description:

**Result (sizes in nm)**

| | Number | Concentration | Volume |
|---|---|---|---|
| Median (X50) | 132.7 | 132.7 | 182.7 |
| StdDev | 50.5 | 50.5 | 110.4 |

Concentration: 9.4E+7 Particles / mL
Dilution Factor: 500
Original Concentration: 4.7E+10 Particles / mL

**Quality**
Average Counted Particles per Frame: 257
Number of Traced Particles: 2699

**Analysis Parameters**
Max Area 1000, Min Area 10, Min Brightness 20, nm/Class: 5

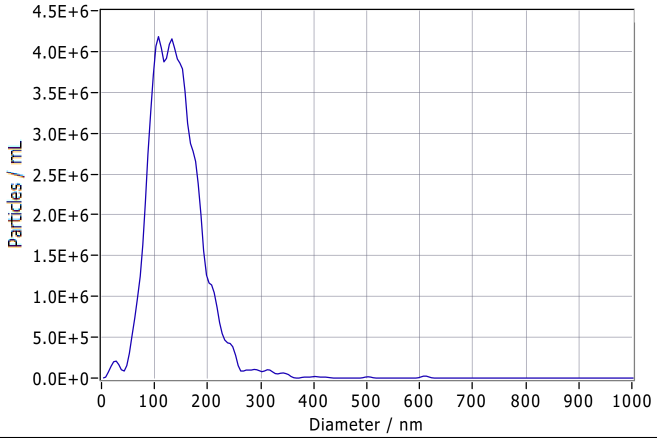


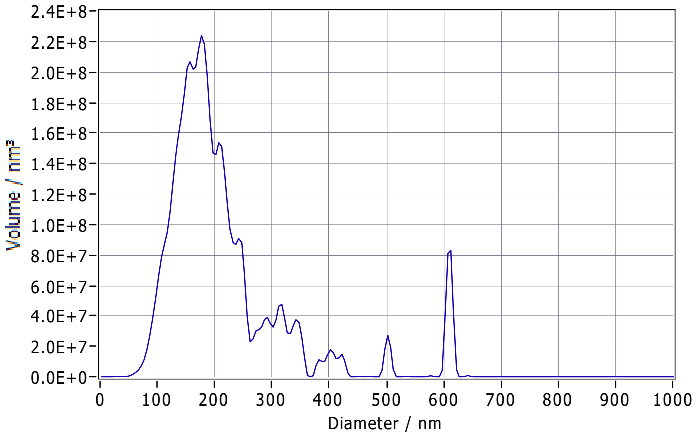


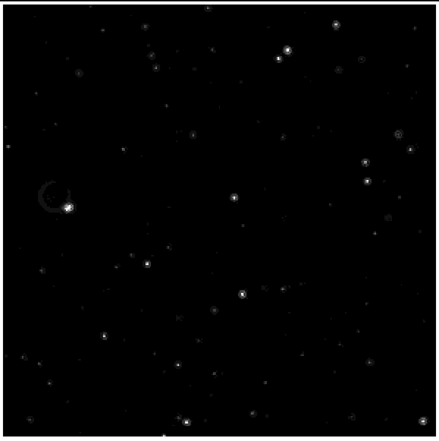

**Peak Analysis (Concentration)**

| Diameter / nm | Particles/mL | FWHM / nm | Percentage |
|---|---|---|---|
| 133.1 | 4.1E+6 | 104.2 | 100.0 |

**X Values** (all sizes are given in nm)

| | Number | Concentration | Volume |
|---|---|---|---|
| X10 | 84.0 | 84.0 | 121.4 |
| X50 | 132.7 | 132.7 | 182.7 |
| X90 | 197.9 | 197.9 | 334.9 |
| Span | 0.9 | 0.9 | 1.2 |
| Mean | 140.5 | 140.5 | 216.1 |
| StdDev | 50.5 | 50.5 | 110.4 |

**Comment**

(Signature)

Analyzed Video: C:\Users\ZetaView\Desktop\g\EDTA GBQ ONHE LABEL\20241011_0023_F5_gbq_edta_onhe_laBEL_2_size_488.avi

Experiment: 2024-10-11 11:43 ZetaView S/N 20-559, Software ZetaView (version 8.05.16 SP3)

Report: 2024-10-11 11:47 Software ZetaView (version 8.05.16 SP3)

PARTICLE METRIX ZETAVIEW® www.particle-metrix.com

Electrophoresis & Brownian Motion
Video Analysis
Laser Scattering Microscopy

Video Operator: ZetaUser

Operator (Report): ZetaUser

**Sample Parameters**
Sample Name:
Sample Info 1:
Sample Info 2:
Sample Info 3:
Electrolyte: $H_2O$
Temperature: 25.43 °C sensed
pH 7.0 entered
Conductivity: 1090.00 µS/cm sensed

**Instrument Parameters**
Laser Wavelength: 488 nm
Filter Wavelength: Scatter
Sensitivity: 75
Shutter: 100

**SOP: SOP100_ExTu**
Size Distribution 1 Cycle 11 Positions
Description:

**Result (sizes in nm)**

| | Number | Concentration | Volume |
|---|---|---|---|
| Median (X50) | 149.3 | 149.3 | 249.2 |
| StdDev | 72.7 | 72.7 | 123.8 |

Concentration: 4.1E+7 Particles / mL
Dilution Factor: 500
Original Concentration: 2.1E+10 Particles / mL

**Quality**
Average Counted Particles per Frame: 114
Number of Traced Particles: 1470
1 Positions Removed for Analysis

**Analysis Parameters**
Max Area 1000, Min Area 10, Min Brightness 20, nm/Class: 5

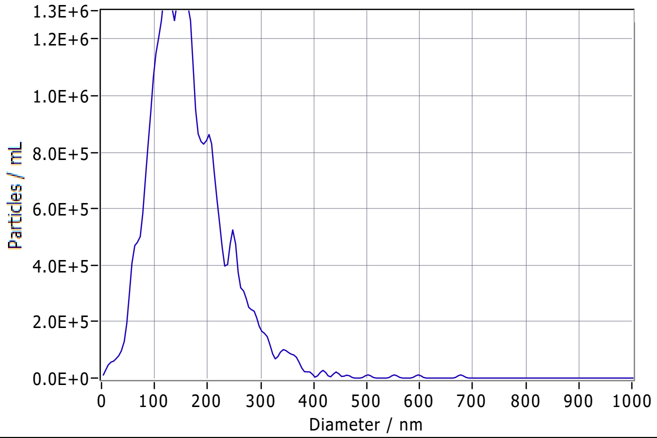


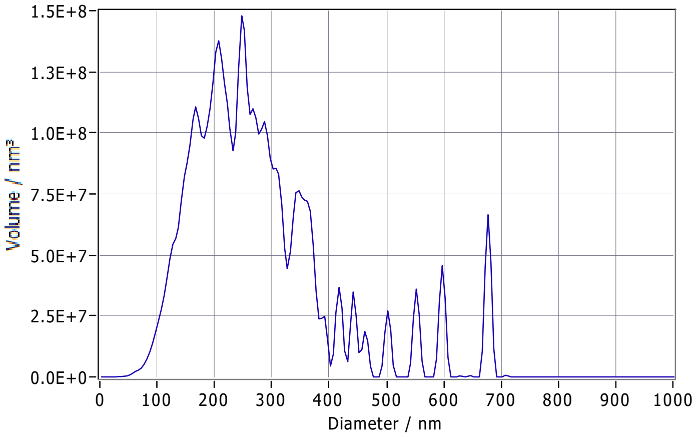


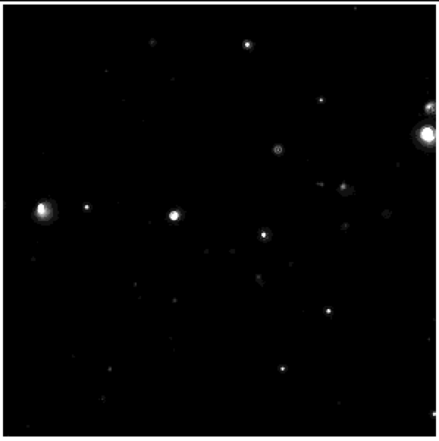

**Peak Analysis (Concentration)**

| Diameter / nm | Particles/mL | FWHM / nm | Percentage |
|---|---|---|---|
| 151.1 | 1.3E+6 | 38.7 | 43.0 |
| 124.7 | 1.4E+6 | 54.4 | 42.0 |
| 247.5 | 5.2E+5 | 38.2 | 15.0 |

**X Values** (all sizes are given in nm)

| | Number | Concentration | Volume |
|---|---|---|---|
| X10 | 81.9 | 81.9 | 147.2 |
| X50 | 149.3 | 149.3 | 249.2 |
| X90 | 254.0 | 254.0 | 422.5 |
| Span | 1.2 | 1.2 | 1.1 |
| Mean | 163.5 | 163.5 | 276.8 |
| StdDev | 72.7 | 72.7 | 123.8 |

**Comment**

(Signature)

Analyzed Video: C:\Users\ZetaView\Desktop\g\EDTA GBQ ONHE LABEL\20241011_0040_F9_gbq_edta_onhe_laBEL_1_size_488.avi

Experiment: 2024-10-11 12:31 ZetaView S/N 20-559, Software ZetaView (version 8.05.16 SP3)    Report: 2024-10-11 12:33 Software ZetaView (version 8.05.16 SP3)

**PARTICLE METRIX** **ZETAVIEW®**

www.particle-metrix.com

Electrophoresis & Brownian Motion
Video Analysis
Laser Scattering Microscopy

Video Operator: ZetaUser

Operator (Report): ZetaUser

**Sample Parameters**
Sample Name:
Sample Info 1:
Sample Info 2:
Sample Info 3:
Electrolyte: $H_2O$
Temperature: 25.00 °C sensed
pH 7.0 entered
Conductivity: 1057.00 µS/cm sensed

**Instrument Parameters**
Laser Wavelength: 488 nm
Filter Wavelength: Scatter
Sensitivity: 75
Shutter: 100

**SOP: SOP100_ExTu**
Size Distribution 1 Cycle 11 Positions
Description:

**Result (sizes in nm)**

| | Number | Concentration | Volume |
|---|---|---|---|
| Median (X50) | 151.6 | 151.6 | 233.3 |
| StdDev | 66.9 | 66.8 | 80.8 |

Concentration: 4.1E+7 Particles / mL
Dilution Factor: 500
Original Concentration: 2.1E+10 Particles / mL

**Quality**
Average Counted Particles per Frame: 113
Number of Traced Particles: 1432

**Analysis Parameters**
Max Area 1000, Min Area 10, Min Brightness 20, nm/Class: 5

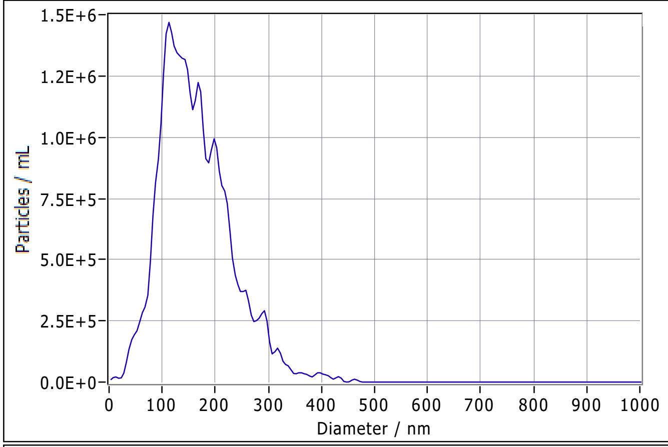


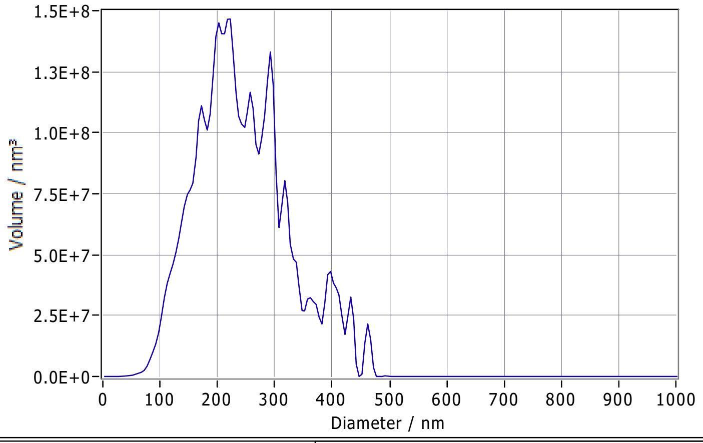


**Peak Analysis (Concentration)**

| Diameter / nm | Particles/mL | FWHM / nm | Percentage |
|---|---|---|---|
| 115.7 | 1.4E+6 | 136.8 | 93.1 |
| 287.5 | 2.8E+5 | 26.9 | 6.9 |

**X Values** (all sizes are given in nm)

| | Number | Concentration | Volume |
|---|---|---|---|
| X10 | 87.5 | 87.5 | 144.6 |
| X50 | 151.6 | 151.6 | 233.3 |
| X90 | 252.7 | 252.7 | 360.0 |
| Span | 1.1 | 1.1 | 0.9 |
| Mean | 164.1 | 164.1 | 245.9 |
| StdDev | 66.9 | 66.8 | 80.8 |

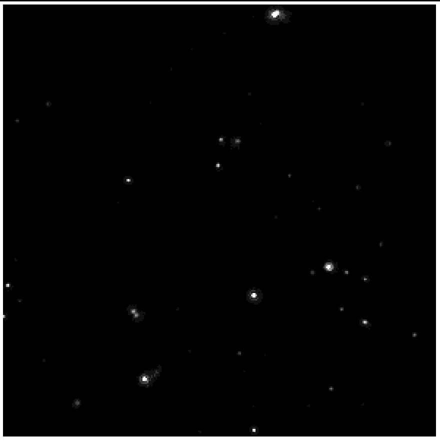

**Comment**

(Signature)

Analyzed Video: C:\Users\ZetaView\Desktop\g\EDTA GBQ ONHE LABEL\20241011_0041_F9_gbq_edta_onhe_laBEL_2_size_488.avi

Experiment: 2024-10-11 12:34 ZetaView S/N 20-559, Software ZetaView (version 8.05.16 SP3)

Report: 2024-10-11 12:35 Software ZetaView (version 8.05.16 SP3)

### e_id: 3468, ZetaScript: SMALL SOP100 zp, Sample name: F1_GBQ_EDTA_onhe_label

Measurement created: 2024-10-11 15:36:09 Report created: 2024-10-11 16:10:59 User: ZetaUser

#### Measurement specification: SOP100(1)

##### Capture settings

| | | | |
|---|---|---|---|
| Temperature / °C | 24.9 | Laser wave length / nm | 488 |
| Dilution | 2000 | Filter wave length / nm | 0 |
| Viscosity / mPa · s | 0.902 | Sensitivity | 80 |
| Positions | None | Shutter | 90 |
| Video length | 30 | | |
| Frame rate / 1/s | 30.303 | | |
| Experiment type | Size distribution | | |

##### Analysis settings

| | |
|---|---|
| Concentration correction | 0.55 |
| Concentration calibration | 636400 |
| Min size /nm | 10 |
| Max size / nm | 1000 |
| Min area / px | 1 |
| Max area / px | 1000 |
| Trace length | 15 |

Comment:

#### Graphics:

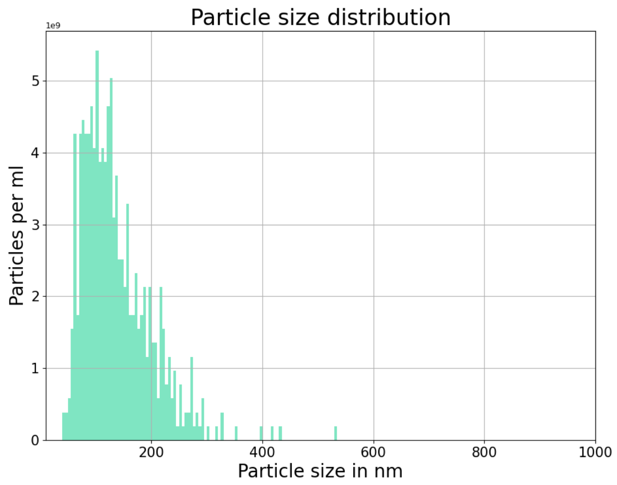


#### Results:

| | Peak /nm | Median /nm | #Particles | Avg. Num Det. | FWHM /nm | Concentration /1/mL | Position |
|---|---|---|---|---|---|---|---|
| | **121.5 (± 29.3)** | **129.7 (± 18.3)** | **569** | **157.4 (± 4.4)** | **124.9 (± 39.3)** | **1.10E+11 (± 3.1E+09)** | |
| 1 | 107.3 | 121.8 | 91 | 155.9 | 109.0 | 1.09E+11 | 0.30 |
| 2 | 134.8 | 133.2 | 50 | 165.6 | 101.9 | 1.16E+11 | 0.40 |
| 3 | 184.0 | 163.0 | 49 | 158.5 | 209.6 | 1.11E+11 | 0.50 |
| 4 | 129.5 | 148.7 | 56 | 156.4 | 155.1 | 1.09E+11 | 0.60 |
| 5 | 102.3 | 116.2 | 89 | 160.2 | 100.4 | 1.12E+11 | 0.70 |
| 6 | 92.5 | 109.8 | 111 | 150.6 | 100.6 | 1.05E+11 | 0.80 |
| 7 | 100.3 | 115.0 | 123 | 154.3 | 97.6 | 1.08E+11 | 0.85 |

### e_id: 3468, ZetaScript: SMALL SOP100 zp, Sample name: F1_GBQ_EDTA_onhe_label

Measurement created: 2024-10-11 15:36:09 Report created: 2024-10-11 16:11:00 User: ZetaUser

#### Measurement specification: SOP100(2)

##### Capture settings

| | | | |
|---|---|---|---|
| Temperature / °C | 24.9 | Laser wave length / nm | 488 |
| Dilution | 2000 | Filter wave length / nm | 0 |
| Viscosity / mPa · s | 0.901 | Sensitivity | 80 |
| Positions | None | Shutter | 90 |
| Video length | 30 | | |
| Frame rate / 1/s | 30.303 | | |
| Experiment type | Size distribution | | |

##### Analysis settings

| | |
|---|---|
| Concentration correction | 0.55 |
| Concentration calibration | 636400 |
| Min size /nm | 10 |
| Max size / nm | 1000 |
| Min area / px | 1 |
| Max area / px | 1000 |
| Trace length | 15 |

Comment:

#### Graphics:

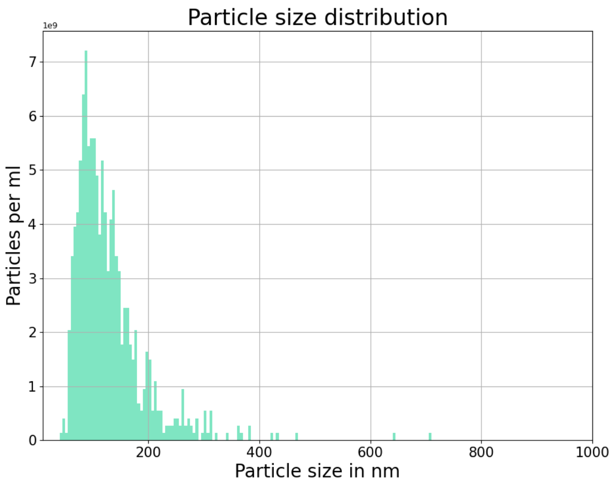


#### Results:

| | Peak /nm | Median /nm | #Particles | Avg. Num Det. | FWHM /nm | Concentration /1/mL | Position |
|---|---|---|---|---|---|---|---|
| | **96.7 (± 8.0)** | **113.1 (± 6.6)** | **835** | **162.4 (± 5.7)** | **95.1 (± 9.1)** | **1.14E+11 (± 4.0E+09)** | |
| 1 | 85.1 | 99.9 | 129 | 156.7 | 81.9 | 1.10E+11 | 0.30 |
| 2 | 87.4 | 107.3 | 117 | 172.3 | 94.2 | 1.21E+11 | 0.40 |
| 3 | 94.7 | 117.6 | 115 | 158.9 | 107.2 | 1.11E+11 | 0.50 |
| 4 | 109.9 | 119.9 | 111 | 162.7 | 101.0 | 1.14E+11 | 0.60 |
| 5 | 101.0 | 118.3 | 121 | 169.3 | 105.3 | 1.19E+11 | 0.70 |
| 6 | 102.4 | 115.0 | 125 | 156.9 | 90.2 | 1.10E+11 | 0.80 |
| 7 | 96.7 | 113.4 | 117 | 159.8 | 85.6 | 1.12E+11 | 0.85 |

### e_id: 3472, ZetaScript: SMALL SOP100 zp, Sample name: F5_GBQ_EDTA_onhe_label

Measurement created: 2024-10-11 15:52:53 Report created: 2024-10-11 16:11:09 User: ZetaUser

#### Measurement specification: SOP100(1)

##### Capture settings

| | | | |
|---|---|---|---|
| Temperature / °C | 25.0 | Laser wave length / nm | 488 |
| Dilution | 2000 | Filter wave length / nm | 0 |
| Viscosity / mPa · s | 0.9 | Sensitivity | 80 |
| Positions | None | Shutter | 90 |
| Video length | 30 | | |
| Frame rate / 1/s | 30.303 | | |
| Experiment type | Size distribution | | |

##### Analysis settings

| | |
|---|---|
| Concentration correction | 0.55 |
| Concentration calibration | 636400 |
| Min size /nm | 10 |
| Max size / nm | 1000 |
| Min area / px | 1 |
| Max area / px | 1000 |
| Trace length | 15 |

Comment:

#### Graphics:

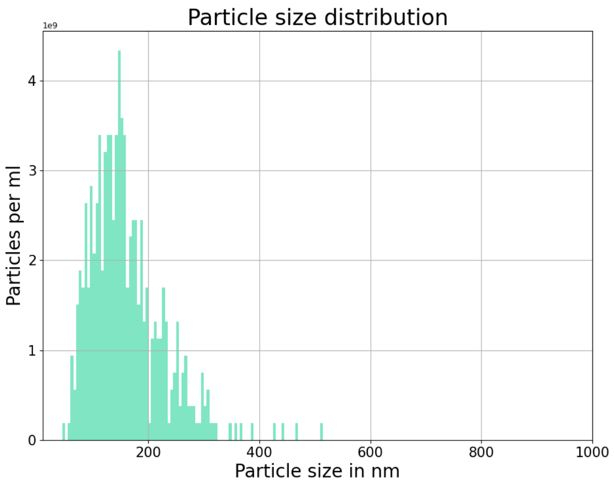


#### Results:

| | Peak /nm | Median /nm | #Particles | Avg. Num Det. | FWHM /nm | Concentration /1/mL | Position |
|---|---|---|---|---|---|---|---|
| | **140.4 (± 16.2)** | **151.3 (± 14.1)** | **452** | **121.7 (± 8.0)** | **127.4 (± 26.5)** | **8.52E+10 (± 5.6E+09)** | |
| 1 | 117.9 | 141.6 | 73 | 109.9 | 129.6 | 7.69E+10 | 0.30 |
| 2 | 169.0 | 170.5 | 55 | 125.8 | 98.7 | 8.81E+10 | 0.40 |
| 3 | 152.6 | 170.6 | 41 | 132.9 | 158.4 | 9.30E+10 | 0.50 |
| 4 | 145.6 | 157.7 | 45 | 129.8 | 170.3 | 9.09E+10 | 0.60 |
| 5 | 141.3 | 147.3 | 58 | 118.1 | 94.2 | 8.27E+10 | 0.70 |
| 6 | 132.5 | 134.6 | 90 | 122.9 | 114.5 | 8.61E+10 | 0.80 |
| 7 | 123.8 | 136.5 | 90 | 112.5 | 126.2 | 7.88E+10 | 0.85 |

### e_id: 3472, ZetaScript: SMALL SOP100 zp, Sample name: F5_GBQ_EDTA_onhe_label

Measurement created: 2024-10-11 15:52:53 Report created: 2024-10-11 16:11:10 User: ZetaUser

#### Measurement specification: SOP100(2)

##### Capture settings

| | | | |
|---|---|---|---|
| Temperature / °C | 25.0 | Laser wave length / nm | 488 |
| Dilution | 2000 | Filter wave length / nm | 0 |
| Viscosity / mPa · s | 0.901 | Sensitivity | 80 |
| Positions | None | Shutter | 90 |
| Video length | 30 | | |
| Frame rate / 1/s | 30.303 | | |
| Experiment type | Size distribution | | |

##### Analysis settings

| | |
|---|---|
| Concentration correction | 0.55 |
| Concentration calibration | 636400 |
| Min size /nm | 10 |
| Max size / nm | 1000 |
| Min area / px | 1 |
| Max area / px | 1000 |
| Trace length | 15 |

Comment:

#### Graphics:

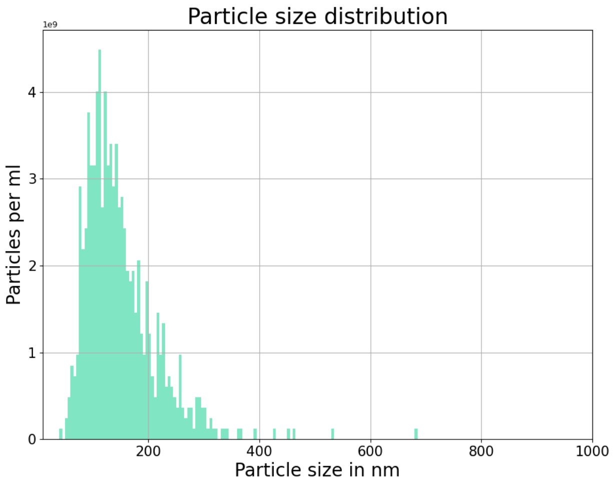


#### Results:

| | Peak /nm | Median /nm | #Particles | Avg. Num Det. | FWHM /nm | Concentration /1/mL | Position |
|---|---|---|---|---|---|---|---|
| | **120.5 (± 10.7)** | **135.0 (± 6.2)** | **702** | **121.7 (± 7.9)** | **115.2 (± 20.6)** | **8.52E+10 (± 5.5E+09)** | |
| 1 | 124.6 | 142.0 | 100 | 112.8 | 140.2 | 7.90E+10 | 0.30 |
| 2 | 114.8 | 133.5 | 116 | 133.0 | 103.9 | 9.31E+10 | 0.40 |
| 3 | 116.0 | 131.2 | 96 | 126.7 | 87.6 | 8.87E+10 | 0.50 |
| 4 | 113.4 | 129.2 | 93 | 124.4 | 102.3 | 8.71E+10 | 0.60 |
| 5 | 108.8 | 126.0 | 106 | 126.0 | 137.3 | 8.82E+10 | 0.70 |
| 6 | 143.7 | 141.1 | 89 | 108.5 | 97.7 | 7.59E+10 | 0.80 |
| 7 | 122.0 | 142.1 | 102 | 120.6 | 137.2 | 8.44E+10 | 0.85 |

### e_id: 3476, ZetaScript: SMALL SOP100 zp, Sample name: F9_GBQ_EDTA_onhe_label

Measurement created: 2024-10-11 16:04:58 Report created: 2024-10-11 16:11:19 User: ZetaUser

#### Measurement specification: SOP100(1)

##### Capture settings

| | | | |
|---|---|---|---|
| Temperature / °C | 24.9 | Laser wave length / nm | 488 |
| Dilution | 1000 | Filter wave length / nm | 0 |
| Viscosity / mPa · s | 0.901 | Sensitivity | 80 |
| Positions | None | Shutter | 90 |
| Video length | 30 | | |
| Frame rate / 1/s | 30.303 | | |
| Experiment type | Size distribution | | |

##### Analysis settings

| | |
|---|---|
| Concentration correction | 0.55 |
| Concentration calibration | 636400 |
| Min size /nm | 10 |
| Max size / nm | 1000 |
| Min area / px | 1 |
| Max area / px | 1000 |
| Trace length | 15 |

Comment:

#### Graphics:

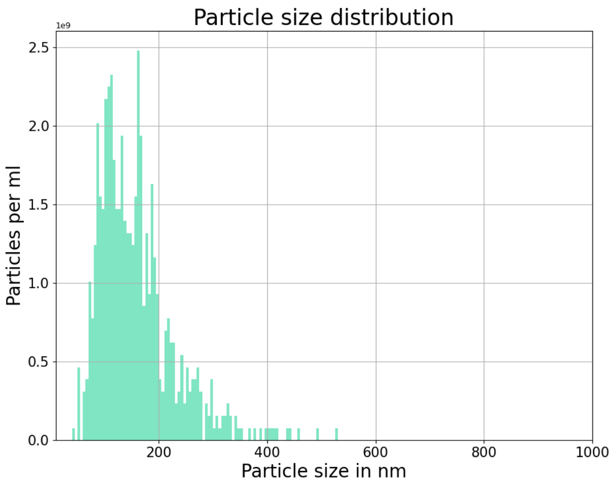


#### Results:

| | Peak /nm | Median /nm | #Particles | Avg. Num Det. | FWHM /nm | Concentration /1/mL | Position |
|---|---|---|---|---|---|---|---|
| | **125.6 (± 16.5)** | **143.4 (± 13.4)** | **657** | **145.5 (± 8.1)** | **128.6 (± 9.1)** | **5.09E+10 (± 2.8E+09)** | |
| 1 | 143.8 | 159.4 | 62 | 149.0 | 114.5 | 5.22E+10 | 0.30 |
| 2 | 150.2 | 161.5 | 78 | 142.3 | 143.6 | 4.98E+10 | 0.40 |
| 3 | 123.6 | 138.7 | 105 | 143.0 | 126.5 | 5.01E+10 | 0.50 |
| 4 | 113.7 | 142.4 | 128 | 155.5 | 139.0 | 5.44E+10 | 0.60 |
| 5 | 132.5 | 147.6 | 98 | 153.8 | 127.2 | 5.38E+10 | 0.70 |
| 6 | 115.8 | 134.6 | 101 | 145.6 | 127.3 | 5.10E+10 | 0.80 |
| 7 | 99.4 | 119.9 | 85 | 129.3 | 122.3 | 4.52E+10 | 0.85 |

### e_id: 3476, ZetaScript: SMALL SOP100 zp, Sample name: F9_GBQ_EDTA_onhe_label

Measurement created: 2024-10-11 16:04:58 Report created: 2024-10-11 16:11:20 User: ZetaUser

#### Measurement specification: SOP100(2)

##### Capture settings

| | | | |
|---|---|---|---|
| Temperature / °C | 24.9 | Laser wave length / nm | 488 |
| Dilution | 1000 | Filter wave length / nm | 0 |
| Viscosity / mPa · s | 0.901 | Sensitivity | 80 |
| Positions | None | Shutter | 90 |
| Video length | 30 | | |
| Frame rate / 1/s | 30.303 | | |
| Experiment type | Size distribution | | |

##### Analysis settings

| | |
|---|---|
| Concentration correction | 0.55 |
| Concentration calibration | 636400 |
| Min size /nm | 10 |
| Max size / nm | 1000 |
| Min area / px | 1 |
| Max area / px | 1000 |
| Trace length | 15 |

Comment:

#### Graphics:

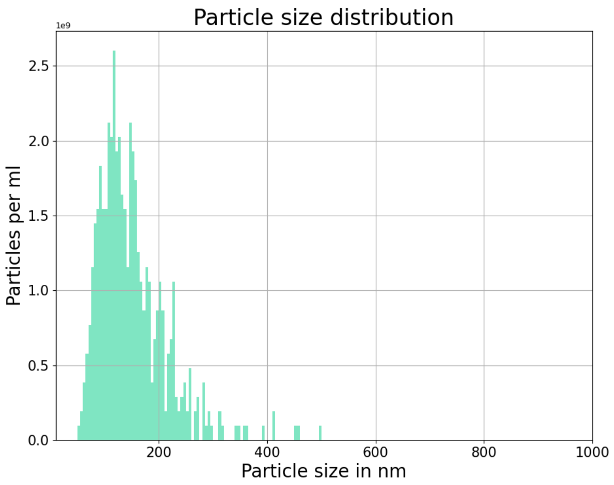


#### Results:

| | Peak /nm | Median /nm | #Particles | Avg. Num Det. | FWHM /nm | Concentration /1/mL | Position |
|---|---|---|---|---|---|---|---|
| | **126.6 (± 17.0)** | **136.8 (± 15.2)** | **498** | **137.2 (± 15.2)** | **105.8 (± 18.5)** | **4.80E+10 (± 5.3E+09)** | |
| 1 | 121.9 | 136.3 | 79 | 148.0 | 125.2 | 5.18E+10 | 0.30 |
| 2 | 113.1 | 123.5 | 55 | 139.6 | 86.0 | 4.89E+10 | 0.40 |
| 3 | 152.9 | 156.4 | 65 | 154.0 | 92.0 | 5.39E+10 | 0.50 |
| 4 | 152.7 | 161.1 | 58 | 144.4 | 129.5 | 5.05E+10 | 0.60 |
| 5 | 118.1 | 137.1 | 66 | 138.5 | 115.7 | 4.85E+10 | 0.70 |
| 6 | 109.7 | 120.7 | 98 | 132.8 | 113.2 | 4.65E+10 | 0.80 |
| 7 | 117.8 | 122.3 | 77 | 103.3 | 78.8 | 3.62E+10 | 0.85 |
