## Supplementary Material 4 for "PHoNUPS: Open-Source Software for Standardized Analysis and Visualization of Multi-Instrument Extracellular Vesicle Measurements"

### e_id: 5099, ZetaScript: SMALL SOP100 zp FM488, Sample name: CD9-GFP alone

Measurement created: 2025-04-07 13:10:18 Report created: 2025-04-07 14:33:38 User: ZetaUser

#### Measurement specification: SOP100_ZP

##### Capture settings

| | | | |
|---|---|---|---|
| Temperature / °C | 25.1 | Laser wave length / nm | 488 |
| Dilution | 1000 | Filter wave length / nm | 0 |
| Viscosity / mPa · s | 0.898 | Sensitivity | 80 |
| Positions | 0.149 | Shutter | 90 |
| Cycle frames | 30 | Dielectric constant | 78.388 |
| cycle amount | 4 | - Uref / mV | -14.349 |
| Frame rate / 1/s | 30.303 | + Uref / mV | 14.788 |
| Experiment type | Zeta Potential | Conductivity / µS/cm | 1229.0 |
| Comment: | | | |

##### Analysis settings

| | |
|---|---|
| Concentration correction | 1 |
| Concentration calibration | 636400 |
| Henry factor | 1.5 |

#### Graphics:

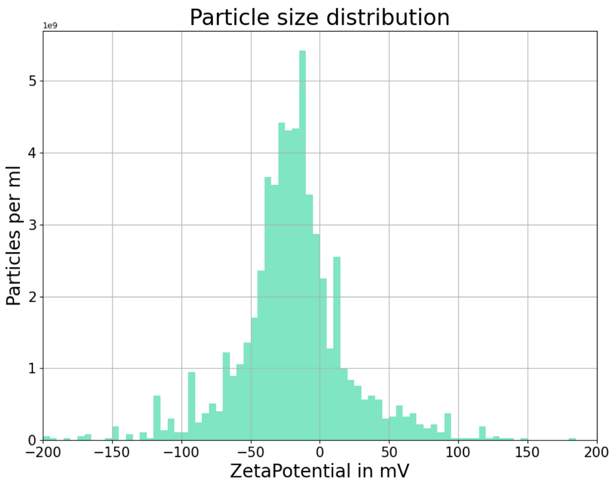


#### Results:

| | Peak /mV | Median /mV | #Zeta particles | #All particles | Avg. Num Det. | FWHM /mV | Concentration /1/mL | Position |
|---|---|---|---|---|---|---|---|---|
| | **-18.3 (± 2.0)** | **-19.5 (± 1.7)** | **2177** | **2577** | **109.8 (± 4.9)** | **50.6 (± 5.0)** | **6.99E+10 (± 3.1E+09)** | |
| 1 | -16.3 | -17.8 | 1160 | 1369 | 104.9 | 55.6 | 6.68E+10 | 0.15 |
| 2 | -20.3 | -21.2 | 1017 | 1208 | 114.7 | 45.5 | 7.30E+10 | 0.85 |

### e_id: 5099, ZetaScript: SMALL SOP100 zp FM488, Sample name: CD9-GFP alone

Measurement created: 2025-04-07 13:10:18 Report created: 2025-04-07 14:33:38 User: ZetaUser

#### Measurement specification: SOP100_ZP_2

##### Capture settings

| | | | |
|---|---|---|---|
| Temperature / °C | 25.1 | Laser wave length / nm | 488 |
| Dilution | 1000 | Filter wave length / nm | 0 |
| Viscosity / mPa · s | 0.899 | Sensitivity | 80 |
| Positions | 0.149 | Shutter | 90 |
| Cycle frames | 30 | Dielectric constant | 78.409 |
| cycle amount | 4 | - Uref / mV | -14.36 |
| Frame rate / 1/s | 30.303 | + Uref / mV | 14.583 |
| Experiment type | Zeta Potential | Conductivity / µS/cm | 1248.0 |
| Comment: | | | |

##### Analysis settings

| | |
|---|---|
| Concentration correction | 1 |
| Concentration calibration | 636400 |
| Henry factor | 1.5 |

#### Graphics:

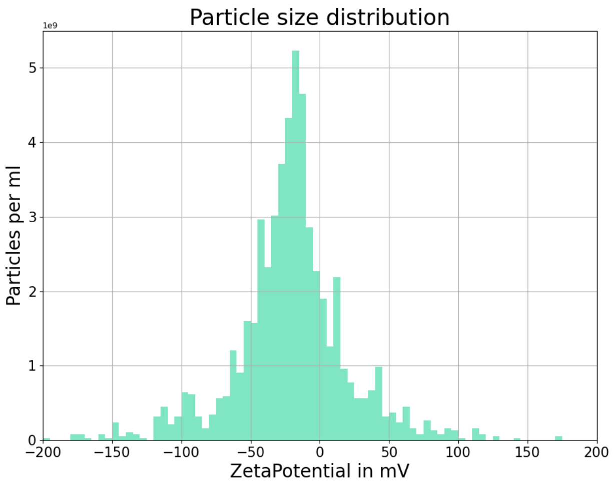


#### Results:

| | Peak /mV | Median /mV | #Zeta particles | #All particles | Avg. Num Det. | FWHM /mV | Concentration /1/mL | Position |
|---|---|---|---|---|---|---|---|---|
| | **-18.9 (± 1.6)** | **-19.9 (± 1.0)** | **2051** | **2470** | **103.6 (± 11.9)** | **48.7 (± 1.5)** | **6.60E+10 (± 7.5E+09)** | |
| 1 | -17.4 | -19.0 | 1098 | 1315 | 115.5 | 50.2 | 7.35E+10 | 0.15 |
| 2 | -20.5 | -20.9 | 953 | 1155 | 91.8 | 47.1 | 5.84E+10 | 0.85 |

### e_id: 5098, ZetaScript: SMALL SOP100 zp, Sample name: R25 alone

Measurement created: 2025-04-07 13:01:37 Report created: 2025-04-07 14:33:33 User: admin

#### Measurement specification: SOP100_ZP

##### Capture settings

| | | | |
|---|---|---|---|
| Temperature / °C | 25.0 | Laser wave length / nm | 488 |
| Dilution | 1000 | Filter wave length / nm | 0 |
| Viscosity / mPa · s | 0.899 | Sensitivity | 80 |
| Positions | 0.149 | Shutter | 90 |
| Cycle frames | 30 | Dielectric constant | 78.42 |
| cycle amount | 4 | - Uref / mV | -14.895 |
| Frame rate / 1/s | 30.303 | + Uref / mV | 15.311 |
| Experiment type | Zeta Potential | Conductivity / µS/cm | 1296.0 |
| Comment: | | | |

##### Analysis settings

| | |
|---|---|
| Concentration correction | 1 |
| Concentration calibration | 636400 |
| Henry factor | 1.5 |

#### Graphics:

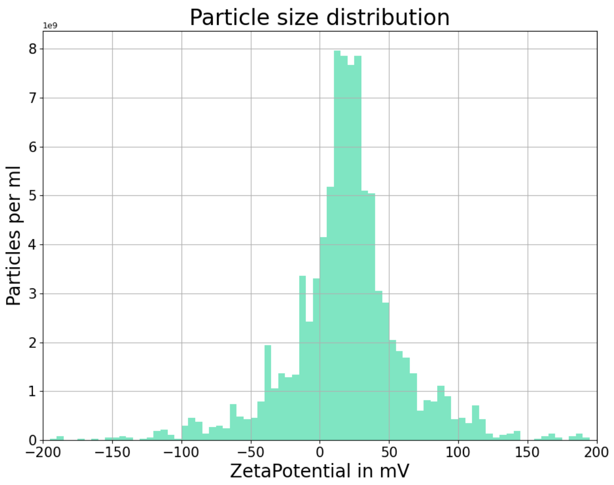


#### Results:

| | Peak /mV | Median /mV | #Zeta particles | #All particles | Avg. Num Det. | FWHM /mV | Concentration /1/mL | Position |
|---|---|---|---|---|---|---|---|---|
| | **18.8 (± 1.7)** | **19.8 (± 1.4)** | **3439** | **4067** | **174.3 (± 2.8)** | **42.7 (± 2.0)** | **1.11E+11 (± 1.8E+09)** | |
| 1 | 20.5 | 21.1 | 1762 | 2074 | 177.1 | 40.7 | 1.13E+11 | 0.15 |
| 2 | 17.1 | 18.4 | 1677 | 1993 | 171.5 | 44.7 | 1.09E+11 | 0.85 |

### e_id: 5098, ZetaScript: SMALL SOP100 zp, Sample name: R25 alone

Measurement created: 2025-04-07 13:01:37 Report created: 2025-04-07 14:33:34 User: admin

#### Measurement specification: SOP100_ZP_2

##### Capture settings

| | | | |
|---|---|---|---|
| Temperature / °C | 25.1 | Laser wave length / nm | 488 |
| Dilution | 1000 | Filter wave length / nm | 0 |
| Viscosity / mPa · s | 0.899 | Sensitivity | 80 |
| Positions | 0.149 | Shutter | 90 |
| Cycle frames | 30 | Dielectric constant | 78.409 |
| cycle amount | 4 | - Uref / mV | -14.983 |
| Frame rate / 1/s | 30.303 | + Uref / mV | 15.405 |
| Experiment type | Zeta Potential | Conductivity / µS/cm | 1190.0 |
| Comment: | | | |

##### Analysis settings

| | |
|---|---|
| Concentration correction | 1 |
| Concentration calibration | 636400 |
| Henry factor | 1.5 |

#### Graphics:

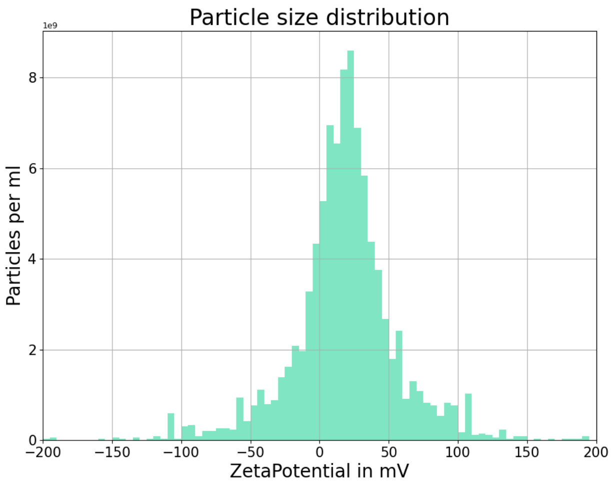


#### Results:

| | Peak /mV | Median /mV | #Zeta particles | #All particles | Avg. Num Det. | FWHM /mV | Concentration /1/mL | Position |
|---|---|---|---|---|---|---|---|---|
| | **18.5 (± 1.1)** | **18.8 (± 1.4)** | **3352** | **4022** | **180.0 (± 7.6)** | **45.5 (± 0.9)** | **1.15E+11 (± 4.8E+09)** | |
| 1 | 19.6 | 20.2 | 1817 | 2169 | 187.6 | 46.4 | 1.19E+11 | 0.15 |
| 2 | 17.4 | 17.5 | 1535 | 1853 | 172.4 | 44.5 | 1.10E+11 | 0.85 |
