## Supplementary Material 5 for "PHoNUPS: Open-Source Software for Standardized Analysis and Visualization of Multi-Instrument Extracellular Vesicle Measurements"

### e_id: 3324, ZetaScript: SMALL SOP100 zp, Sample name: P-S-RPMI-SOP100

Measurement created: 2024-09-27 11:39:49 Report created: 2025-03-26 14:56:32 User: ZetaUser

#### Measurement specification: SOP100_ZP

##### Capture settings

| | | | |
|---|---|---|---|
| Temperature / °C | 24.9 | Laser wave length / nm | 488 |
| Dilution | 5000 | Filter wave length / nm | 0 |
| Viscosity / mPa · s | 0.903 | Sensitivity | 80 |
| Positions | 0.149 | Shutter | 90 |
| Cycle frames | 30 | Dielectric constant | 78.473 |
| cycle amount | 4 | - Uref / mV | -14.46 |
| Frame rate / 1/s | 30.303 | + Uref / mV | 14.57 |
| Experiment type | Zeta Potential | Conductivity / µS/cm | 1180.0 |
| Comment: | | | |

##### Analysis settings

| | |
|---|---|
| Concentration correction | 1 |
| Concentration calibration | 636400 |
| Henry factor | 1.5 |

#### Graphics:

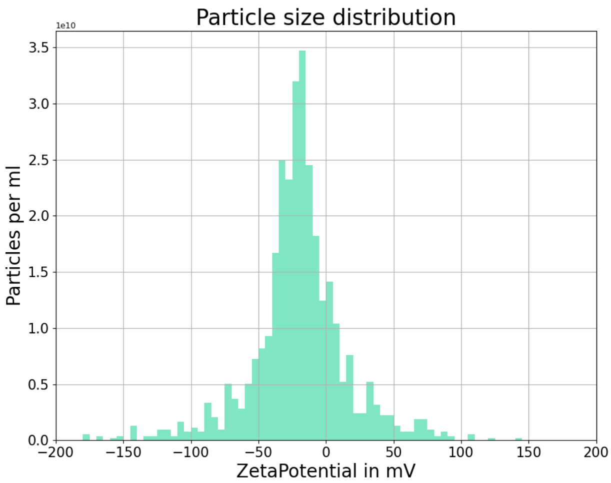


#### Results:

| | Peak /mV | Median /mV | #Zeta particles | #All particles | Avg. Num Det. | FWHM /mV | Concentration /1/mL | Position |
|---|---|---|---|---|---|---|---|---|
| | **-20.2 (± 0.4)** | **-20.1 (± 1.2)** | **1669** | **1972** | **115.2 (± 2.4)** | **38.7 (± 1.8)** | **3.67E+11 (± 7.5E+09)** | |
| 1 | -20.6 | -21.2 | 817 | 963 | 112.8 | 40.5 | 3.59E+11 | 0.15 |
| 2 | -19.8 | -18.9 | 852 | 1009 | 117.5 | 37.0 | 3.74E+11 | 0.85 |

### e_id: 3324, ZetaScript: SMALL SOP100 zp, Sample name: P-S-RPMI-SOP100

Measurement created: 2024-09-27 11:39:49 Report created: 2025-03-26 14:56:32 User: ZetaUser

#### Measurement specification: SOP100_ZP_2

##### Capture settings

| | | | |
|---|---|---|---|
| Temperature / °C | 25.1 | Laser wave length / nm | 488 |
| Dilution | 5000 | Filter wave length / nm | 0 |
| Viscosity / mPa · s | 0.897 | Sensitivity | 80 |
| Positions | 0.149 | Shutter | 90 |
| Cycle frames | 30 | Dielectric constant | 78.381 |
| cycle amount | 4 | - Uref / mV | -14.101 |
| Frame rate / 1/s | 30.303 | + Uref / mV | 14.25 |
| Experiment type | Zeta Potential | Conductivity / µS/cm | 1267.0 |
| Comment: | | | |

##### Analysis settings

| | |
|---|---|
| Concentration correction | 1 |
| Concentration calibration | 636400 |
| Henry factor | 1.5 |

#### Graphics:

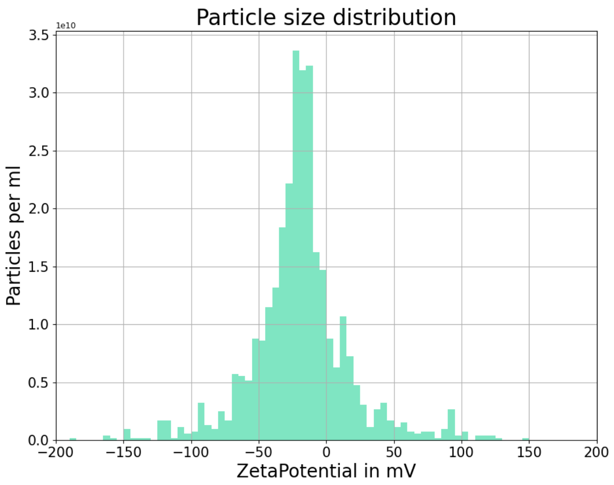


#### Results:

| | Peak /mV | Median /mV | #Zeta particles | #All particles | Avg. Num Det. | FWHM /mV | Concentration /1/mL | Position |
|---|---|---|---|---|---|---|---|---|
| | **-17.1 (± 2.1)** | **-19.5 (± 1.3)** | **1613** | **1905** | **114.5 (± 7.0)** | **34.5 (± 0.2)** | **3.64E+11 (± 2.2E+10)** | |
| 1 | -19.1 | -20.8 | 808 | 956 | 107.5 | 34.7 | 3.42E+11 | 0.15 |
| 2 | -15.0 | -18.1 | 805 | 949 | 121.5 | 34.4 | 3.86E+11 | 0.85 |

### e_id: 4784, ZetaScript: SMALL SOP100 zp, Sample name: MDA_MB_361

Measurement created: 2025-02-20 14:35:26 Report created: 2025-03-26 14:56:40 User: ZetaUser

#### Measurement specification: SOP100_ZP

##### Capture settings

| | | | |
|---|---|---|---|
| Temperature / °C | 25.0 | Laser wave length / nm | 488 |
| Dilution | 4000 | Filter wave length / nm | 0 |
| Viscosity / mPa · s | 0.9 | Sensitivity | 80 |
| Positions | 0.149 | Shutter | 90 |
| Cycle frames | 30 | Dielectric constant | 78.434 |
| cycle amount | 4 | - Uref / mV | -15.564 |
| Frame rate / 1/s | 30.303 | + Uref / mV | 16.115 |
| Experiment type | Zeta Potential | Conductivity / µS/cm | 1085.0 |
| Comment: | | | |

##### Analysis settings

| | |
|---|---|
| Concentration correction | 1 |
| Concentration calibration | 636400 |
| Henry factor | 1.5 |

#### Graphics:

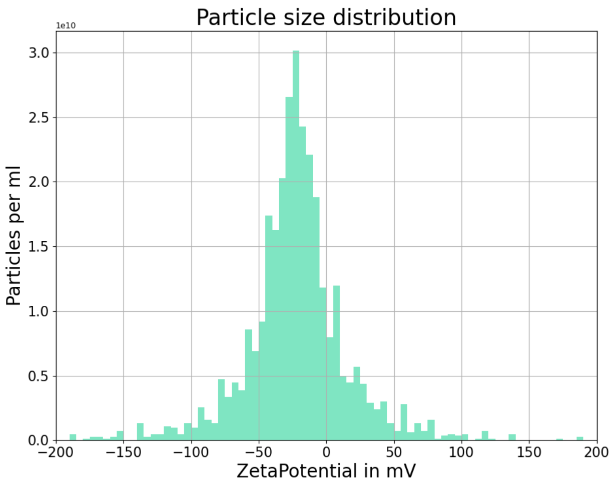


#### Results:

| | Peak /mV | Median /mV | #Zeta particles | #All particles | Avg. Num Det. | FWHM /mV | Concentration /1/mL | Position |
|---|---|---|---|---|---|---|---|---|
| | **-24.6 (± 4.9)** | **-22.1 (± 5.5)** | **2523** | **3040** | **144.1 (± 12.2)** | **40.2 (± 6.9)** | **3.67E+11 (± 3.1E+10)** | |
| 1 | -29.5 | -27.5 | 1417 | 1705 | 156.3 | 47.1 | 3.98E+11 | 0.15 |
| 2 | -19.6 | -16.6 | 1106 | 1335 | 131.9 | 33.3 | 3.36E+11 | 0.85 |

### e_id: 4784, ZetaScript: SMALL SOP100 zp, Sample name: MDA_MB_361

Measurement created: 2025-02-20 14:35:26 Report created: 2025-03-26 14:56:40 User: ZetaUser

#### Measurement specification: SOP100_ZP_2

##### Capture settings

| | | | |
|---|---|---|---|
| Temperature / °C | 25.2 | Laser wave length / nm | 488 |
| Dilution | 4000 | Filter wave length / nm | 0 |
| Viscosity / mPa · s | 0.896 | Sensitivity | 80 |
| Positions | 0.149 | Shutter | 90 |
| Cycle frames | 30 | Dielectric constant | 78.356 |
| cycle amount | 4 | - Uref / mV | -15.222 |
| Frame rate / 1/s | 30.303 | + Uref / mV | 15.676 |
| Experiment type | Zeta Potential | Conductivity / µS/cm | 1111.0 |
| Comment: | | | |

##### Analysis settings

| | |
|---|---|
| Concentration correction | 1 |
| Concentration calibration | 636400 |
| Henry factor | 1.5 |

#### Graphics:

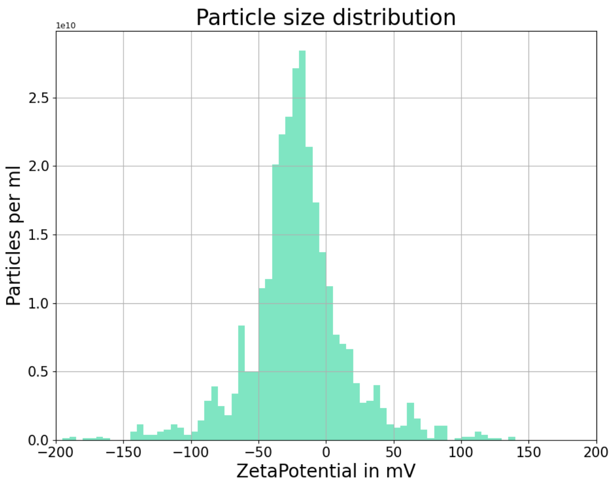


#### Results:

| | Peak /mV | Median /mV | #Zeta particles | #All particles | Avg. Num Det. | FWHM /mV | Concentration /1/mL | Position |
|---|---|---|---|---|---|---|---|---|
| | **-23.6 (± 6.2)** | **-21.5 (± 6.3)** | **2309** | **2758** | **141.3 (± 14.9)** | **36.1 (± 1.2)** | **3.60E+11 (± 3.8E+10)** | |
| 1 | -29.9 | -27.8 | 1283 | 1528 | 156.2 | 37.3 | 3.98E+11 | 0.15 |
| 2 | -17.4 | -15.3 | 1026 | 1230 | 126.4 | 34.9 | 3.22E+11 | 0.85 |

### e_id: 5027, ZetaScript: SMALL SOP100 zp, Sample name: THP1 msd calibrator 1

Measurement created: 2025-04-01 15:06:05 Report created: 2025-04-01 15:11:03 User: ZetaUser

#### Measurement specification: SOP100_ZP

##### Capture settings

| | | | |
|---|---|---|---|
| Temperature / °C | 24.6 | Laser wave length / nm | 488 |
| Dilution | 3000 | Filter wave length / nm | 0 |
| Viscosity / mPa · s | 0.908 | Sensitivity | 80 |
| Positions | 0.149 | Shutter | 90 |
| Cycle frames | 30 | Dielectric constant | 78.566 |
| cycle amount | 4 | - Uref / mV | -16.659 |
| Frame rate / 1/s | 30.303 | + Uref / mV | 16.844 |
| Experiment type | Zeta Potential | Conductivity / µS/cm | 1429.0 |
| Comment: | | | |

##### Analysis settings

| | |
|---|---|
| Concentration correction | 1 |
| Concentration calibration | 636400 |
| Henry factor | 1.5 |

#### Graphics:

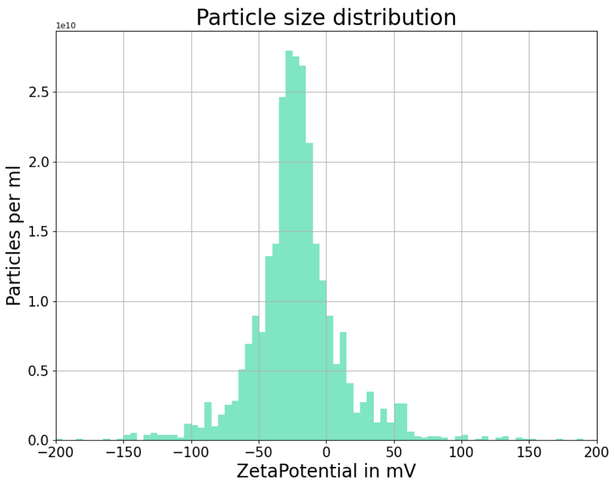


#### Results:

| | Peak /mV | Median /mV | #Zeta particles | #All particles | Avg. Num Det. | FWHM /mV | Concentration /1/mL | Position |
|---|---|---|---|---|---|---|---|---|
| | **-22.8 (± 4.1)** | **-22.9 (± 3.0)** | **3033** | **3606** | **172.3 (± 3.2)** | **38.9 (± 0.5)** | **3.29E+11 (± 6.1E+09)** | |
| 1 | -26.8 | -25.9 | 1624 | 1915 | 175.5 | 39.4 | 3.35E+11 | 0.15 |
| 2 | -18.7 | -19.9 | 1409 | 1691 | 169.1 | 38.5 | 3.23E+11 | 0.85 |

### e_id: 5027, ZetaScript: SMALL SOP100 zp, Sample name: THP1 msd calibrator 1

Measurement created: 2025-04-01 15:06:05    Report created: 2025-04-01 15:11:04    User: ZetaUser

#### Measurement specification: SOP100_ZP_2

##### Capture settings

| | | | |
|---|---|---|---|
| Temperature / °C | 24.8 | Laser wave length / nm | 488 |
| Dilution | 3000 | Filter wave length / nm | 0 |
| Viscosity / mPa · s | 0.905 | Sensitivity | 80 |
| Positions | 0.149 | Shutter | 90 |
| Cycle frames | 30 | Dielectric constant | 78.516 |
| cycle amount | 4 | - Uref / mV | -16.767 |
| Frame rate / 1/s | 30.303 | + Uref / mV | 16.946 |
| Experiment type | Zeta Potential | Conductivity / µS/cm | 1342.0 |
| Comment: | | | |

##### Analysis settings

| | |
|---|---|
| Concentration correction | 1 |
| Concentration calibration | 636400 |
| Henry factor | 1.5 |

#### Graphics:

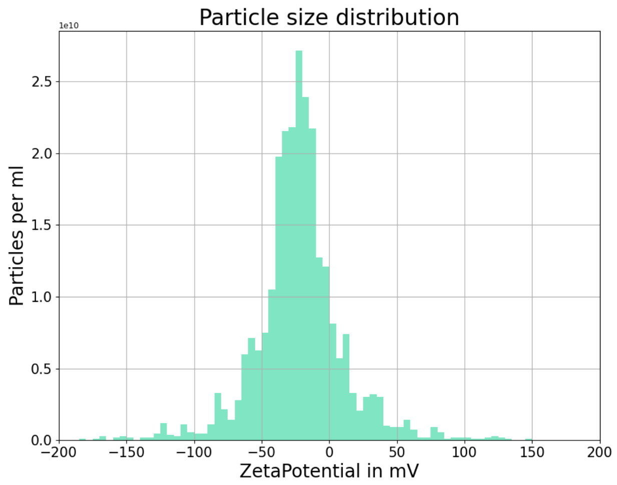


#### Results:

| | Peak /mV | Median /mV | #Zeta particles | #All particles | Avg. Num Det. | FWHM /mV | Concentration /1/mL | Position |
|---|---|---|---|---|---|---|---|---|
| | **-21.4 (± 3.2)** | **-22.6 (± 2.1)** | **2774** | **3344** | **164.1 (± 5.9)** | **35.7 (± 1.4)** | **3.13E+11 (± 1.1E+10)** | |
| 1 | -24.6 | -24.7 | 1482 | 1790 | 170.0 | 37.1 | 3.25E+11 | 0.15 |
| 2 | -18.2 | -20.5 | 1292 | 1554 | 158.1 | 34.3 | 3.02E+11 | 0.85 |
