## Supplementary Material 6 for "PHoNUPS: Open-Source Software for Standardized Analysis and Visualization of Multi-Instrument Extracellular Vesicle Measurements"

### e_id: 5099, ZetaScript: SMALL SOP100 zp FM488, Sample name: CD9-GFP alone

Measurement created: 2025-04-07 13:10:18 Report created: 2025-07-14 11:50:29 User: ZetaUser

#### Measurement specification: SOP100_ZP_2

##### Capture settings

| | | | |
|---|---|---|---|
| Temperature / °C | 25.1 | Laser wave length / nm | 488 |
| Dilution | 1000 | Filter wave length / nm | 0 |
| Viscosity / mPa · s | 0.899 | Sensitivity | 80 |
| Positions | 0.149 | Shutter | 90 |
| Cycle frames | 30 | Dielectric constant | 78.409 |
| cycle amount | 4 | - Uref / mV | -14.36 |
| Frame rate / 1/s | 30.303 | + Uref / mV | 14.583 |
| Experiment type | Zeta Potential | Conductivity / µS/cm | 1248.0 |
| Comment: | | | |

##### Analysis settings

| | |
|---|---|
| Concentration correction | 1 |
| Concentration calibration | 636400 |
| Henry factor | 1.5 |

#### Graphics:

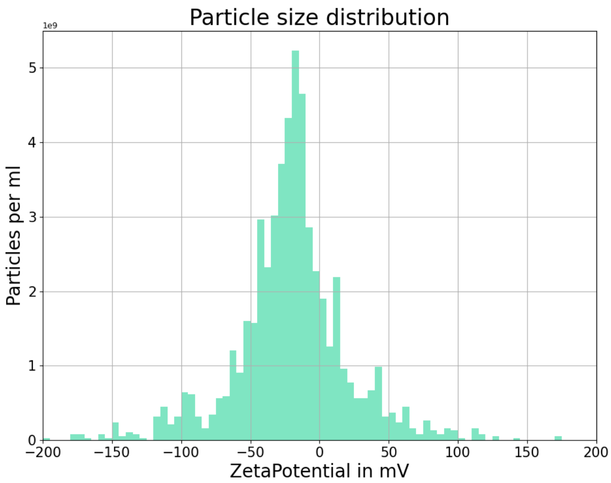


#### Results:

| | Peak /mV | Median /mV | #Zeta particles | #All particles | Avg. Num Det. | FWHM /mV | Concentration /1/mL | Position |
|---|---|---|---|---|---|---|---|---|
| | **-18.9 (± 1.6)** | **-19.9 (± 1.0)** | **2051** | **2470** | **103.6 (± 11.9)** | **48.7 (± 1.5)** | **6.60E+10 (± 7.5E+09)** | |
| 1 | -17.4 | -19.0 | 1098 | 1315 | 115.5 | 50.2 | 7.35E+10 | 0.15 |
| 2 | -20.5 | -20.9 | 953 | 1155 | 91.8 | 47.1 | 5.84E+10 | 0.85 |

### e_id: 5099, ZetaScript: SMALL SOP100 zp FM488, Sample name: CD9-GFP alone

Measurement created: 2025-04-07 13:10:18 Report created: 2025-07-14 11:50:29 User: ZetaUser

#### Measurement specification: SOP100_ZP

##### Capture settings

| | | | |
|---|---|---|---|
| Temperature / °C | 25.1 | Laser wave length / nm | 488 |
| Dilution | 1000 | Filter wave length / nm | 0 |
| Viscosity / mPa · s | 0.898 | Sensitivity | 80 |
| Positions | 0.149 | Shutter | 90 |
| Cycle frames | 30 | Dielectric constant | 78.388 |
| cycle amount | 4 | - Uref / mV | -14.349 |
| Frame rate / 1/s | 30.303 | + Uref / mV | 14.788 |
| Experiment type | Zeta Potential | Conductivity / µS/cm | 1229.0 |
| Comment: | | | |

##### Analysis settings

| | |
|---|---|
| Concentration correction | 1 |
| Concentration calibration | 636400 |
| Henry factor | 1.5 |

#### Graphics:

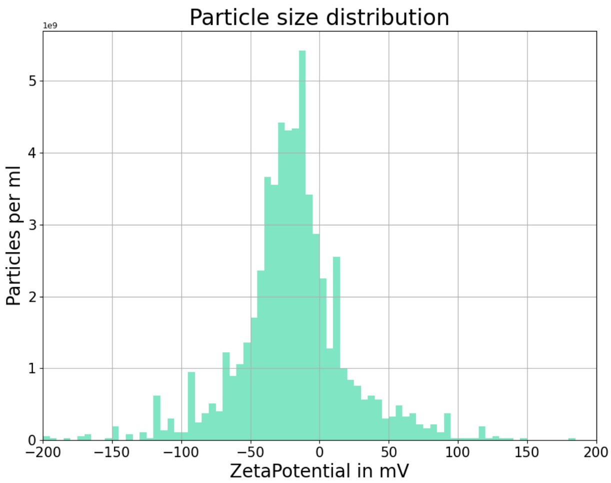


#### Results:

| | Peak /mV | Median /mV | #Zeta particles | #All particles | Avg. Num Det. | FWHM /mV | Concentration /1/mL | Position |
|---|---|---|---|---|---|---|---|---|
| | **-18.3 (± 2.0)** | **-19.5 (± 1.7)** | **2177** | **2577** | **109.8 (± 4.9)** | **50.6 (± 5.0)** | **6.99E+10 (± 3.1E+09)** | |
| 1 | -16.3 | -17.8 | 1160 | 1369 | 104.9 | 55.6 | 6.68E+10 | 0.15 |
| 2 | -20.3 | -21.2 | 1017 | 1208 | 114.7 | 45.5 | 7.30E+10 | 0.85 |

### e_id: 3606, ZetaScript: SMALL SOP100 zp, Sample name: F1_SOP100_EDTA_IN_onhe_label

Measurement created: 2024-11-06 15:07:27    Report created: 2024-11-06 16:44:06    User: ZetaUser

#### Measurement specification: SOP100_ZP

##### Capture settings

| | | | |
|---|---|---|---|
| Temperature / °C | 24.9 | Laser wave length / nm | 488 |
| Dilution | 4000 | Filter wave length / nm | 0 |
| Viscosity / mPa · s | 0.901 | Sensitivity | 80 |
| Positions | 0.149 | Shutter | 90 |
| Cycle frames | 30 | Dielectric constant | 78.452 |
| cycle amount | 4 | - Uref / mV | -17.086 |
| Frame rate / 1/s | 30.303 | + Uref / mV | 17.645 |
| Experiment type | Zeta Potential | Conductivity / µS/cm | 1339.0 |
| Comment: | | | |

##### Analysis settings

| | |
|---|---|
| Concentration correction | 1 |
| Concentration calibration | 636400 |
| Henry factor | 1.5 |

#### Graphics:

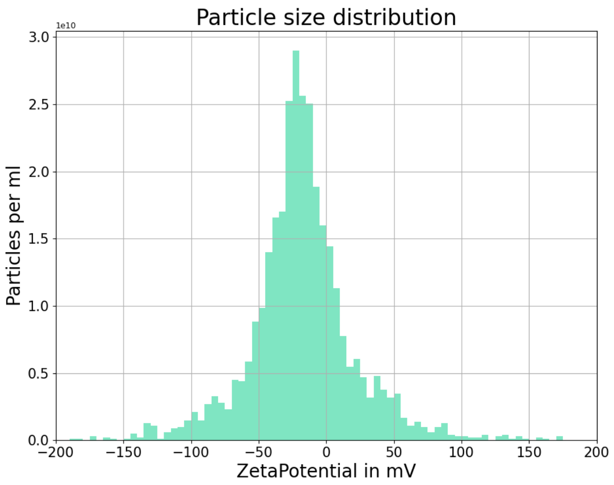


#### Results:

| | Peak /mV | Median /mV | #Zeta particles | #All particles | Avg. Num Det. | FWHM /mV | Concentration /1/mL | Position |
|---|---|---|---|---|---|---|---|---|
| | **-19.9 (± 3.1)** | **-19.3 (± 1.9)** | **3256** | **3855** | **150.4 (± 4.8)** | **46.3 (± 4.6)** | **3.83E+11 (± 1.2E+10)** | |
| 1 | -23.0 | -21.2 | 1727 | 2040 | 155.3 | 41.8 | 3.95E+11 | 0.15 |
| 2 | -16.9 | -17.3 | 1529 | 1815 | 145.6 | 50.9 | 3.71E+11 | 0.85 |

### e_id: 3606, ZetaScript: SMALL SOP100 zp, Sample name: F1_SOP100_EDTA_IN_onhe_label

Measurement created: 2024-11-06 15:07:27    Report created: 2024-11-06 16:44:06    User: ZetaUser

#### Measurement specification: SOP100_ZP_2

##### Capture settings

| | | | |
|---|---|---|---|
| Temperature / °C | 25.1 | Laser wave length / nm | 488 |
| Dilution | 4000 | Filter wave length / nm | 0 |
| Viscosity / mPa · s | 0.898 | Sensitivity | 80 |
| Positions | 0.149 | Shutter | 90 |
| Cycle frames | 30 | Dielectric constant | 78.395 |
| cycle amount | 4 | - Uref / mV | -16.919 |
| Frame rate / 1/s | 30.303 | + Uref / mV | 17.35 |
| Experiment type | Zeta Potential | Conductivity / µS/cm | 1387.0 |
| Comment: | | | |

##### Analysis settings

| | |
|---|---|
| Concentration correction | 1 |
| Concentration calibration | 636400 |
| Henry factor | 1.5 |

#### Graphics:

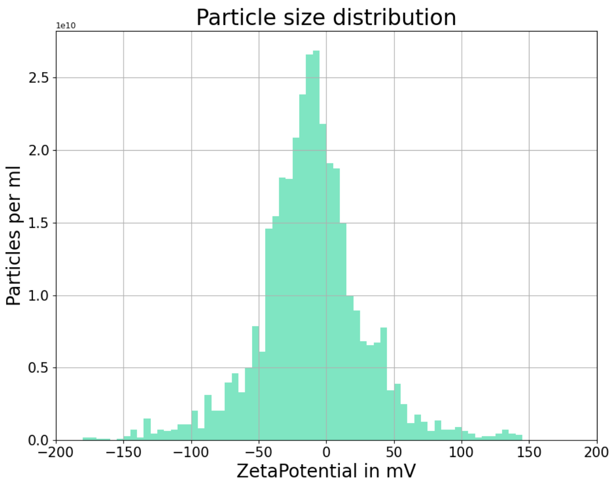


#### Results:

| | Peak /mV | Median /mV | #Zeta particles | #All particles | Avg. Num Det. | FWHM /mV | Concentration /1/mL | Position |
|---|---|---|---|---|---|---|---|---|
| | **-13.5 (± 7.8)** | **-13.7 (± 8.1)** | **3876** | **4717** | **171.2 (± 22.0)** | **45.4 (± 0.3)** | **4.36E+11 (± 5.6E+10)** | |
| 1 | -21.4 | -21.8 | 1571 | 1967 | 149.2 | 45.1 | 3.80E+11 | 0.15 |
| 2 | -5.7 | -5.6 | 2305 | 2750 | 193.2 | 45.8 | 4.92E+11 | 0.85 |

### e_id: 3620, ZetaScript: SMALL SOP100 zp, Sample name: F9_SOP100_EDTA_IN_onhe_label

Measurement created: 2024-11-06 16:29:32    Report created: 2024-11-06 16:44:21    User: ZetaUser

#### Measurement specification: SOP100_ZP

##### Capture settings

| | | | |
|---|---|---|---|
| Temperature / °C | 25.0 | Laser wave length / nm | 488 |
| Dilution | 1000 | Filter wave length / nm | 0 |
| Viscosity / mPa · s | 0.9 | Sensitivity | 80 |
| Positions | 0.149 | Shutter | 90 |
| Cycle frames | 30 | Dielectric constant | 78.43 |
| cycle amount | 4 | - Uref / mV | -17.392 |
| Frame rate / 1/s | 30.303 | + Uref / mV | 18.081 |
| Experiment type | Zeta Potential | Conductivity / µS/cm | 1326.0 |
| Comment: | | | |

##### Analysis settings

| | |
|---|---|
| Concentration correction | 1 |
| Concentration calibration | 636400 |
| Henry factor | 1.5 |

#### Graphics:

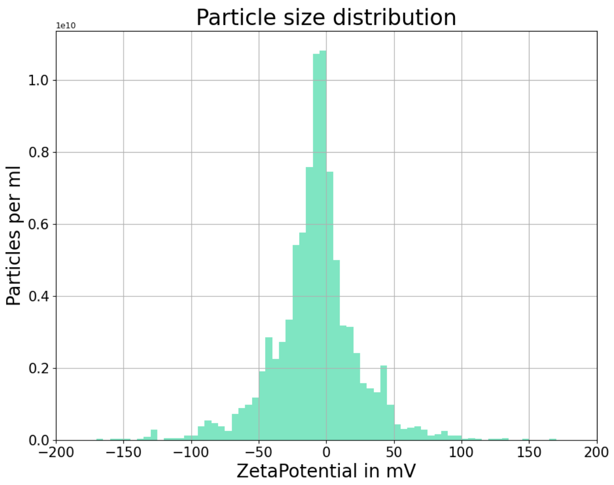


#### Results:

| | Peak /mV | Median /mV | #Zeta particles | #All particles | Avg. Num Det. | FWHM /mV | Concentration /1/mL | Position |
|---|---|---|---|---|---|---|---|---|
| | **-5.2 (± 1.8)** | **-6.9 (± 2.2)** | **2884** | **3456** | **173.0 (± 26.6)** | **31.7 (± 4.3)** | **1.10E+11 (± 1.7E+10)** | |
| 1 | -7.0 | -9.1 | 1345 | 1632 | 146.4 | 36.0 | 9.32E+10 | 0.15 |
| 2 | -3.4 | -4.7 | 1539 | 1824 | 199.6 | 27.4 | 1.27E+11 | 0.85 |

### e_id: 3620, ZetaScript: SMALL SOP100 zp, Sample name: F9_SOP100_EDTA_IN_onhe_label

Measurement created: 2024-11-06 16:29:32 Report created: 2024-11-06 16:44:21 User: ZetaUser

#### Measurement specification: SOP100_ZP_2

##### Capture settings

| | | | |
|---|---|---|---|
| Temperature / °C | 25.0 | Laser wave length / nm | 488 |
| Dilution | 1000 | Filter wave length / nm | 0 |
| Viscosity / mPa · s | 0.9 | Sensitivity | 80 |
| Positions | 0.149 | Shutter | 90 |
| Cycle frames | 30 | Dielectric constant | 78.434 |
| cycle amount | 4 | - Uref / mV | -17.44 |
| Frame rate / 1/s | 30.303 | + Uref / mV | 18.077 |
| Experiment type | Zeta Potential | Conductivity / µS/cm | 1323.0 |
| Comment: | | | |

##### Analysis settings

| | |
|---|---|
| Concentration correction | 1 |
| Concentration calibration | 636400 |
| Henry factor | 1.5 |

#### Graphics:

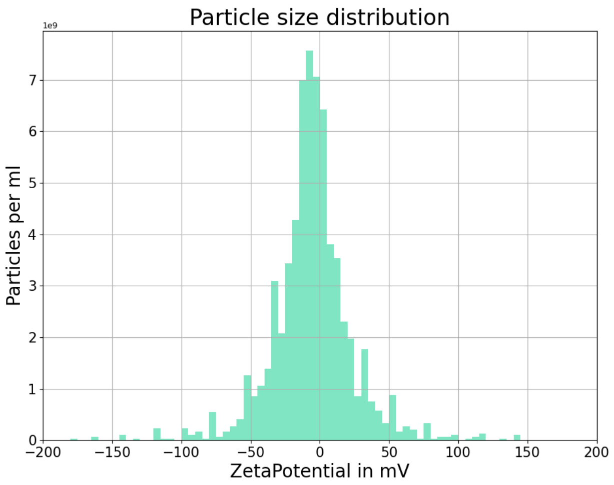


#### Results:

| | Peak /mV | Median /mV | #Zeta particles | #All particles | Avg. Num Det. | FWHM /mV | Concentration /1/mL | Position |
|---|---|---|---|---|---|---|---|---|
| | **-2.9 (± 2.9)** | **-3.0 (± 3.0)** | **1958** | **5338** | **284.9 (± 50.5)** | **15.3 (± 15.3)** | **1.81E+11 (± 3.2E+10)** | |
| 1 | -5.7 | -6.0 | 1958 | 2313 | 234.4 | 30.7 | 1.49E+11 | 0.15 |
| 2 | 0.0 | 0.0 | 0 | 3025 | 335.4 | 0.0 | 2.13E+11 | 0.85 |
