## Supplementary figures and images for "PHoNUPS: Open-Source Software for Standardized Analysis and Visualization of Multi-Instrument Extracellular Vesicle Measurements"

### Supplementary Material 7

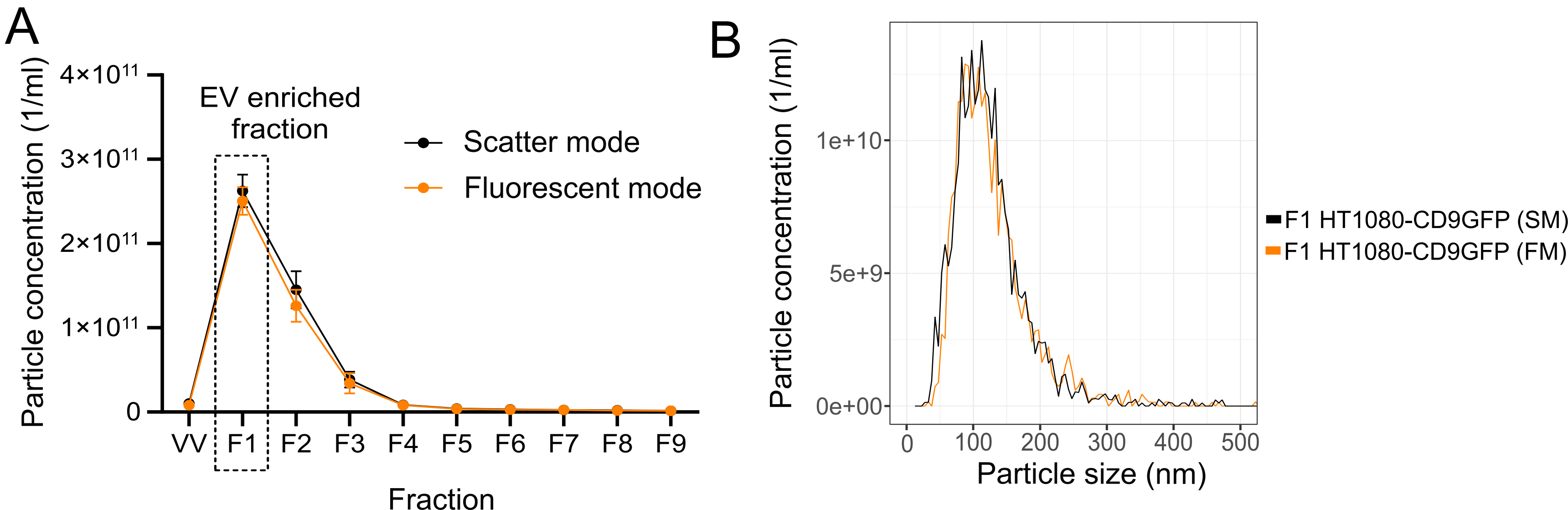
